# Spatial N-glycan rearrangement on α_5_β_1_ integrin nucleates galectin-3 oligomers to determine endocytic fate

**DOI:** 10.1101/2023.10.27.564026

**Authors:** Massiullah Shafaq-Zadah, Estelle Dransart, Christian Wunder, Valérie Chambon, Cesar A. Valades-Cruz, Ludovic Leconte, Nirod Kumar Sarangi, Jack Robinson, Siau-Kun Bai, Raju Regmi, Aurélie Di Cicco, Agnès Hovasse, Richard Bartels, Ulf J. Nilsson, Sarah Cianférani-Sanglier, Hakon Leffler, Tia E. Keyes, Daniel Lévy, Stefan Raunser, Daniel Roderer, Ludger Johannes

## Abstract

Membrane glycoproteins frequently adopt different conformations when altering between active and inactive states. Here, we discover a molecular switch that exploits dynamic spatial rearrangements of N-glycans during such conformational transitions to control protein function. For the conformationally switchable cell adhesion glycoprotein α_5_β_1_ integrin, we find that only the bent-closed state arranges N-glycans to nucleate the formation of up to tetrameric oligomers of the glycan-binding protein galectin-3. We propose a structural model of how these galectin-3 oligomers are assembled and how they clamp the bent-closed state to prime it for endocytic uptake and subsequent retrograde trafficking to the Golgi for polarized distribution in cells. Our findings highlight an unexpectedly dynamic regulation of the glycan landscape at the cell surface to achieve oligomerization of galectin-3. Galectin-3 oligomers are thereby identified as decoders of defined spatial patterns of N-glycans and as functional extracellular interactors of specifically the bent-closed conformational state of α_5_β_1_ integrin and possibly other family members.

## Introduction

Integrins are heterodimeric glycoproteins with key functions in the adhesion of cells to extracellular matrix ligands such as fibronectin (Bridgewater et al., 2012; Marsico et al., 2018; Moreno-Layseca et al., 2019). Integrins exist in a continuum of conformations between bent-closed non-ligand-bound (also termed inactive) and extended-open ligand-bound (also termed active) states (Li et al., 2017). N-glycosylation positions were suggested to affect the equilibrium between these two conformers (Hang et al., 2017), but the underlying mechanisms remain largely unexplored.

Glycosylation affects the functions of integrins (Gu and Taniguchi, 2004; Hang et al., 2017; Hou et al., 2016; Isaji et al., 2009; Isaji et al., 2006; Lagana et al., 2006; Pan and Song, 2010; Pretzlaff et al., 2000; Seales et al., 2005; Boscher and Nabi, 2013; Goetz et al., 2008). Distal N-glycosylation sites within the β-propeller domain of α_5_ and at the I-like domain of β_1_ integrins are required for heterodimerization and efficient cell surface expression (Hou et al., 2016; Isaji et al., 2006). Membrane proximal N-glycosylation sites are needed for the biological activities of integrins, including interactions with other membrane proteins such as the epidermal growth factor receptor (Caswell et al., 2008; Hou et al., 2016).

Both clathrin-dependent and independent endocytic mechanisms have been documented for integrin internalization (Almeida-Souza et al., 2018; Chao and Kunz, 2009; Ezratty et al., 2009; Ezratty et al., 2005; Furtak et al., 2001; Lakshminarayan et al., 2014; Moreno-Layseca et al., 2021; Shi and Sottile, 2011; Sottile and Chandler, 2005). While the clathrin pathway remains the best characterized endocytic route (Kaksonen and Roux, 2018; Mettlen et al., 2018; Robinson, 2015), the mechanisms by which membranes are bent and cargoes selected in the absence of the clathrin coat remain under investigation (Boucrot et al., 2015; Caldieri et al., 2017; Lakshminarayan et al., 2014; Moreno-Layseca et al., 2021; Sathe et al., 2018). Glycans on proteins and lipids as well as glycan binding proteins of the galectin family are at center stage in one of the proposed models, termed glycolipid-lectin (GL-Lect) driven endocytosis (Johannes et al., 2015; Johannes et al., 2016; Kenworthy et al., 2021; Pezeshkian et al., 2021; Renard and Boucrot, 2021). Notably galectin-3 (Gal3) has been analyzed in depth (Furtak et al., 2001; Lakshminarayan et al., 2014). For building endocytic pits, Gal3 recognizes N-glycans on proteins (including integrins), and oligomerizes. Only oligomerized Gal3 acquires the capacity to bind to glycosphingolipids (GSLs) to induce the formation of tubular endocytic pits from which clathrin-independent endocytic carriers (CLICs) emerge (Lakshminarayan et al., 2014). Of note, Gal3 has also been described to form lattice assemblies at the plasma membrane (Nabi et al., 2015). The interplay between GL- Lect and lattice processes controls the cell surface dynamics of glycoproteins, including integrins, in an intertwined manner (Dennis, 2015; Mathew and Donaldson, 2019).

The oligomerization capacity of Gal3 has been shown to be important for its function in endocytosis (Lakshminarayan et al., 2014), but shapes and assembly mechanisms of Gal3 oligomers remained controversial (Ahmad et al., 2004; Lepur et al., 2012). Here, we have found that a conformational state-specific spatial arrangement of N-glycans on α_5_β_1_ integrin sets its capacity to nucleate Gal3 oligomers that range up to ring-shaped tetramers. This determines the integrin’s endocytic fate and ensuing intracellular compartmentalization. We have termed this glycan pattern recognition mechanism the conformational glycoswitch, which may also apply to other members of the integrin family. The conformational glycoswitch positions glycosylation as a highly dynamic regulator of integrin function at the cell surface.

## Results

### Retrograde trafficking of inactive bent-closed α_5_β_1_ integrin depends on Gal3

We have previously described that only the inactive bent-closed non-ligand-bound conformational state of α_5_β_1_ integrin (Figure 1A), and not the active extended-open ligand-bound state, follows the retrograde trafficking route from the plasma membrane to the Golgi apparatus from where the protein then undergoes polarized secretion for its dynamic localization to the leading edge of migrating cells (Shafaq-Zadah et al., 2016). We discovered this dichotomic behavior using conformational state-specific antibodies, i.e., mAb13 against the β_1_ chain of inactive bent-closed α_5_β_1_ integrin (Mould et al., 1995), and 9EG7 against the β_1_ chain of active ligand-bound α_5_β_1_ integrin (Lenter et al., 1993). These were modified with benzylguanine (BG) to be captured in live cell antibody uptake experiments by a GFP-labeled SNAP-tag fusion protein that was localized to the Golgi apparatus (Figure 1B) (Johannes and Shafaq-Zadah, 2013; Shafaq-Zadah et al., 2016).

**Figure 1.**
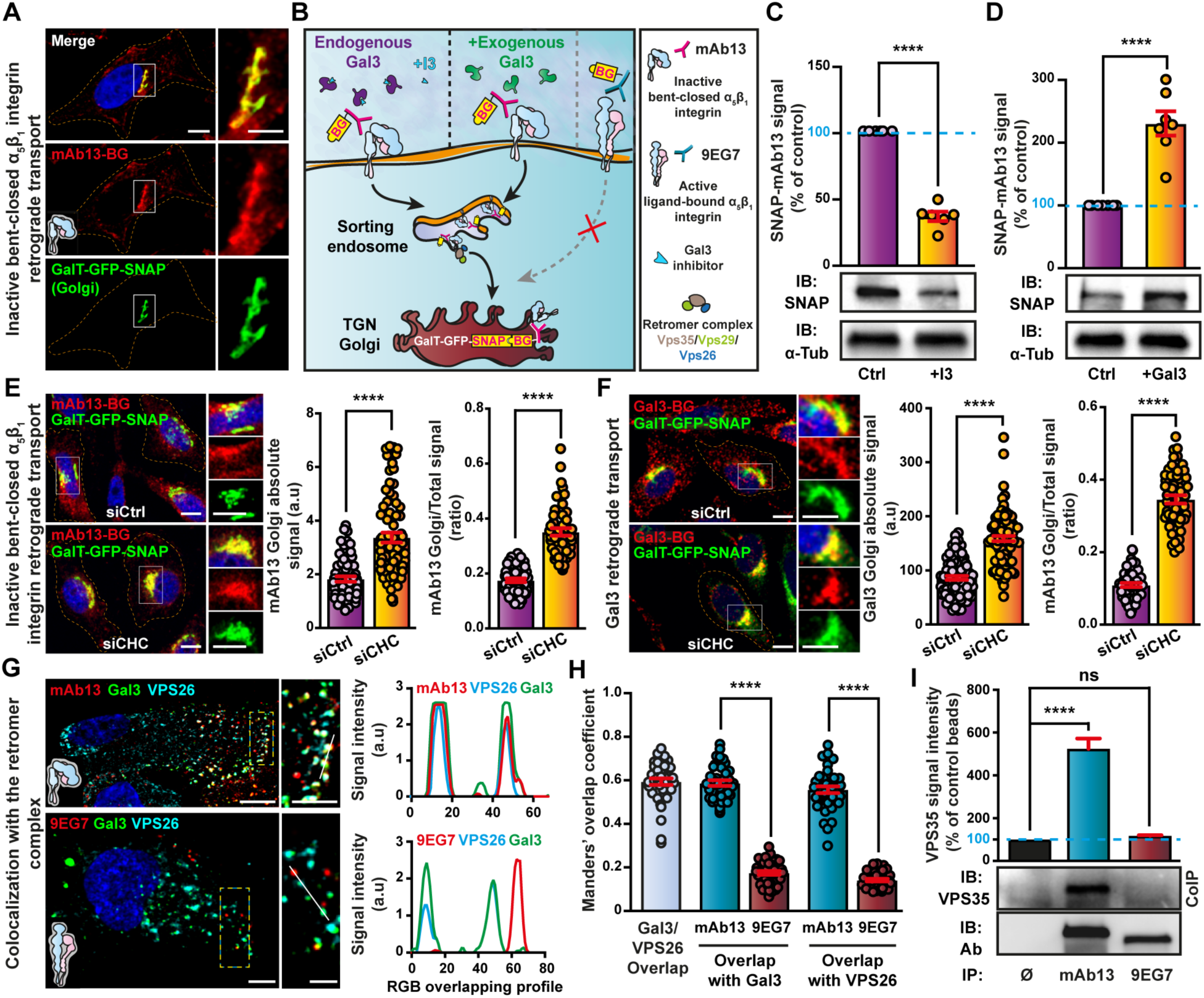
Gal3 drives the clathrin-independent retrograde trafficking of the inactive bent- closed α_5_β_1_ integrin. (**A**) Retrograde trafficking of inactive bent-closed α_5_β_1_ integrin (mAb13) to the Golgi compartment. Continuous incubation of GalT-GFP-SNAP expressing HeLa cells for 1 h at 37 °C with Cy3 and benzylguanine (BG)-labeled mAb13 (mAb13-BG). Note that mAb13-BG colocalized with Golgi-localized GalT-GFP-SNAP. Scale bars = 10 µm. Nuclei in blue. (B) Regulation of mAb13 retrograde trafficking by Gal3. Schematics of experimental set up and conditions. Note that the active ligand-bound conformation (9EG7) does not undergo retrograde trafficking, as previously reported (Shafaq-Zadah et al., 2016). (**C**,**D**) Use of GFP-trap to pull down stably expressed Golgi-localized GalT-GFP-SNAP from HeLa cells that had continuously been incubated for 3 h at 37 °C with mAb13-BG in the presence or absence of the Gal3 inhibitor I3 (10 µM) (C), or exogenous Gal3 (200 nM) (D). mAb13-BG that reached the Golgi was quantified by immunoblotting (IB: SNAP) for the corresponding SNAP-mAb13 conjugates. α-tubulin was used for normalization. 6-7 independent experiments, means ± SEM, unpaired t-test; ****P < 0.0001. (**E**) As in (A), either in control condition (siCtrl) or after inhibition of clathrin expression (siCHC). Intensities of absolute mAb13 fluorescent signals in the Golgi were quantified. 3 independent experiments, means ± SEM, unpaired t-test; ****P < 0.0001. Note that mAb13-BG colocalized with Golgi-resident GalT-GFP-SNAP, and that inhibition of clathrin induced an increased accumulation of mAb13 in the Golgi. The fractions of mAb13 fluorescent signal reaching the Golgi over the total signal were also quantified and reported as ratios. 3 independent experiments, means ± SEM, unpaired t-test; ****P < 0.0001. (**F**) Gal3 trafficking to the Golgi compartment. Cy3 and BG-labeled Gal3 (200 nM) was incubated for 45 min at 37 °C with GalT-GFP-SNAP expressing HeLa cells, either in control (siCtrl) or clathrin depletion (siCHC) conditions. Absolute intensities of Gal3 fluorescence signals in the Golgi were quantified. 3 independent experiments, means ± SEM, unpaired t-test; ****P < 0.0001. Note that Gal3-BG colocalized with Golgi-resident GalT-GFP-SNAP, and that inhibition of clathrin induced an increased accumulation of Gal3 in the Golgi, as observed for mAb13-BG. The fractions of Gal3 fluorescent signal reaching the Golgi over the total signal were also quantified and reported as ratios. 3 independent experiments, means ± SEM, unpaired t-test; ****P < 0.0001. Scale bars = 10 µm. Nuclei in blue. (**G**) α_5_β_1_ integrin colocalization with retromer. Sequential binding of Gal3 (200 nM) and mAb13 or 9EG7 antibodies to RPE-1 cells at 4 °C, incubation for 15 min at 37 °C, followed by labeling of fixed cells for Vps26. Right panels show intensities along the white lines in the zooms to the left. Scale bars = 10 µm, and 5 µm for zoomed insets. Nuclei in blue. (**H**) Quantification of overlaps from experiments as in (G). 3 independent experiments, means ± SEM, unpaired t-test; ****P < 0.0001. (**I**) α_5_β_1_ integrin interaction with retromer. mAb13 and 9EG7 were continuously incubated for 20 min at 37 °C with RPE-1 cells and then immunoprecipitated (IP). Cells without antibodies (Ø) served as controls. Immunoblotting for Vps35 (IB: VPS35) and antibodies (IB: Ab) revealed that only inactive α_5_β_1_ integrin (mAb13) co-immunoprecipitated Vps35. 3 independent experiments, means ± SEM, one-way ANOVA; ns = P > 0.05, ****P < 0.0001.

Here, we found that retrograde transport of inactive bent-closed α_5_β_1_ integrin (mAb13) from the plasma membrane to the Golgi apparatus was strongly decreased by incubation in the presence of the GB0149-03 compound, a membrane impermeable inhibitor of Gal3 that we termed I3 in the current study (Salameh et al., 2010; Stegmayr et al., 2019) (Figure 1C), and stimulated by exogenous Gal3 (Figure 1D). In clear contrast, the depletion of clathrin heavy chain (CHC) not only did not inhibit retrograde trafficking of mAb13, but even increased it (Figures 1E and S1A). Since BG-tagged mAb13 that underwent retrograde trafficking was irreversibly captured in the Golgi apparatus by the GalT-GFP-SNAP-tag fusion protein (Figures 1A and 1B), the observed phenotype was not due to a possible effect of CHC depletion on Golgi exit. In contrast to retrograde transport, the overall endocytosis of inactive bent-closed α_5_β_1_ integrin (mAb13) was slightly reduced upon CHC depletion (Figure S1B) (Kyumurkov et al., 2023). The endocytosis of the clathrin pathway marker transferrin, which served as a positive control (Hopkins et al., 1985), was strongly inhibited in CHC depleted cells (Figure S1C), and not affected by Gal3 inhibition (I3) (Figure S1D).

A similar effect of CHC depletion was found for Gal3 itself. We here show for the first time that Gal3 undergoes retrograde trafficking (Figure S1E and S1F), and that accumulation in the Golgi apparatus of exogenously added BG-tagged Gal3 was increased upon CHC depletion (Figure 1F), while its endocytosis was partly inhibited under these conditions (Figure S1G). Taken together, our findings strongly suggest that only the fractions of inactive bent-closed α_5_β_1_ integrin and of Gal3 that enter cells by clathrin-independent endocytosis gain access to the retrograde route.

Internalized inactive bent-closed α_5_β_1_ integrin (mAb13) and Gal3 both strongly colocalized on endosomal membranes with the Vps26 component of the retromer complex (Figure 1G and 1H), a retrograde sorting machinery (Seaman, 2004). The colocalization of internalized active ligand- bound α_5_β_1_ integrin (9EG7) with Gal3 and Vps26 was much lower (Figure 1G and 1H), as expected from the inability of this conformer to undergo retrograde trafficking (Shafaq-Zadah et al., 2016). Furthermore, inactive bent-closed endosomal α_5_β_1_ integrin (mAb13) co-immunoprecipitated the retromer subunit Vps35, while active ligand-bound α_5_β_1_ integrin (9EG7) did not (Figure 1I).

These data demonstrate that retrograde trafficking of inactive bent-closed α_5_β_1_ integrin is dependent on Gal3 and not on clathrin, and indicate that Gal3 and this α_5_β_1_ integrin conformer may traffic together to the Golgi apparatus.

### Specifically inactive bent-closed α_5_β_1_ integrin is internalized by GL-Lect driven endocytosis

We investigated whether the select Gal3 dependency for retrograde trafficking of inactive bent- closed α_5_β_1_ integrin was already set at the plasma membrane. This conformer efficiently interacted with Gal3 in coimmunoprecipitation experiments (Figure 2A, mAb13), and colocalized with Gal3 by immunofluorescence at the plasma membrane (mAb13 for β_1_ integrin, Figure 2B top; mAb16 for α_5_ integrin, Figure S2A left) and in endosomes (Figure 2C, top). Both at plasma membrane (Figure S2B) and in endosomes (Figure S2C), colocalization was much reduced with Gal3ΔNter, a Gal3 mutant whose N-terminal oligomerization domain was deleted (Nieminen et al., 2007). Gal3ΔNter is deficient in all functions that have been attributed to Gal3, including GL-Lect driven endocytosis (Lakshminarayan et al., 2014). This finding indicates that Gal3 oligomerization is important for its efficient co-distribution with the inactive bent-closed α_5_β_1_ integrin conformer.

**Figure 2.**
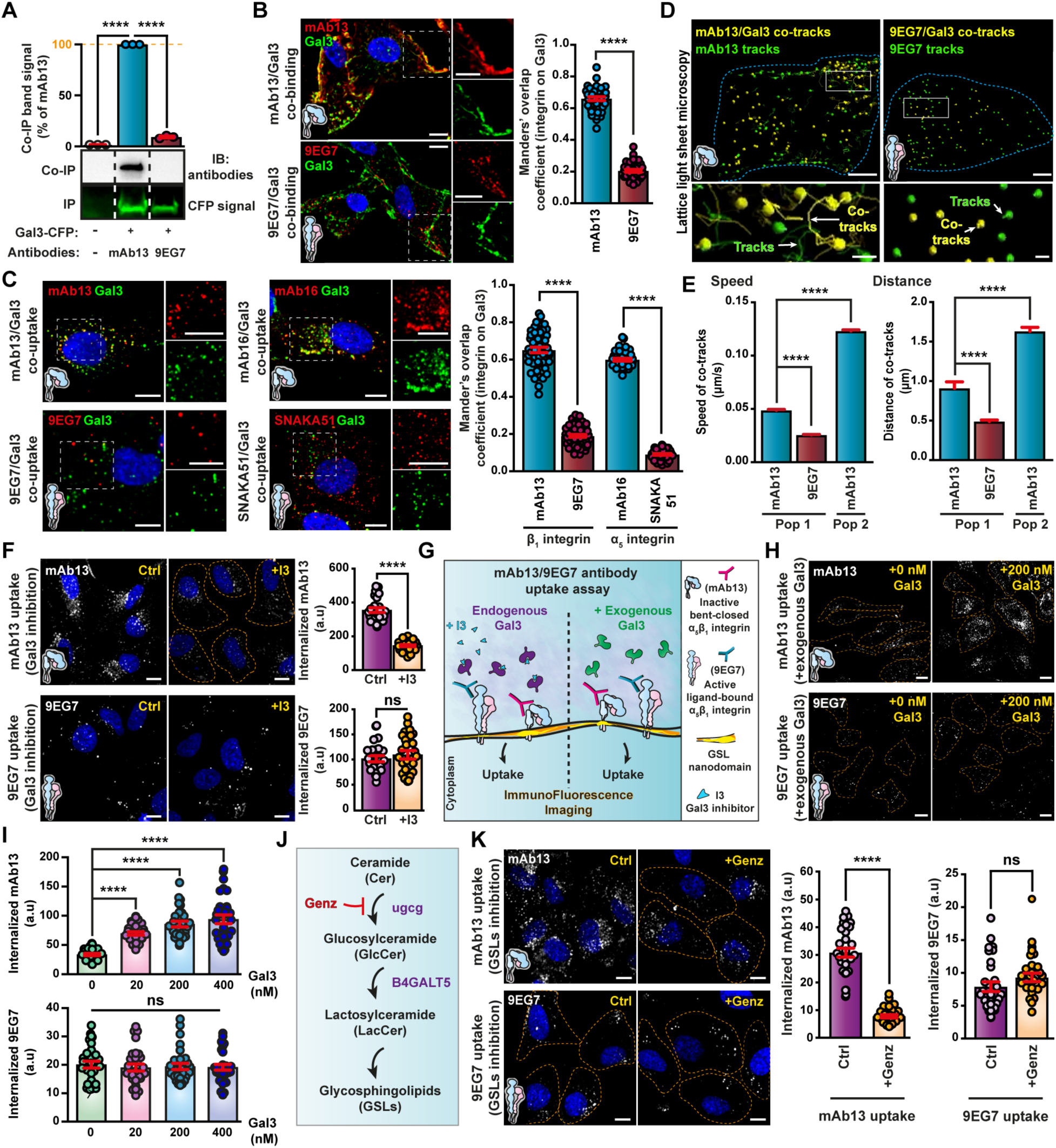
Gal3 specifically interacts with inactive bent-closed α_5_β_1_ integrin, and promotes its GL-Lect driven endocytosis. (**A**) α_5_β_1_ integrin interaction with Gal3. Immunoprecipitation (IP) of Gal3-CFP transiently expressed in RPE-1 cells and immunoblotting for co-immunoprecipitated (Co-IP) mAb13 and 9EG7 (IB: antibodies) that had been surface-bound (4 °C) to these cells. CFP fluorescent signal was used for normalization. 3 independent experiments, means ± SEM, one- way ANOVA; ****P < 0.0001. (**B**) Colocalization of exogenous Gal3 (200 nM) with mAb13 (inactive bent-closed) or 9EG7 (active ligand-bound) antibodies after incubation at 4 °C with RPE- 1 cells. Scale bars = 10 µm. Nuclei in blue. 3 independent experiments, means ± SEM, unpaired t-test; ****P < 0.0001. (**C**) Co-uptake of Gal3 with α_5_β_1_ integrin. RPE-1 cells were sequentially incubated at 4 °C with Gal3 (200 nM) and either mAb13 or 9EG7, or the equally conformational state-specific α_5_ integrin antibodies mAb16 (inactive) or SNAKA51 (active). Cells were then shifted for 10 min to 37 °C. The overlap of fluorescence signals was quantified. 3 independent experiments, means ± SEM, unpaired t-test; ****P < 0.0001. Scale bars = 10 µm. Nuclei in blue. (**D**) Lattice light sheet microscopy. Co-tracking of Gal3 with mAb13 (7 cells) or 9EG7 (10 cells) for 3 min at 27 °C. Co-tracks in yellow, Gal3-negative mAb13 or 9EG7 tracks in green. Scale bars = 5 µm. Inset: 3D rendering of cargo (mAb13 and 9EG7 antibodies) detection (spheres) and tracks (lines) from representative cells. Scale bars = 1 µm. (**E**) Left: quantification of Gal3/mAb13 (blue bars) or Gal3/9EG7 co-tracks (red bars) velocity and delimitation of two speed-populations: Pop 1 co-tracks with speeds inferior to 0.08 µm/s, and Pop 2 co-tracks with speeds above 0.08 µm/s. Right: quantification of co-tracking distance within the two speed-delimited populations. Note that both, distance, and speed of mAb13/Gal3 co-tracks were higher than 9EG7/Gal3 co-tracks, and that co-tracks with speeds above 0.08 µm/s (Pop 2) were observed exclusively for mAb13/Gal3 co-tracks. At least 289 co-tracks were analyzed per condition. Means ± SEM, one-way ANOVA; ****P < 0.0001. (**F**) Effect of Gal3 inhibitor I3 on α_5_β_1_ integrin endocytosis. Quantification by confocal microscopy of internalized mAb13 and 9EG7 after continuous incubation for 10 min at 37 °C with RPE-1 cells pre-treated with I3 (+ I3, 10 µM). 3 independent experiments, means ± SEM, unpaired t-test; ns = P > 0.05, ****P < 0.0001. Scale bars = 10 µm. Nuclei in blue. (**G**) GL- Lect driven endocytosis of inactive bent-closed α_5_β_1_ integrin. Schematics of experimental procedures. (**H**) Effect of exogenous Gal3 on α_5_β_1_ integrin endocytosis. After mAb13 or 9EG7 binding to RPE-1 cells (dashed lines) at 4 °C, internalization was measured after 10 min incubation at 37 °C in the presence of the indicated concentrations of exogenous Gal3. (**I**) Quantification from experiments as in (H) of fluorescence signals of internalized mAb13 or 9EG7. 3 independent experiments, means ± SEM, one-way ANOVA; ****P < 0.0001. Scale bars = 10 µm. (**J**) Pathway of GSL biosynthesis and effect of Genz. (**K**) Effect of GSL depletion on α_5_β_1_ integrin endocytosis. Quantification of internalized mAb13 and 9EG7 antibodies after continuous incubation for 5 min at 37 °C with RPE-1 cells in conditions of GSL depletion (+ Genz). 3 independent experiments, means ± SEM, unpaired t-test; ns = P > 0.05, ****P < 0.0001. Scale bars = 10 µm. Nuclei in blue.

For active ligand-bound α_5_β_1_ integrin, interaction (Figure 2A, 9EG7) and colocalization (9EG7 for the β_1_ chain, Figure 2B bottom; SNAKA51 for the α_5_ chain, Figure S2A right) with Gal3 were very low. This dichotomic colocalization behavior with Gal3 between inactive bent-closed (mAb13) and active ligand-bound (9EG7) α_5_β_1_ integrin was even more pronounced at the leading edge of migrating cells (Figure S2D for plasma membrane, and Figure S2E for endosomes). Time-resolved imaging by lattice light sheet microscopy furthermore demonstrated the exclusive association of inactive bent-closed α_5_β_1_ integrin (mAb13) and Gal3 onto dynamic co-tracks, when compared to the active ligand-bound state (9EG7) (population 2 (Pop2), Figures 2D, 2E and S2F; Movies S1 and S2).

In accordance with its strong interaction and colocalization with Gal3, the endocytic uptake of inactive bent-closed α_5_β_1_ integrin (mAb13) turned out to be GL-Lect driven, since it was (i) inhibited by the Gal3 inhibitor I3 (Figure 2F, top); (ii) stimulated by exogenously added wild-type Gal3 (Figures 2G, 2H, and 2I, top), but not by Gal3ΔNter (Figure S3A); and (iii) inhibited by the depletion of another component of the GL-Lect machinery, GSLs, using the glucosylceramide synthase inhibitor Genz-123346 (Zhao et al., 2007) (Figures 2J and 2K, top), much as seen for Gal3 itself (Figure S3B). In contrast, all these treatments did not affect the uptake of active ligand-bound 9EG7-labeled α_5_β_1_ integrin (Figures 2F, 2H, 2I and 2K, bottom), nor that of transferrin (Figure S3C). The dichotomic behavior between inactive bent-closed and active ligand-bound α_5_β_1_ integrin was thereby also observed for GL-Lect driven endocytosis.

One of the hallmarks of GL-Lect driven endocytosis is the formation of tubular often crescent-shaped clathrin-independent endocytic carriers (CLICs) (Lakshminarayan et al., 2014). To analyze the structures via which inactive bent-closed and active ligand-bound α_5_β_1_ integrins are internalized, respectively mAb13 and 9EG7 were coupled to horseradish peroxidase (HRP) and incubated with cells at 37 °C for a time course between 6-9 min. Endocytic carriers were then analyzed by electron microscopy (EM) for the presence of these markers. Inactive bent-closed α_5_β_1_ integrin (mAb13) was found extensively in short tubular CLICs, and much less in vesicles (Figure 3A), which corroborates the conclusion that a substantial fraction of this conformer enters cells by GL-Lect driven endocytosis. We also found that the dynein inhibitor ciliobrevin (CBD), which interferes with friction-driven scission that specifically operates during clathrin-independent endocytosis (Renard et al., 2015; Simunovic et al., 2017), strongly reduced the endocytosis of inactive bent-closed α_5_β_1_ integrin (mAb13), an effect that was further potentiated when exogenous Gal3 was added to cells (Figure 3B). Ciliobrevin also strongly inhibited the uptake of Gal3 (Figure S3D), and only weakly that of transferrin (Figure S3E).

**Figure 3.**
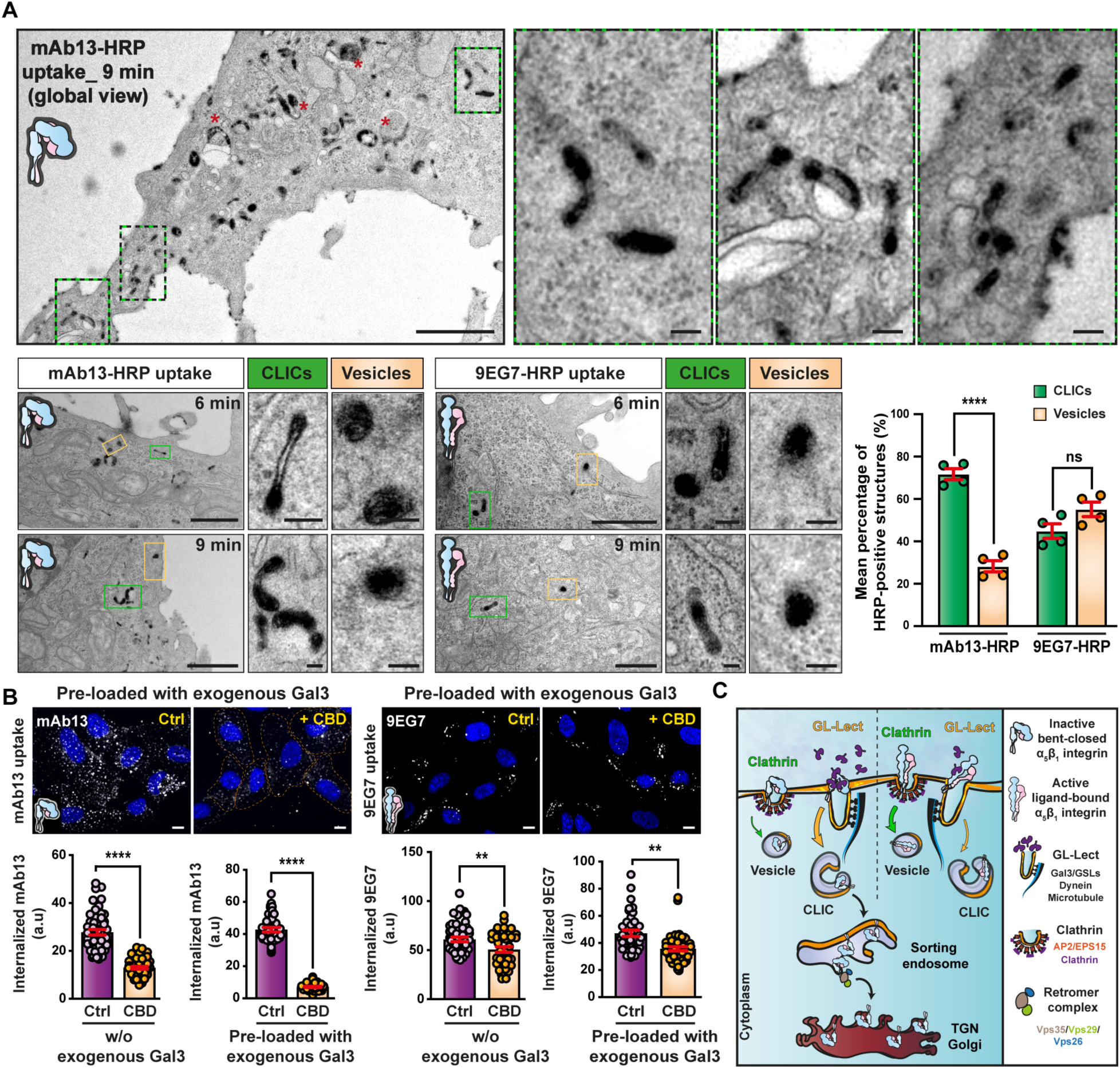
Endocytosis of the inactive bent-closed α_5_β_1_ integrin via CLICs. (**A**) Inactive bent- closed α_5_β_1_ integrin accumulates in clathrin-independent carriers (CLICs). Top, electron microscopy micrographs of HeLa cells that were incubated continuously with HRP-coupled mAb13 (mAb13-HRP) for 9 min at 37 °C; scale bar = 1 µm. Green/black dashed insets: Zooms of areas illustrating accumulation of mAb13-HRP in typical crescent-shaped CLIC structures. Red stars indicate larger, likely endosomal compartments. Scale bars = 100 nm. Bottom, same experiments as above, for both mAb13-HRP or 9EG7-HRP antibodies (6 or 9 min uptake). Quantification of endocytic structures: 4 independent experiments, means ± SEM, unpaired t-test; ns = P > 0.05, ****P < 0.0001. Scale bars = 1 µm. Green insets: Zooms of CLICs. Orange insets: Zooms of endocytic vesicles. Scale bars = 100 nm. (**B**) Effect of ciliobrevin D (CBD) on α_5_β_1_ integrin endocytosis. Quantification by confocal microscopy of mAb13 or 9EG7 uptake after incubation for 10 min at 37 °C with RPE-1 cells incubated or not with CBD (+ CBD). Cells were preloaded or not with exogenous Gal3 (200 nM) for 30 min at 4 °C. Note that preloading of Gal3 induced a stronger inhibition of mAb13 uptake upon CBD treatment, while 9EG7 endocytosis was not sensitive to Gal3 preloading. 3 independent experiments, means ± SEM, unpaired t-test; **P < 0.002, ****P < 0.0001. Scale bars = 10 µm. Nuclei in blue. (**C**) Schematic of key outcomes. Note the dichotomic effect of Gal3 on binding, GL-Lect driven endocytosis, and retrograde transport of specifically the inactive bent-closed conformational state of α_5_β_1_ integrin.

The presence of some inactive bent-closed α_5_β_1_ integrin (mAb13) in vesicles is consistent with a previous report on the contribution of clathrin-dependent endocytosis to the cellular uptake of this conformer close to focal adhesions (Kyumurkov et al., 2023), and with our findings that depletion of CHC (Figure S1B) or expression of a dominant negative mutant of the clathrin accessory protein Eps15 (EPS15DN) (Benmerah et al., 1998) had measurable effects on the cellular uptake of inactive bent-closed α_5_β_1_ integrin (mAb13, Figure S3F, top) and Gal3 (Figure S3G, top).

Active ligand-bound α_5_β_1_ integrin (9EG7) was almost equally distributed between CLICs and vesicles (Figure 3A). The vesicular fraction likely reflects clathrin-dependent uptake, based on: (i) previous findings (Almeida-Souza et al., 2018; Chao and Kunz, 2009; Ezratty et al., 2009); (ii) their size (120 nm ±29 nm, n=95) (Kirchhausen et al., 2014); (iii) the observation of a substantial inhibition of this conformer (9EG7; Figure S3F, bottom) and transferrin uptake (Figure S3G, bottom) in EPS15DN expressing cells; (iv) coimmunoprecipitation with CHC (Figure S3H); and (v) the pronounced colocalization of endocytic tracks of active ligand-bound α_5_β_1_ integrin (9EG7) with the clathrin pathway component adaptor protein 2 (AP2), as monitored under unperturbed conditions (Kural et al., 2015; Renard et al., 2020) using lattice light sheet microscopy (Figures S3I-L and Movies S3-S6).

The presence of a fraction of active ligand-bound α_5_β_1_ integrin (9EG7) in CLICs is consistent with a previous report (Moreno-Layseca et al., 2021) and suggests that different mechanisms exist for the generation of CLICs.

The cell-based experiments from above on interaction, codistribution at the plasma membrane and in endosomes, molecular mechanisms of endocytic uptake and retrograde trafficking to the Golgi apparatus led to the discovery of a dichotomic imprint of Gal3 onto specifically the inactive bent-closed conformational state of α_5_β_1_ integrin (Figure 3C). To understand the molecular basis for this dichotomic behavior, we set out to combine biophysical and structural techniques applied to different biochemical reconstitution systems.

### Specifically inactive bent-closed α_5_β_1_ integrin nucleates the formation of Gal3 oligomers

α_5_β_1_ integrin was purified from rat liver in detergent micelles (Dransart et al., 2022) (Figures 4A and S4A) and reconstituted into microcavity array-suspended lipid bilayers (MSLBs) in which α_5_β_1_ integrin heterodimers are free to diffuse laterally (Figure 4B), as described previously (Sarangi# et al., 2022). In this reductionist model of the plasma membrane, electrochemical impedance spectroscopy allowed the label-free measurement of Gal3 binding to α_5_β_1_ integrin through changes in membrane resistance and capacitance. At low nanomolar concentrations of Gal3, the capacitance signal ΔQ increased for membranes containing inactive bent-closed α_5_β_1_ integrin and plateaued at Gal3 concentrations above 5 nM (Figure 4C, circles), pointing to highly efficient Gal3 binding. In contrast, the capacitance signal remained unchanged on incubation with Gal3 concentrations as high as 37 nM on membranes containing α_5_β_1_ integrin that had been activated with Mn^2+^ and the minimal extracellular matrix mimicking cRGD peptide (Bazzoni et al., 1995) (Figure 4C, triangles).

**Figure 4.**
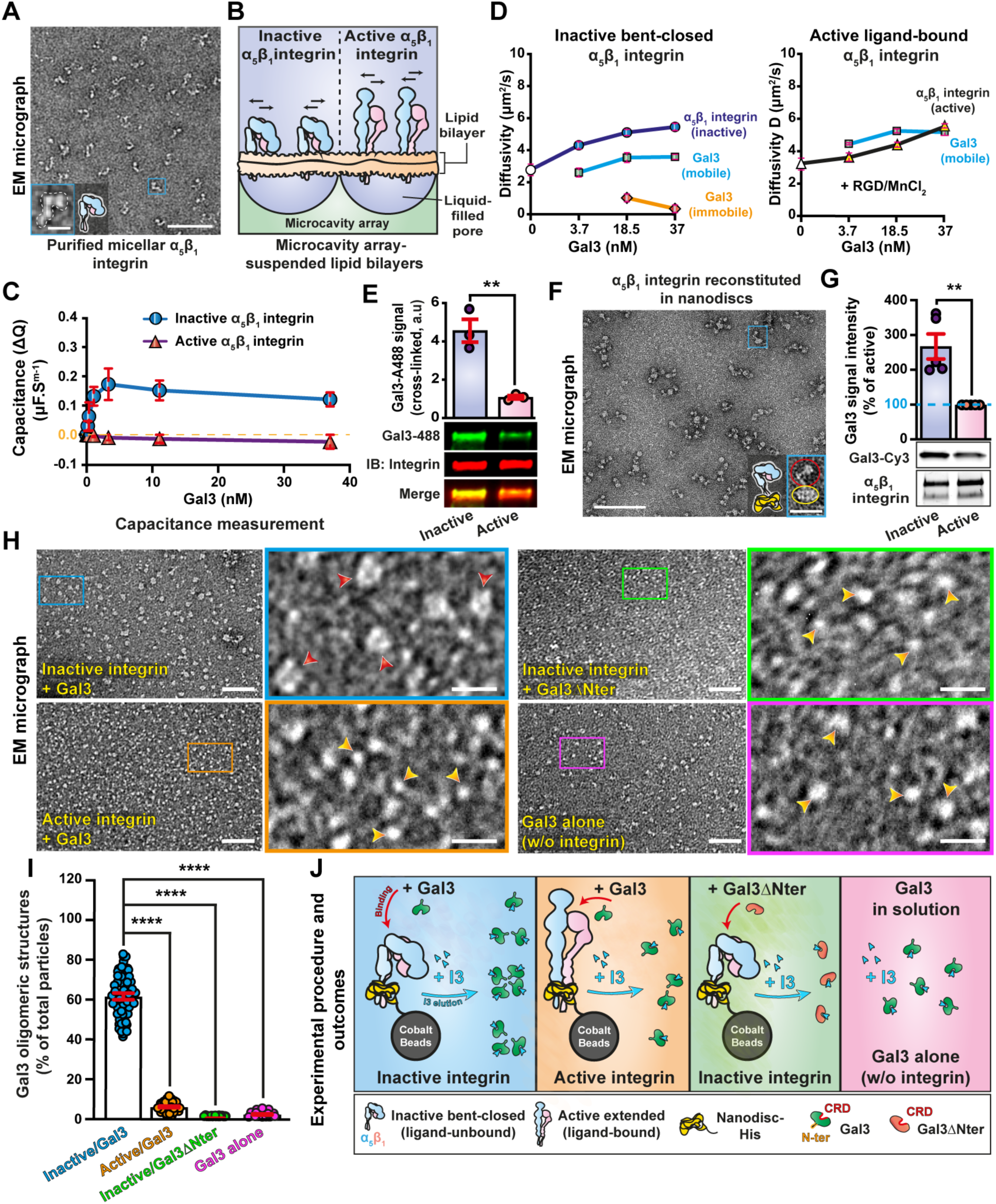
**Only inactive bent-closed α_5_β_1_ integrin nucleates the formation of Gal3 oligomers.** (**A**) Negative stain EM micrograph of micelle-embedded α_5_β_1_ integrin. Scale bar = 100 nm. Inset: Zoomed view of one α_5_β_1_ integrin. Scale bar = 10 nm. (**B**) Schematics of inactive bent-closed and active extended ligand-bound α_5_β_1_ integrin embedded in a microcavity array-supported lipidic bilayer (MSLB). Arrows indicate the possibility for integrins to diffuse laterally. (**C**) Capacitance measurements as a function of Gal3 concentrations on MSLBs containing α_5_β_1_ integrin in the indicated conformational states. Note that capacitance increase, indicative of membrane thickness decrease, only occurs upon incubation of Gal3 with the inactive bent-closed α_5_β_1_ integrin. 3 independent experiments, means ± SD. (**D**) Diffusivity measurements by FLIM. Micellar α_5_β_1_ integrin was reconstituted in microcavity array-suspended lipid bilayers, and incubated with increasing concentrations of Gal3. On the inactive bent-closed inactive conformational state of α_5_β_1_ integrin, the occurrence of 2 diffusional behaviors was observed for Gal3 with the appearance of a slowly diffusion population at high Gal3 concentrations. We interpret this finding as an indication for the transformation of an initial Gal3 organization (possibly Gal3 oligomers) on the inactive bent-closed inactive integrin (fast diffusion) into some higher order lateral organization (possibly lattices, slow diffusion). In contrast, this 2-phase diffusional behavior of Gal3 was not observed on the active extended ligand-bound conformational state of α_5_β_1_ integrin. White circle, inactive integrin alone; white triangle, active integrin alone. (**E**) Pulldown experiments that reveal the preferred interaction of Gal3 (Gal3-A488, 200 nM) with inactive micelle-embedded α_5_β_1_ integrin, as compared to *in vitro* activated α_5_β_1_ integrin. Immunoblotting (IB) documents that the same amounts of integrin were pulled down in both conditions. 3 independent experiments, means ± SEM, unpaired t-test; **P < 0.002. (**F**) EM micrograph of nanodisc-embedded α_5_β_1_ integrin (yellow and red circles in zoom, respectively). Scale bar = 100 nm, and 20 nm for zoomed inset. (**G**) Pulldown of nanodisc-embedded α_5_β_1_ integrin showing preferred interaction of Gal3-Cy3 (200 nM) with the inactive bent-closed conformational state. 5 independent experiments, means ± SEM, unpaired t-test; **P < 0.002. (**H**) EM micrographs of Gal3 that was incubated at 4 µM with nanodisc-embedded inactive or *in vitro* activated α_5_β_1_ integrin, and then eluted with I3. Note the massive presence of larger and defined particle with a central lumen (Gal3 oligomers, red arrowheads) only in the eluate from inactive α_5_β_1_ integrin. These Gal3 oligomers were neither observed with Gal3ΔNter, even when eluted from inactive α_5_β_1_ integrin, nor for Gal3 in solution that were largely unorganized and spherical (orange arrowheads). Scale bars = 40 nm, and 10 nm for zoomed insets. (**I**) Oligomers were visually quantified. Inactive/Gal3: 50 EM fields with 15,707 total particles; active/Gal3: 44 EM fields with 21,096 total particles; inactive/ Gal3ΔNter: 31 EM fields with 9,888 total particles; Gal3 alone: 34 EM-fields with 24,478 total particles. Means ± SEM, one-way ANOVA; ****P < 0.0001. (**J**) Schematics of experimental procedures and outcomes.

This preferred binding of Gal3 to inactive bent-closed α_5_β_1_ integrin reconstituted in MSLBs was confirmed by fluorescence lifetime imaging microscopy (FLIM) (Figure S4B). Interestingly, co- diffusivity measurements at high Gal3 concentrations revealed a biphasic behavior of the protein in the presence of inactive bent-closed α_5_β_1_ integrin (Figure 4D). This indicated the existence of both a dynamic Gal3 binding process (mobile fraction), likely on individual α_5_β_1_ integrin heterodimers, and the formation of an immobile fraction that may have resulted from lateral cross- linking between several α_5_β_1_ integrins (lattices) (Figure 4D, left). Of note, for the inactive bent-closed conformer, two α_5_β_1_ integrin-Gal3 velocity populations had also been measured by lattice light sheet microscopy on cells (Figures 2D, 2E and S2F). In contrast, such biphasic behavior was not observed with active ligand-bound α_5_β_1_ integrin (Figure 4D, right). These findings establish that also in a minimal model of the plasma membrane, inactive bent-closed α_5_β_1_ integrin is the preferred binding partner for Gal3.

As in the microcavity array suspended lipid bilayer system, purified α_5_β_1_ integrin in micelles (Figure 4A), in nanodiscs (Dransart et al., 2022) (Figure 4F), or peptidiscs (Figure S4C and S4D) was switchable from the inactive bent-closed to the extended ligand-bound conformation upon incubation with Mn^2+^ and cRGD peptide (Figure S4D). Importantly, Gal3 again interacted to a significantly greater extent with the inactive bent-closed α_5_β_1_ integrin, as revealed by cross-linking (for micelles, Figure 4E) or pulldown (for nanodiscs, Figure 4G) assays.

These experiments in different model membrane systems establish the preferred dichotomic interaction of Gal3 with inactive bent-closed α_5_β_1_ integrin as an intrinsic property of the system. Next, we further dissected the structural basis for this finding.

Since this interaction was found to be glycan-dependent (Figures S4E and S4F), we eluted Gal3 molecules bound to nanodisc-embedded α_5_β_1_ integrin using the I3 compound and analyzed the eluted species immediately via negative stain EM. Strikingly, while the Gal3 molecules that were added to α_5_β_1_ integrin were initially monomeric in solution, at least 50% had assembled into ring-shaped Gal3 oligomers on the inactive bent-closed conformer (Figure 4H), as quantified both by visual (Figure 4I) and automated (Figure S4G) approaches. In contrast, the vast majority of Gal3 eluted as undefined and small spherical particles (likely monomeric) on active ligand-bound α_5_β_1_ integrin (Figures 4H, 4I and S4G). Eluted Gal3 oligomers from integrins disassembled over time into monomers (Figure S4H), demonstrating that oligomerization was reversible. As for Gal3 that had not been in contact with α_5_β_1_ integrin, the Gal3ΔNter mutant was also mostly monomeric, even when the latter was eluted from the inactive bent-closed α_5_β_1_ integrin conformer (Figures 4H and 4I). This correlated with the mutant’s inefficient binding to the integrin (Figure S4I).

These experiments led to the discovery of regular ring-shaped Gal3 oligomers. In line with the dichotomy theme from the cell-based and the *in vitro* interaction experiments, these oligomers were nucleated only upon Gal3 binding to the inactive bent-closed conformational state of α_5_β_1_ integrin, and their nucleation required Gal3’s N-terminal oligomerization domain (Figure 4J).

### Gal3 oligomers are nucleated on the plasma membrane

We then investigated whether these Gal3 oligomers also formed within the complex environment of the plasma membrane, where GSLs as well as many different glycosylated proteins, including α_5_β_1_ integrin and other integrins are present. For this, lactose-washed RPE-1 cells were first incubated on ice with exogenous Gal3, and then with I3 to elute Gal3 species (Figure 5A), which were then analyzed by negative stain EM (Figure 5B). Remarkably, ring-shaped oligomer structures were again abundantly detected (Figure 5B), resembling those eluted from nanodisc-reconstituted inactive bent-closed α_5_β_1_ integrin (Figure 4H) both in shape and size.

**Figure 5.**
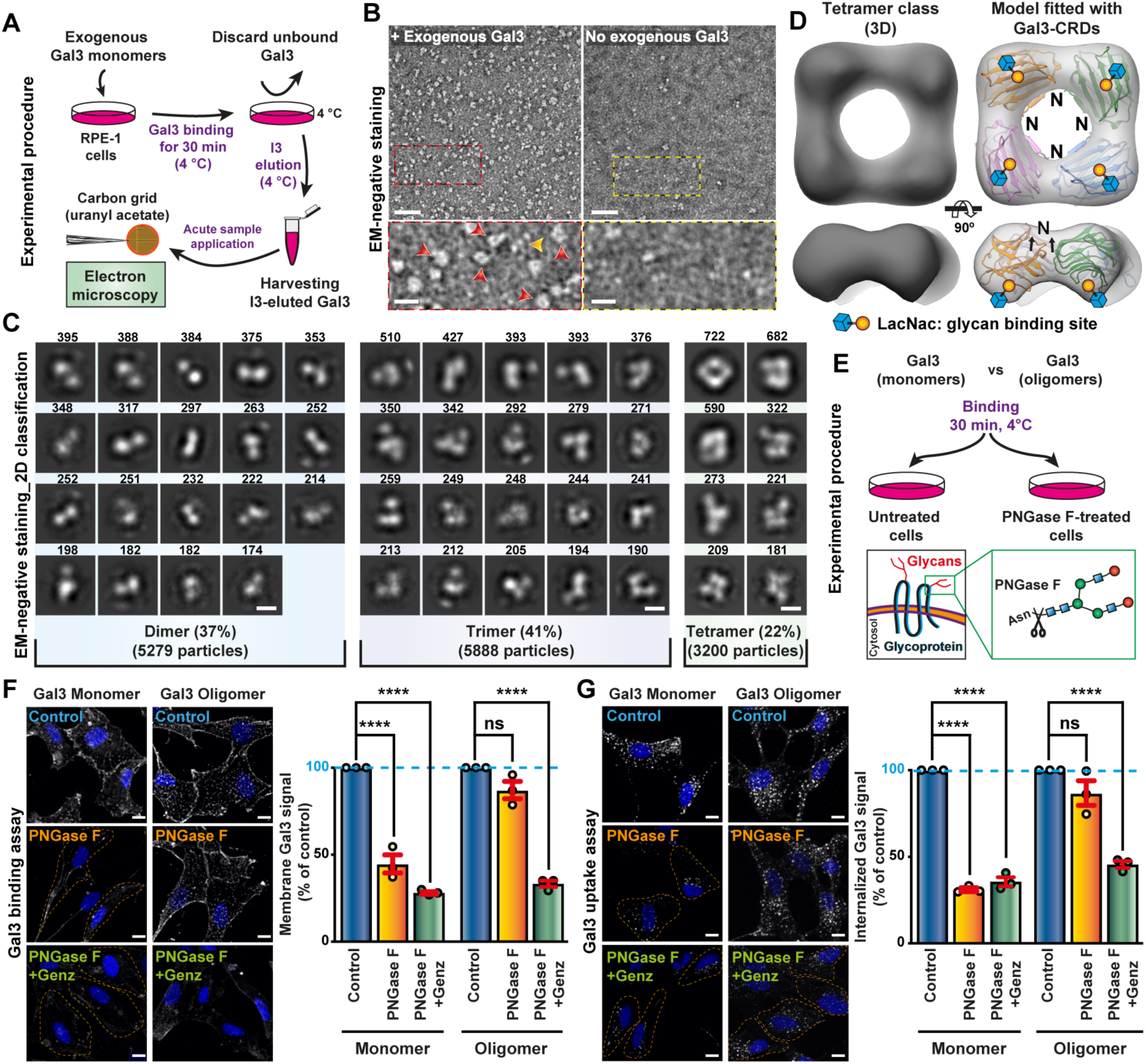
Structural and functional characterization of Gal3 oligomers assembled on cellular membranes. (**A**) Schematics for the analysis of exogenous Gal3 oligomers that are assembled on cell membrane (Gal3, 2 µM) and subsequently eluted with I3 (10 µM). (**B**) EM micrographs of Gal3 eluted from RPE-1 cells as described in (A). Red and orange arrowheads indicate oligomeric and likely monomeric Gal3, respectively. Scale bars = 40 nm, and 10 nm for the zoomed insets. (**C**) 2D class averages of negative stain EM data as in (B). Particle numbers in dimer, trimer, and tetramer class averages are indicated. The three largest classes showed annular tetramers. Scale bar = 5 nm. (**D**) 3D reconstruction of a Gal3 oligomer, originating from annular tetramer 2D classes, in which four Gal3 CRDs (PDB 1KJL) were fitted. The models are placed such that the non-resolved N-terminal domains of each individual Gal3 point towards the lumen of the ring, compatible with an oligomerization pocket, and the glycan binding pockets face outwards. (**E**) Schematics for use of PNGase F to assess the role of N-glycans in monomeric versus oligomeric Gal3 binding to cells. (**F**) Binding of Gal3 monomers versus Gal3 oligomers onto RPE-1 cells that have been treated with PNGase F, alone as described in (E), or in combination with Genz (GSL depletion). Gal3 oligomers applied here had been eluted from RPE-1 cells as in (A,B). Note that only oligomeric Gal3 still efficiently bound to PNGase F-treated cells, and that this binding was significantly decreased upon GSL depletion (Genz). Results from 3 independent experiments, means ± SEM, one-way ANOVA; ns = P > 0.05, ****P < 0.0001. (**G**) Uptake of Gal3 monomers versus oligomers into RPE-1 cells. Continuous incubation for 2 min at 37 °C in the same conditions as in (F). For Gal3 oligomers, uptake was affected only on PNGase F-treated cells that were also GSL depleted (Genz). 3 independent experiments, means ± SEM, one-way ANOVA; ns = P > 0.05, ****P < 0.0001.

By negative stain EM 2D classification of these cell derived Gal3 oligomers, we discovered that out of 14,367 particles, 5279 (37%) resembled dimers in size and shape, 5888 (41%) trimers, and 3200 (22%) tetramers (Figure 5C). Of all 2D classes, the ones depicting annular tetramers were the largest (722, 682, and 590 particles, Figure 5C). A 3D model generated from particles in the tetramer 2D classes produced a ring-shaped structure with a large central lumen (Figure 5D). Size and shape of the tetramer density map allowed to fit four Gal3 carbohydrate recognition domains (CRDs) as rigid bodies. The most consistent fit positioned them such that the N-termini of the CRDs are oriented towards the central lumen of the tetramer where the non-resolved N-terminal domains may interact, and the glycan binding sites are facing to the exterior (Figure 5D). Although other orientations cannot be ruled out due to the limited resolution of negative stain EM, the proposed CRD arrangement is compatible with two key functional properties of Gal3. First, oligomerization is dependent on the N-terminal domain (Figures 4H and 4I) and probably involves biomolecular condensate formation (Zhao et al., 2021). Second, simultaneous interaction with several spatially separated galactoses on protein N-glycans and/or GSLs is favored. Such relative orientation of glycan binding pockets towards the membrane surface has also been predicted by molecular dynamics simulations (Lete et al., 2022). In contrast, a tetramer model obtained by x-ray crystallography with N-terminally truncated versions of Gal3 assembled under non-natural conditions appears too compact to fit the EM density (Figure S4J), and orients the glycan binding pockets inwards, which is incompatible with simultaneous interaction with cargo and GSL glycans (Flores-Ibarra et al., 2018). Our data thereby represents the first 3D model of Gal3 oligomers that were made from full-length protein and nucleated on natural substrates on membranes.

The functional properties of these Gal3 oligomers were assessed on cells after treatment with PNGase F to remove surface N-glycans (Figure 5E). For monomeric Gal3, binding (Figure 5F) and internalization (Figure 5G) were significantly impaired on PNGase F-treated cells, whereas preformed Gal3 oligomers performed almost indiscriminately under control and PNGase F conditions (Figures 5F and 5G). We reasoned that while monomeric Gal3 strictly needs protein glycans for initial recruitment to the cell surface, as indeed described before (Lakshminarayan et al., 2014), pre-oligomerized Gal3 may bypass this step to functionally interact with GSLs. In agreement with this hypothesis, binding onto and internalization of preformed Gal3 oligomers into PNGase F-treated cells were significantly decreased when the cells were also GSL depleted (Figures 5F and 5G).

These findings document that Gal3 oligomers indeed have key fundamental properties as expected from the GL-Lect model, i.e., the capacity to functionally interact not only with N-glycans on glycoproteins, but also with glycans of GSLs.

### Gal3 clamps the inactive bent-closed conformational state of α_5_β_1_ integrin

To explore the molecular mechanism underlying the conformational state-specific dichotomic Gal3 oligomerization process, we set out to reveal the assembly state of Gal3 directly on inactive bent-closed α_5_β_1_ integrins. For single molecule photobleaching experiments, α_5_β_1_ integrins in the bent-closed conformation were individually inserted into biotinylated peptidiscs, desialylated to increase affinity for Gal3 (Krzeminski et al., 2011; Zhuo et al., 2008), and then complexed with fluorescently labeled Gal3 (Figure 6A). This preparation, which was devoid of visible aggregates (Figure S4K), was immobilized on neutravidin-coated substrates (Figures 6A and S4L). Up to 4 photobleaching steps could be monitored (Figures 6B and 6C), suggestive of Gal3 oligomers up to tetramers. In the absence of biotin from the peptidiscs (Figure 6C and S4L, no biotin) or of integrin altogether (Figure S4M, Gal3 alone), we observed primarily one-step photobleaching, indicative of non-specific Gal3 monomer binding to the neutravidin surface.

**Figure 6.**
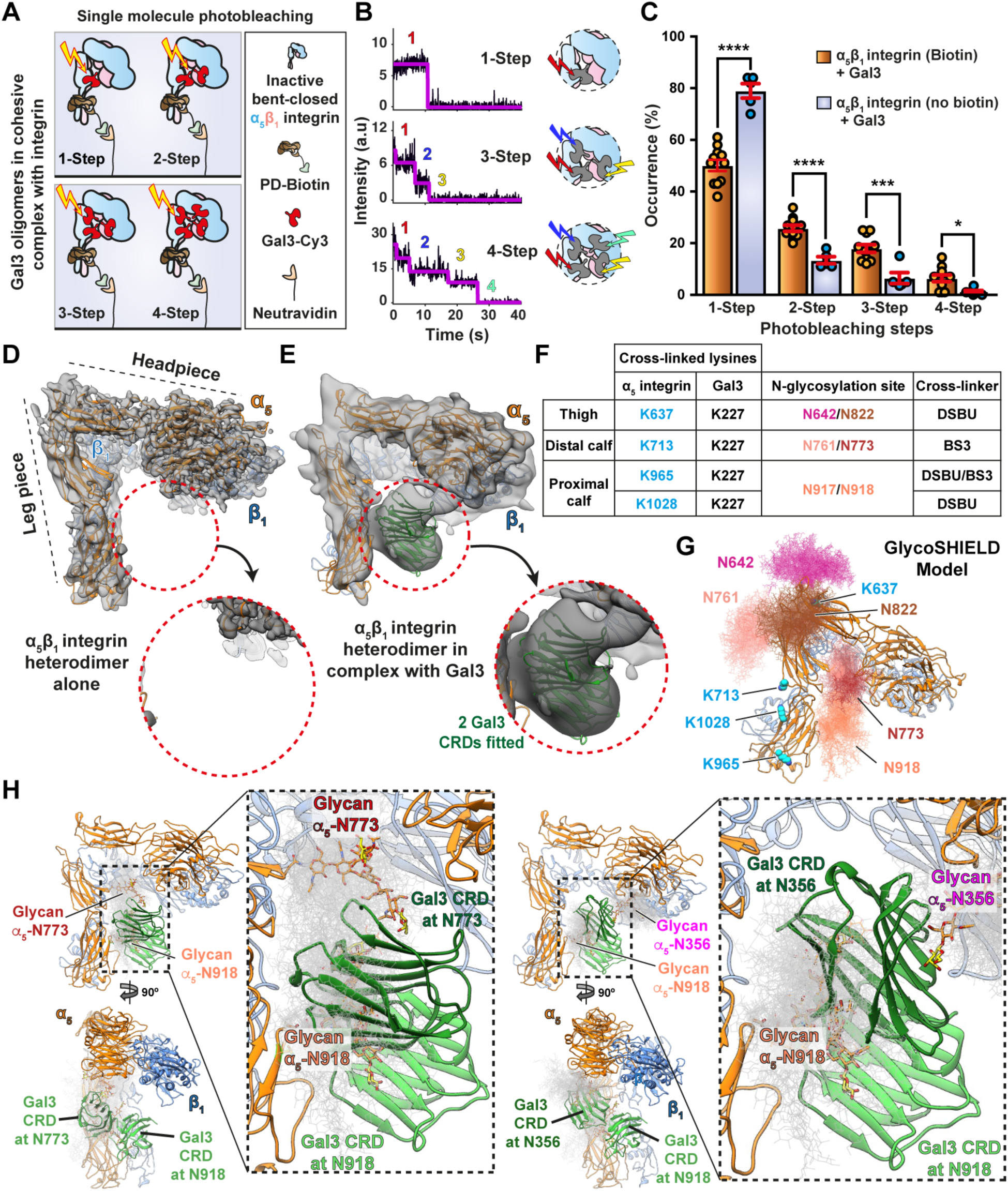
Structure of the inactive bent-closed α_5_β_1_ integrin in complex with Gal3. (**A**) Schematics of the different inactive α_5_β_1_ integrin-Gal3 cohesive complex configurations in photobleaching experiments (PD: peptidisc). (**B**) Qualitative visualization of single Gal3 molecule photobleaching step counting in these complexes. (**C**) Quantification in the indicated conditions of experiments as in (A,B). Peptidisc-embedded α_5_β_1_ that did not specifically bind to the neutravidin substrate (no biotin) served as controls. 11 fields for biotin condition, 5 fields for no-biotin condition. Means ± SEM, unpaired t-test; *P < 0.05, ***P < 0.0002, ****P < 0.0001. (**D**) Cryo-EM density map of peptidisc-embedded α_5_β_1_ integrin alone (gray) with modeled α_5_β_1_ integrin (orange for α_5_ and blue for β_1_ subunits). The red dashed circle indicates the density-free space between the head and leg pieces. (**E**) Cryo-EM density map of α_5_β_1_ integrin in complex with Gal3. Here, the red dashed circle clearly indicates the presence of additional cryo-EM densities into which two Gal3 CRDs (PDB 1KJL, green) were fitted. The density map between head and leg pieces was lowpass filtered to 15 Å for representation purposes. (**F**) Cross-linking proteomics on the α_5_β_1_ integrin-Gal3 complex. Cross-linked lysine positions from trans-peptides (Gal3 and α_5_β_1_ integrin) as well as the cross-linkers used are reported in the table. (**G**) Cross-linked lysines (in blue) and complex glycans in their vicinity (in red tones) were projected onto the α_5_β_1_ integrin structure from (D). GlycoSHIELD conformations were computed for each of these glycans, 100 possible conformations are shown each. (**H**) Model of α_5_β_1_ integrin-Gal3 complex as in (E). Left, GlycoSHIELD conformations are shown for glycans at N773 and N918 (left) or at N356 and N918 (right). Two Gal3 CRDs modeled within the cryo-EM density are highlighted along with glycans at N773 and N918 or N356 and N918 that fitted best. The other GlycoSHIELD conformations are shown in light gray.

To structurally characterize Gal3 binding to α_5_β_1_ integrin via cryogenic electron microscopy (cryo-EM), we first vitrified peptidisc-embedded α_5_β_1_ integrin alone (Figure S5A) and refined the structure of the inactive bent-closed conformation to resolutions of 3.7 Å in the headpiece and 4.7 Å in the leg piece, respectively (Figures S5B-J). A homology model of rat α_5_β_1_ integrin (Mirgorodskaya et al., 2022) was fitted individually into the density maps of the headpiece (flexible fit, Figure 6D and Figures S5K and S5L) and leg piece (rigid body fit, Figure 6D). The structure revealed a bent-closed conformation with an angle between headpiece and leg piece that is larger than for the bent-closed conformations of other integrins (Adair et al., 2023), and similar to human α_5_β_1_ integrin (Schumacher et al., 2021). There is no density bridge between the headpiece and the leg piece (Figure 6D and Figure S7D, circle).

In the next step, complexes between Gal3 and desialylated peptidisc-embedded inactive bent- closed α_5_β_1_ integrins were assembled, then detached from beads and immediately vitrified to minimize complex dissociation and aggregation (Figure S4K, Figure S6A-C). The cryo-EM density map of all particles (4.1 Å resolution, Figure S6A) revealed the presence of fragmented density between head and leg pieces of α_5_β_1_ integrin, indicative of Gal3 binding in this region (Figure S6A and compare Figures 6D, 6E and S7D, red dashed circles). To precisely position Gal3, we performed focused 3D classification and identified two subsets with densities at slightly different positions (Figure S6A). Subset 1 with the highest level of density (6.9 Å resolution, Figures 6E, S6A and S6D-F) comprised 13% of all particles, indicating a high level of heterogeneity.

These structural data pointed to the intriguing possibility that Gal3 oligomers might physically bridge membrane-proximal (leg piece) and distal (headpiece) α_5_β_1_ integrin parts that face each other in the inactive bent-closed conformer, with two distinct Gal3 CRDs fitting into the density bridge (Figure 6E). This bent-closed state would thereby be stabilized by a Gal3-mediated clamping effect. The propensity of α_5_β_1_ integrin to undergo the switch from the bent-closed to the extended ligand-bound conformational state was indeed found to be reduced on cells (Figure S7A) or micelles (Figure S7B) when Gal3 had been pre-bound into each system (schematics in Figure S7C). This was not observed with Gal31′Nter (Figure S7A), which reinforced the notion that Gal3 oligomerization was required for this clamping effect. A slight decrease of the angle between head and leg pieces occurred when clearly visible Gal3 densities were present on α_5_β_1_ integrin (Figure S7E), which is consistent with the hypothesis on a clamping effect.

To identify the glycans that are involved in Gal3-mediated clamping of the bent-closed conformational state of α_5_β_1_ integrin, we combined cross-linking proteomics, molecular modeling, and site-directed mutagenesis in subsequent experiments.

### Specifically inactive bent-closed α_5_β_1_ integrin requires defined glycans for endocytic uptake

Cross-linking mass spectrometry revealed the proximity between position 227 in the CRD of Gal3 with positions in the membrane distal and membrane-proximal calf regions of the leg piece of α_5_ integrin (Figures 6F and S8A). Several N-glycans at sites close to cross-linking positions, i.e., N642, N761, N773, N822, and N917/N918 of the α_5_ chain, are complex-type multi-antennae structures that are of galectin-binding competent nature, as determined by site-specific glycoproteomics on the same α_5_β_1_ integrin preparation from rat liver as used in the current study (Mirgorodskaya et al., 2022). Using GlycoSHIELD (Gecht et al., BioRxiv), possible conformations of these N-glycans were projected onto the α_5_β_1_ integrin model (Figures 6G and S8B).

Notably the glycan at N918 protrudes towards the head/leg interspace (Figure S8C), and its terminal galactose residues allowed for the fitting of a Gal3 CRD into the additional cryo-EM density (Figures 6E, 6H left, and S8D). Furthermore, a second Gal3 CRD could be placed into this density for interaction with a galactose residue on the N773 glycan on the membrane-distal calf (Figures 6E, 6H left, and S8D).

From these structural data we deduced that N-glycans from the membrane-proximal calf region of α_5_ integrin may be involved in Gal3 binding and oligomerization. To assess the functional relevance of these glycans, the corresponding glycosylation sites were mutated in human α_5_ integrin (termed ΔMP, Figures S8E and S8F), and expressed together with wildtype human β_1_ integrin in mouse kidney fibroblasts with a double knockout for endogenous α_5_β_1_ integrin (MKF- dKO) (Figure S8G). We found that α_5_β_1_ integrin cell surface levels were similar (mAb13) or even higher (9EG7) in ΔMP expressing cells, when compared to wildtype α_5_β_1_ integrin-expressing cells (Figure S8H), documenting efficient export from the endoplasmic reticulum and trafficking to the plasma membrane. The increased level of active ligand-bound α_5_β_1_ integrin at the cell surface may explain the observed increased size of focal adhesions (Figure S8I) and cell area (Figure S8J) in ΔMP expressing cells.

Despite efficient localization of ΔMP-mutant α_5_β_1_ integrin to the cell surface, the overlap of mAb13 (inactive bent-closed α_5_β_1_ integrin) with Gal3 dropped to similarly low levels on these ΔMP cells (Figure 7A) as observed with the oligomerization deficient Gal3ΔNter mutant on wildtype α_5_β_1_ integrin expressing cells (Figure S8K). This striking result suggests that the removal of the membrane-proximal glycosylation sites on the α_5_ chain diminishes the capacity of bent- closed α_5_β_1_ integrin to trigger Gal3 oligomerization. Consistently we found that the weak overlap of Gal3 with Gal3 oligomerization incompetent active ligand-bound α_5_β_1_ integrin (9EG7; Figures 4H and 4I) was not reduced any further on ΔMP-expressing cells (Figure 7A). Furthermore, the endocytic uptake of the inactive bent-closed conformational state (mAb13) of α_5_β_1_ integrin ΔMP mutant also was much reduced, when compared to that of wildtype α_5_β_1_ integrin, while no difference was observed for the active ligand-bound conformational state (9EG7) (Figure 7B).

**Figure 7.**
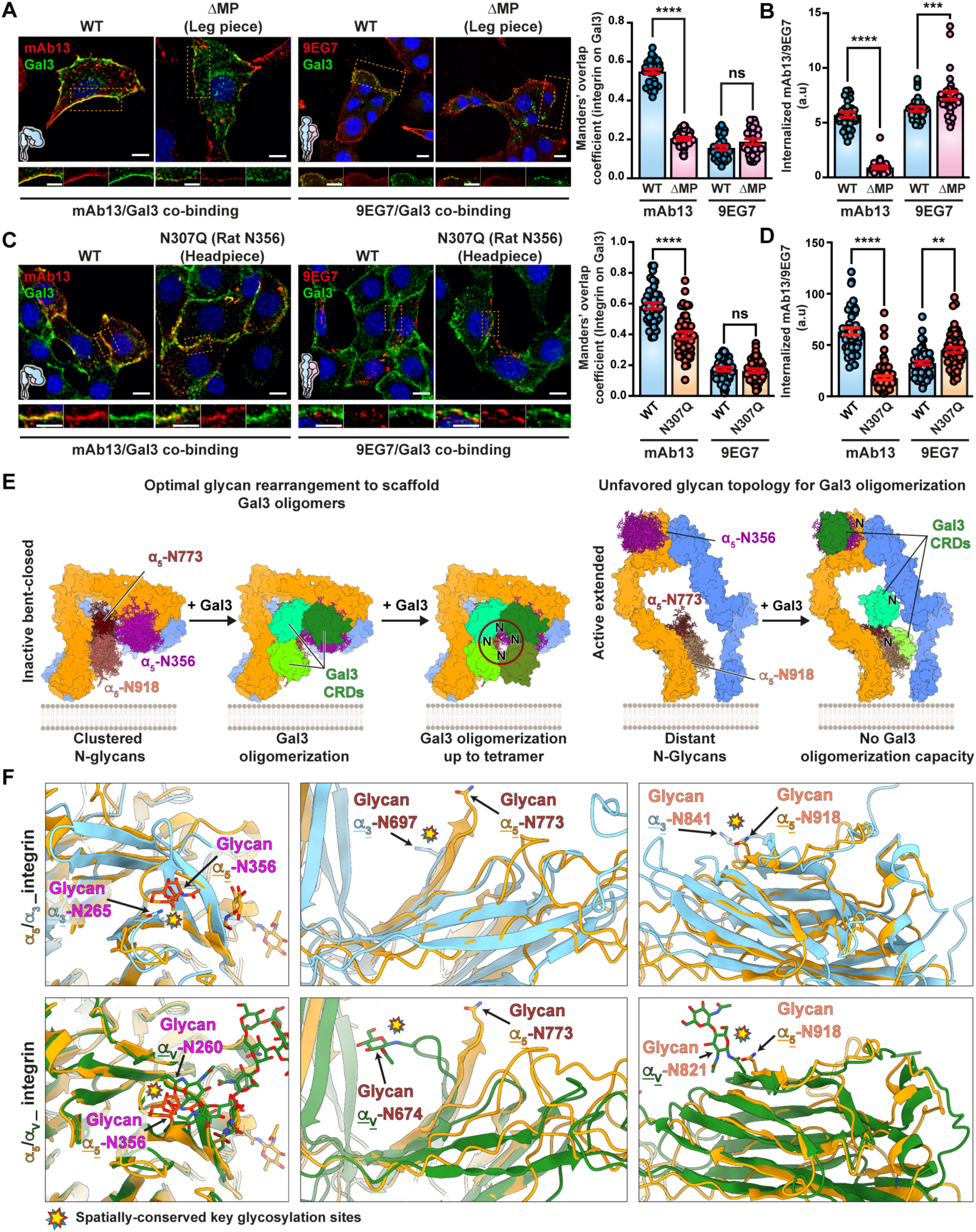
The inactive bent-closed conformation of α_5_β_1_ integrin presents an optimal glycan arrangement for nucleating Gal3 oligomerization. (**A**) Colocalization of Gal3 with mAb13 or 9EG7 antibodies after binding at 4 °C to α_5_β_1_ integrin-deficient MKF-dKO cells exogenously expressing heterodimers of wildtype or ΔMP (leg piece) mutant human α_5_ integrin with wildtype human β_1_ integrin. Nuclei in blue. Scale bars = 10 µm. 3 independent experiments, means ± SEM, unpaired t-test; ns = P > 0.5, ****P < 0.0001. (**B**) Endocytosis with ΔMP mutant. Continuous incubation for 10 min at 37 °C of mAb13 or 9EG7 with MKF-dKO cells under α_5_β_1_ integrin expression conditions as in (A). Internalized signals were quantified for 3 independent experiments, means ± SEM, unpaired t-test; ***P < 0.0002, ****P < 0.0001. (**C**) Colocalization experiments as in (A) in which position N307 (headpiece) of human α_5_ integrin (position N356 in rat) was mutated and expressed with wildtype human β_1_ integrin. Nuclei in blue. Scale bars = 10 µm. 3 independent experiments, means ± SEM, unpaired t-test; ns = P > 0.5, ****P < 0.0001. (**D**) Internalization experiments as in (B) on MKF-dKO cells expressing the heterodimer of human α_5_- N307Q integrin mutant with wildtype human β_1_ integrin. As in (B), only the uptake of the inactive conformational state of the integrin (mAb13) was inhibited under glycosylation site mutation conditions. 3 independent experiments, means ± SEM, unpaired t-test; **P < 0.002, ****P < 0.0001. (**E**) Working model. Only for the inactive bent-closed α_5_β_1_ integrin conformation are key glycosylation sites positioned such that Gal3 monomers are in an oligomerization-compatible distance. In the active extended conformational state, these glycosylation sites are spaced too distant from each other to nucleate Gal3 oligomerization. (**F**) The key glycosylation sites that have been identified on rat α_5_ integrin were used to search for corresponding sites (indicated by stars) in the predicted or solved structures of α_3_ or α_v_ (PDB: 4G1M) integrins, respectively.

These results demonstrate a dichotomic relationship again in which the membrane proximal glycosylation sites on the α_5_ chain are critical for the efficient colocalization with Gal3 and endocytic uptake of specifically the inactive bent-closed conformational state of α_5_β_1_ integrin. We then addressed the role of glycosylation sites in the headpiece.

The complex glycan at α_5_ chain position N356 (Mirgorodskaya et al., 2022) of the headpiece (N307 in human, Figure S8E) also points towards the additional cryo-EM density that we ascribed to Gal3 (Figure 6E). Based on GlycoSHIELD modeling, it was indeed possible to position another Gal3 CRD for interaction with a galactose residue on the N356 glycan, resulting in a Gal3 dimer on the N356 and N918 glycan that forms a bridge between headpiece and leg piece (Figure 6H, right). The corresponding α_5_ chain glycosylation site mutant N307Q, when co-expressed with wildtype β_1_ integrin in MKF-dKO cells, was again efficiently localized at the cell surface (Figure S8L). Yet, for the inactive bent-closed conformational state (mAb13), the overlap with Gal3 was significantly reduced (Figure 7C), and mAb13 uptake was largely inhibited (Figure 7D). In contrast, for the active ligand-bound conformational state (9EG7), overlap with Gal3 (Figure 7C) and endocytosis (Figure 7D) were similar between N307Q and wildtype α_5_β_1_ integrin expressing cells.

These data consolidate the dichotomy notion between inactive bent-closed and active ligand-bound conformational states of α_5_β_1_ integrin as to their interaction with Gal3 and their GL-Lect driven endocytic uptake into cells. Specifically, it appears that only on inactive bent-closed α_5_β_1_ integrin N-glycans from head and leg pieces are arranged such that Gal3 oligomerization can be nucleated (Figure 7E, left). In contrast, in extended ligand-bound α_5_β_1_ integrin conformer, these glycans would be too distant from each other to ensure such oligomerization (Figure 7E, right).

Also α_V_ and α_3_ integrins were found to be strong Gal3 interactors and cargoes of the retrograde route (Figure S8M). In both cases, N-glycosylation sites are present at equivalent position as the key α_5_ integrin sites N356, N773 and N918 (Figure 7F). In contrast, for α integrins that were poorly enriched in our retrograde and/or Gal3 interaction proteomics lists, such as α_2_ and α_6_ integrins (Figure S8M), no strict structural conservation could be detected for the N356, N773 and N918 sites (Figure S8N). The N-glycan signature that we have identified here as a trigger for Gal3 oligomerization may therefore apply beyond α_5_β_1_ integrin.

## Discussion

Based on our structural and functional data, we propose a mechanism according to which the conformational state of proteins can dynamically arrange the spatial organization of N-glycans to facilitate nucleation of Gal3 oligomers and drive endocytosic fate. We demonstrate this with α_5_β_1_ integrin for which only the non-ligand-bound bent-closed conformational state positions N-glycans ideally for nucleation of Gal3 oligomerization. We propose this conformational glycoswitch mechanism to be widely applicable to cell membrane resident glycoproteins.

Our study identifies Gal3 as the first extracellular factor that specifically interacts with the inactive bent-closed conformational state of an integrin. As a nucleator for Gal3 oligomerization, the bent-closed state may thereby be considered as an active player in the functional cycle of integrins, including Gal3 oligomer-driven GL-Lect endocytosis, retrograde trafficking to the Golgi apparatus, and subsequent polarized secretion to the leading edge of migrating cells to enter a new functional cycle (Shafaq-Zadah et al., 2016). This cycle is important for integrin-mediated functions, including cell adhesion and persistent cell migration (Shafaq-Zadah et al., 2016). Of note, Gal3 itself, which we identified here as a confirmed retrograde cargo, has also been shown to drive cell migration and invasion (Bojić-Trbojević et al., 2019; Boscher and Nabi, 2013; Furtak et al., 2001; Liu et al., 2012; MacDonald et al., submitted).

Our study focused on α_5_β_1_ integrin and members of the integrin family. Other Gal3 binding partners such as CD44 (Jamison et al., 2011; Suzuki et al., 2015) and epidermal growth factor receptor (Kaplan et al., 2016) also undergo conformational changes, potentially resulting in spatial rearrangement of N-glycans. Therefore, the identified conformational glycoswitch mechanism may apply more generally.

Our work sheds new light on the nature and shape of Gal3 oligomers, which may have wider relevance to the large family of galectins and other glycan-binding proteins. Based on biochemical evidence, Gal3 oligomers have been proposed to exist as ill-defined pentamers (Ahmad et al., 2004), as higher order assemblies (Lepur et al., 2012), or as tetramers with inconsistent features made from N-terminally truncated protein (Flores-Ibarra et al., 2018). Here, using tissue-derived reconstituted α_5_β_1_ integrin or the surface of living cells for assembly, we demonstrate that full-length Gal3 can form well-defined dimers, trimers and tetramers. Pentamers were not found, which could either mean that they are absent altogether, have very low abundance, and/or are very unstable.

The ring-shaped oligomeric organization with a central pore that orients four Gal3 with glycan binding sites pointing in the same direction on the outside are striking hallmarks of the Gal3 tetramer that we have discovered. These elements are shared with the GSL-binding subunits of bacterial Shiga and cholera toxins and the VP1 protein of simian virus 40, which like Gal3 drive GSL-dependent narrow membrane bending and the biogenesis of tubular endocytic pits from which CLICs are formed (Ewers et al., 2010; Kabbani et al., 2020; Römer et al., 2007). These elements may represent the structural signature of GL-Lect driven endocytic processes. How the molecular organization of Gal3 oligomers on inactive bent-closed α_5_β_1_ integrin, as revealed here, reaches onto the membrane to drive GSL-dependent membrane deformation remains to be established in future studies.

The N-terminal proline-rich domain of Gal3 is not visible in our tetramer model, which indicates that it might remain unstructured (Birdsall et al., 2001). This domain has been described to undergo biomolecular condensate formation (Chiu et al., 2020; Lin et al., 2017; Zhao et al., 2021), which would be one possible mechanism by which Gal3 oligomers might be stabilized in a way that allows them to accommodate variable numbers of Gal3 molecules.

While our study reveals the mechanism of Gal3 oligomer assembly at the cell surface, it remains to be established how oligomer disassembly is operated. Our observation that Gal3 remains colocalized with bent-closed α_5_β_1_ integrin in endosomes and is itself transported to the Golgi apparatus points to the possibility that Golgi-specific functions may contribute to the disassembly process, e.g., acidification (von Mach et al., 2014) and sialylation on N-glycans (Zhuo and Bellis, 2011), as we show elsewhere (MacDonald et al., submitted). This would then free α_5_β_1_ integrin and reset the protein for *de novo* activation after its polarized secretion to the leading edge (Hou et al., 2016; Pan and Song, 2010; Seales et al., 2005; Shafaq-Zadah et al., 2016; Yu et al., 2013).

Our study identifies the inactive bent-closed conformational state of α_5_β_1_ integrin as a nucleator of Gal3 oligomerization to selectively drive endocytosis. This finding provides a hitherto undescribed scenario for recognition of patterns of glycans on glycoproteins and the importance of considering the rearrangement of glycans on glycoproteins for the acute regulation of functions and dynamics at the cell surface. Elsewhere we propose a complementary mechanism for galectin-mediated endocytosis of membrane glycoproteins in which acute desialylation at the cell surface by growth factor signaling exposes high affinity ligands for galectin binding (MacDonald et al., submitted). Since α_5_β_1_ integrin function is sensitive to sialylation (Hou et al., 2016; Pan and Song, 2010; Pretzlaff et al., 2000; Seales et al., 2005), the protein appears as a likely client for the desialylation mechanism as well. These two mechanisms are indeed complementary and not mutually exclusive as they may function in a staged manner, i.e., growth factor induced N-glycan desialylation followed by conformational state-specific rearrangements in the N-glycan landscape to drive nucleation of galectin-oligomers. Sialylation would thereby set the fraction of inactive bent-closed α_5_β_1_ integrin that is available for nucleating Gal3 oligomers.

In conclusion, our work highlights the dynamic nature of changes in glycans at the cell surface, which calls for profound rethinking of the roles and functions of glycosylation on membrane glycoproteins.

## Materials and Methods

### Cells and tissues

HeLa cells, HeLa cells stably expressing the TGN-localized GalT-GFP-SNAP fusion protein (Shi et al., 2012), RPE-1 cells, genome edited RPE-1 cells stably expressing AP2- GFP (see below), α_5_β_1_ integrin double KO mouse kidney fibroblast (MKF-dKO, kindly provided by Reinhard Fässler). Rat livers (Charles Rivers), *E. coli* strain Rossetta2-pLysS (Novagen).

### Reagents

WGA-agarose column (Sigma-Aldrich, Ref. 61768-5 mL), FN-III_9-10_-agarose column (GE Healthcare), cRGD (CliniSciences, Ref. A8164), soluble fibronectin (Sigma, Ref. F0895- 2mg), recombinant FNIII 9-10 fibronectin fragment (Dransart et al., 2022), NHS-HiTrap (GE Healthcare, Ref. 17071701), HisPur™ Cobalt Resin (Thermofisher, Ref. 89965), protein G-sepharose beads (Sigma, Ref. P3296), GFP-Trap MA beads (Chromotek, Ref. gtma-20), Bio- Beads™ SM2 adsorbent media (Biorad, Ref. 152-3920) 300-400 mesh carbon-coated copper grids for electron microscopy (Delta Microscopy, Ref. DG400-Cu), Quantifoil Cu 300 mesh QF2/1 grids for cryo-EM, HEPES 1M pH 7-7.4 (Sigma-Aldrich, Ref. H0887), protease inhibitors (Sigma-Aldrich, Ref. P8849), EDTA pH 8, chicken egg phosphatidylcholine (ePC, Avanti Polar, Ref. 840051C), Brain PS (Avanti Polar, Ref. 840032P), Transferrin-Alexa546 (Tf-A546) (Invitrogen, Ref. T23364), recombinant purified Gal3, Gal3-His, Gal3ΔNter (Lakshminarayan et al., 2014), Ingenio® electroporation buffer (Mirus, Ref. MIR 50111), HiPerFect transfection reagent (Qiagen), siCHC (Qiagen, Refs. SI00299880S1; SI00299873S1), hamster anti-rat α_5_ and β_1_ integrin primary antibody (BioLegend, Ref. 103902/102202), HRP-coupled secondary anti-hamster antibody, mAb13 antibody (BD Bioscience, Ref. 552828), 9EG7 antibody (BD Bioscience, Ref. 553715), mAb16 and SNAKA51 antibodies (kindly provided by Patrick Caswell), TS2/16 antibody (BioLegend, Ref. 303036), anti-SNAP-tag antibody (New England Biolabs, Ref. P9310S), anti-clathrin heavy chain antibody (BD Bioscience, Ref. 610500), anti- paxilin antibody (BD Bioscience, Ref. 610051), anti-α tubulin antibody (BD Bioscience, Ref. T5168), anti-vps35 antibody (kind gift from Juan Bonifacino), anti-vps26 antibody for immunoblotting (Abcam, ref. ab6051) or immunofluorescence (Abcam, ab23892), NHS-ATTO- 488 (ATTO-TEC, Ref. AD488-31), NHS-Alexa647 (Invitrogen, Ref. A20006), secondary anti-mouse-HRP (Beckman Coulter, Ref. 715-035-151), secondary anti-rabbit-HRP (Beckman Coulter, Ref. 711-035 -152), secondary anti-rat-HRP (Beckman Coulter, Ref. 712-035-153), secondary anti-rat Cy3 (Beckman Coulter, Ref. 712-166-153), HRP-NHS (AAT Bioquest, Ref. 11025), sialidase (neuraminidase from *Arthrobacter ureafaciens*, Roche Ref. 10269611001), BG-GLA-NHS (New England Biolabs, Ref. S0151S), ciliobrevin D (CBD) (Merck, Ref. 250401), Genz- 123346 (Sigma, Ref. 5382850001), Gal3 inhibitor compound GB0149-03 (Galecto Biotech; 1,1′- sulfanediyl-bis-{3-deoxy-3-[4-(butylaminocarbonyl)-1*H*-1,2,3-triazol-1-yl]-β-D galactopyranoside}, referred to as I3), β-D-lactose (Sigma, Ref. L3750), NHS-Cy3 (GE, PA23001), Cy3 mono-reactive succinimidyl ester (Cytivia, Ref. GEPA23001), BS3-d0/BS3-d4 ((bis(sulfosuccinimidyl)suberate-d0/d4, Thermoscientific, Ref. 21590 and 21595), DSBU (Disuccinimidyl Dibutyric Urea, Thermoscientific, Ref. A35459, SNAP-Cell® Block (New England Biolabs Ref. S9106S), glutaraldehyde, protein ladder (Thermoscientific, Ref. 26619). Triton X-100™ (Anatrace, Ref. T1001-500 mL), DAB (Sigma Aldrich, Ref. D8001), ascorbic acid, Nonidet P-40 (NP40, Sigma, Ref. 21-3277), MSP1D1 protein scaffold (MSP1D1-His, Cube Biotech, Ref. 26112), peptidiscs (MSP1D1-derived scaffold peptidisc His/Biotin, Peptidisc Lab), dialysis cassettes 10 kDa cut-off (MWCO, Thermo Scientific), 100 kDa cut-off concentrator (GE Healthcare, Vivaspin 500 (Merck, Ref. Z614092-25EA), PD-10 desalting column (GE, Ref. 17- 0851-01), Zeba spin column 7 kDa cut-off (Thermo Fisher, Ref. 89882), 4-15% Stain-Free™ pre- casted polyacrylamide gels (BioRad, Ref. 4561084), ECL reagent, 5 mm coverglass (Electron Microscopy Sciences, Ref. 72195-05).

### Media and buffers

DMEM high glucose (Thermofisher, Ref. 41965039), DMEM-F12 Gibco (Thermofisher, Ref. 11320033), PBS++ (PBS supplemented with 1 mM MgCl_2_ and 0.5 mM CaCl_2_, pH 7.4), HEPES buffer (20 mM HEPES, 150 mM NaCl). Hepes-Tx100 buffer: HEPES buffer supplemented with 0.2% (v/v) Triton X-100, β-D-lactose solution (150 mM, iso-osmolarized), I3 solution (10 μM or 50 μM), TNE buffer (10 mM Tris, 150 mM NaCl and 5 mM EDTA), TNE 1% NP40 supplemented with proteases inhibitor cocktail (lysis buffer), acid wash buffer (glycine 0.5 M, pH 2.2). PFA 4% solution (Electron Microscopy Sciences, Ref. 1570). 3x non-reducing SDS sample buffer (2 M Tris/HCl, pH 6.8, 20% SDS, 30% glycerol, 0.03% phenol red), 3x non-reducing low SDS sample buffer (2 M Tris/HCl, pH 6.8, 2% SDS, 30% glycerol, 0.03% phenol red), BSA-saponin buffer (0.5% saponin, 2% BSA, in PBS), HEPES buffer (20 mM HEPES, 150 mM NaCl). HEPES/Tx-100 buffer: HEPES buffer supplemented with 0.2% (v/v) Triton X-100.

### Equipment

Transmission electron microscope (TEM) 80 kV (Tecnai Spirit, ThermoFisher, USA), equipped with QUEMESA camera (Olympus), TEM 80 kV (TEM 900, Zeiss) equipped with a Morada G2 camera (Olympus), TEM 120 kV (Talos L120C, ThermoFisher), equipped with Ceta16M camera (ThermoFisher), TEM 300 kV (Titan Krios, ThermoFisher), equipped with K3 direct electron detector with energy filter (Gatan), TEM 300 kV (Titan Krios, ThermoFisher), equipped with Falcon3 direct electron detector (ThermoFisher), cryoplunger (Vitrobot Mark IV, ThermoFisher), confocal microscope (A1RHD25 microscope, Nikon Imaging Center, Curie Institute), lattice light sheet microscope (3i, Intelligent Imaging Innovations). BioRad ChemiDoc for protein detection (chemiluminescence and fluorescence detection). Microtime 200 system (PicoQuant GmbH, Germany) for fluorescence lifetime imaging and correlation spectroscopy measurements. CHI760e Electrochemical Workstation (CH Instruments, USA) for electrochemical impedance spectroscopy. nanoAcquity UPLC device (WatersCorporation, Milford, MA, USA), Q-Exactive HF-X mass spectrometer (ThermoFisher Scientific, Waltham, MA, USA).

### Recombinant protein expression

Human recombinant Gal3-6xHis and Gal3ΔNter-6xHis with aa115-250 of BC053667 were cloned in pHis-Parallel2 and purified as described (Ivashenka et al., 2022; Lakshminarayan et al., 2014). Briefly, for cloning, Gal3 and 6xHis were separated by a Leu-Glu linker at the C-terminus. Proteins were expressed overnight at 20 °C in Rossetta2-pLysS using LB-media with 60 μM IPTG, and purified with cobalt resin affinity chromatography and gel filtration (Superdex75 16x60) in PBS at pH 7.4. Coupling with Alexa488 sulfodichlorophenol ester (Alexa488-Gal3) or Cy3 mono-reactive succinimidyl ester (Gal3-Cy3) was performed overnight at 16 °C using a 4-fold molar dye excess in PBS and purified with PD10 columns. The labeling efficiency was 1 to 1.5. Human recombinant Gal3-TEV-6xHis was cloned in pHis- Parallel2 without linker between Gal3 and TEV cleavage site, and 26 amino acids between the TEV site and 6xHis (FW:5’ GGAATTCCATATGGCAGACAATTTTTCGCTCCATGATGCG, RV:5’ CTGGATCCGCCCTGAAAATACAGGTTTTCTATCATGGTATATGAAGCACTGG).

Protein expression was performed as for Gal3-6xHis. The bacterial pellet was resuspended in buffer-A (20 mM HEPES pH 7.8, 200 mM NaCl, 80 mM lactose, 10 mM imidazole, 1 mM TCEP), sonicated, centrifuged for 60 min at 75000 g, and the supernatant was incubated for 1 h at 4 °C with Ni-Sepharose-Fast Flow beads (Cytiva). The beads were then washed with buffer-A containing 300 mM NaCl, and eluted in buffer-B (20 mM HEPES pH 7.8, 20 mM NaCl, 40 mM lactose, 0.5 mM TCEP, 500 mM imidazol pH 7.8). The eluted protein was diluted to 2 mg/mL and dialyzed in buffer-C (20 mM HEPES, 25 mM NaCl, 40 mM lactose, 0.5 mM TCEP, 0.5 mM EDTA) to remove imidazole. 20 µg shTEV-His protease per mg of Gal3 was injected into the dialysis chamber. After 24 h at 4 °C, the dialysis chamber was transferred for 8 h to a lactose and EDTA-free buffer-D (20 mM HEPES 7.3, 150 mM NaCl, 50 μM TCEP). ShTEV-His and non-cleaved Gal3-TEV-6xHis was removed by incubation with a cobalt resin. We noticed highly purified Gal3 was very sticky to Ni-NTA. The non-bound protein fraction was concentrated and purified by FPLC-gel filtration (Superdex75 16x60) using buffer-D without TCEP, with an elution peak at 68 mL. When required, Gal3 was fluorophore-labeled as for Gal3-6xHis using 20 mM HEPES, 150 mM NaCl pH7.3, snap-frozen and stored at -80 °C.

### Preparation of antibody-HRP conjugates

100 μg of mAb13 or 9EG7 antibodies were added to lyophilized HRP-NHS (300 μg) at a final molar ratio of 1/1 and incubated for 2 h at room temperature. Unreacted HRP was removed using 100 kDa cut-off concentrator.

### mAb13-HRP/9EG7-HRP internalization and sample preparation for electron microscopy

10 μg/mL of HRP-conjugated mAb13 or 9EG7 antibody were continuously incubated for 6- and 9- min at 37 °C with HeLa cells. Cells were immediately placed on ice, washed once with DMEM supplemented with 15 mM HEPES and 1% BSA, and twice with DMEM supplemented with 15 mM HEPES without BSA. Surface-bound antibodies were removed by incubation for 10 min at 4 °C with ascorbic acid solution of DAB (in DMEM, 15 mM HEPES, 1% BSA). Enzymatic reaction of internalized HRP was developed by incubation for 20 min at 4 °C with the same solution supplemented with H_2_O_2_. The cells were washed 3 times at 4 °C with DMEM supplemented with 15 mM HEPES, and fixed over night at 4 °C with 2.5% glutaraldehyde in PBS. Cells were washed 3 times at room temperature during 70 min with 0.1 M Na-Cacodylate in H_2_O. Membrane fixation was performed with 1% OsO4 in 0.1 M Na-Cacodylate in H_2_O. After 1 wash with 0.1 M Na-cacodylate in H_2_O and 2 washes with H_2_O, contrast was obtained by incubation for 45 min with aqueous 4% uranyl acetate solution. Cells were again washed with H_2_O. Samples were then dehydrated for 10 min at room temperature by incubation with increasing concentration of aqueous ethanol solutions (1 x 50% 5 min, 1 x 70% 5 min, 2 x 90% 10 min) and 3 x 100% anhydrous ethanol solution. Cells were finally embedded in LX112 resin, ultrathin 65 nm sections were obtained using a Reichert Leica UCT ultramicrotome, and mounted on Ni/formvar/carbon-coated grids for observations. Micrographs were acquired by electron microscopy.

### Clathrin heavy chain (CHC) cell surface co-immunoprecipitation

mAb13 or 9EG7 antibodies (10 μg/mL) were incubated for 30 min at 4 °C with RPE-1 cells. Excess antibody removed by washing with ice-cold PBS^++^, and cells were lysed with lysis buffer. Antibodies were pulled down from cleared lysates (post-nuclear supernatants) by overnight incubation with 40 μl bed volume protein G-sepharose at 4 °C. Beads were washed 3-times with TNE 0.1% NP40 and immunoprecipitated proteins were denatured by boiling (95 °C) in SDS sample buffer. Eluted proteins were separated on denaturing non-reducing SDS-PAGE gels and immunoblotted against mAb13 and 9EG7 (anti-rat-HRP) and CHC (anti-mouse-HRP).

### Antibody uptake

10 μg/mL of mAb13 and 9EG7 antibodies were continuously incubated for 5 or 10 min at 37 °C with RPE-1 cells, according to experimental conditions. These were shifted to 4 °C and excess of antibodies removed by washing with ice-cold PBS^++^. Cell surface bound antibodies were removed with 3 acid washes of 45 sec each. Acidic pH was then neutralized with 3-times ice-cold PBS^++^ washes, cells were fixed in 4% PFA, followed by secondary antibody labeling (see below) and confocal microscopy.

### Exogenous Gal3 concentration

200 nM of Gal3 were typically used for cellular binding and uptake assays, as analyzed by confocal microscopy imaging. For experiments in which cargo internalization was stimulated by Gal3 addition (Figures 2H, 2I and S3A), we have used a range of concentrations from 20 nM to 400 nM. Note that a substantial increase of mAb13 uptake was already observed with only 20 nM of Gal3. In microcavity array supported lipidic bilayer (MSLB) experiments, low Gal3 concentration from 0.2 to 37 nM were used (Figure 4C-D and Figure S4B). In a previous study, the endocytosis of CD44 was indeed stimulated with as little as 0.3 nM of exogenous Gal3 on mouse embryonic fibroblasts that had been depleted for endogenous Gal3 (Lakshminarayan et al., 2014). For the Gal3 oligomer experiments on reconstituted a5b1 integrin (Figure 4H-I) and on cells (Figure 5A-D), the μM Gal3 concentrations were used to obtain maximal amounts of oligomer material.

### Transferrin and Gal3 uptake

5 μg/mL of Tf-A546 or 200 nM of Alexa488-Gal3 were continuously incubated for 5 or 10 min at 37 °C with RPE-1 cells, according to experimental conditions.

### Immunofluorescence

Cells were fixed with 4% PFA at 4 °C for 5 min and additional 5 min at room temperature. Excess of PFA was then quenched with 50 mM NH_4_Cl, followed by incubation for 30 min at room temperature with BSA-saponin saturation/permeabilization solution (intracellular immunostaining), or only saturation with PBS, 0.2% BSA (plasma membrane labeling). Cells were incubated for 30 min at room temperature either with primary and secondary antibodies for the labeling of cellular antigens, or only with secondary antibody to detect primary antibodies that had been put in contact with living cells (uptaken antibody). Coverslips were mounted on slides with Mowiol supplemented with DAPI. All fixed cell immunofluorescence images were acquired with a Nikon A1RHD25 confocal microscope. The quantification of signal was measured using the ImageJ program, and displayed as mean intensity per cell after maximal projection of Z-stacked images. For co-localization analysis, the JACoP plugin run in ImageJ was used to measure Manders’ coefficient signal co-occurrences. This quantification method calculates the percentage of signal from one channel that overlaps with signal from the other channel. To do so, it is of importance to image samples with high signal to noise ratios, and to select pixels from both channels that are relevant from the biological perspective (Pike et al., 2017).

### siCHC transfection

HeLa cells stably expressing Golgi-localized GalT-GFP-SNAP were transfected with 8 nM of siCHC (or of siCtrl) to inhibit clathrin heavy chain expression, using HiPerFect reagent. Antibody uptake and retrograde trafficking experiments were performed 48 h after transfection. Inhibition level of CHC expression was assessed by immunoblotting.

### EPS15DN-GFP transfection

RPE-1 cells were electroporated with 10 μg EPS15DN-GFP encoding plasmid for 24 h expression prior to performing antibody uptake experiments. GFP positive signal (GFP+) served to identify cells that expressed the dominant negative EPS15 mutant, and GFP-negative cells (GFP^-^) served as internal controls.

### Ciliobrevin D (CBD) treatment

RPE-1 cells were pre-treated for 30 min at 37 °C with 50 μM CBD. CBD was then kept during all subsequent incubations at 37 °C. For experiments with pre- loaded exogenous Gal3, CBD pre-treatment was followed by sequential incubation of 200 nM Gal3 and 10 μg/mL of mAb13 or 9EG7 antibodies on RPE-1 cells for 30 min at 4 °C, before further incubation at 37 °C for 10 min.

### CRISPR of RPE-1 AP2-eGFP

CRISPR-Cas9 genome editing of RPE-1 cells to generate isogenic mTagGFP-labeled AP2 was performed by electroporation of the following molecules. Homology directed repair template: Purified AP2-mTagGFP fragment after Snf-I digestion of pMK-RQ AP2M1-mTagGFP (vector: gift of Guillaume Montagnac, mTagGFP inserted between Ser236 and Glyc237 of AP2M1 (NM_004068.4), mTagGFP-spacer at N-term: GSTGGS, at C-term: AGSGT), and four gRNAs in pX330-Cas9, designed with CHOPCHOP (Labun et al., 2021). gRNA-2-FW: 5’CACCgagaagaggtctcattggtac, gRNA-2-RV: 5’AAACgtaccaatgagacctcttctC, gRNA-3-FW CACCgattgcttcccgctgcaagc, gRNA-3-RV AAACgcttgcagcgggaagcaatcC. Transfected cells were expanded for 2 weeks. The genome-edited population was identified by three cycles of FACS sorting and expansion, and finally tested for colocalization between AP2-mTagGFP and Tf- Alexa647 (ThermoFisher Scientific) during endocytic pit formation.

### mAb13-ATTO488 and 9EG7-ATTO488 conjugates

A 10-fold molar excess of NHS-ATTO488 compound was incubated for 2 h at 21 °C in preservative-free PBS buffer with the corresponding antibodies. Unreacted NHS-ATTO488 was first neutralized for 10 min at 21 °C with 10 mM Tris, and further eliminated using 7 kDa cut-off Zeba spin desalting columns.

### Dynamic mAb13 and 9EG7 tracking by lattice light sheet microscopy (LLSM)

Endocytosis experiments were performed in RPE-1 cells stably expressing AP2-GFP (RPE-1 AP2-mTagGFP), as previously described with minor modifications (Renard et al., 2020). In brief, RPE-1 AP2- mTagGFP cells were seeded 24 h before imaging on 5 mm #1.5 thickness cover glass. The medium was changed to CO_2_-independent lattice light-sheet (LLS) imaging medium (phenol-red free DMEM, high glucose, glutamax, supplemented with sterile 1% BSA, 0.01% penicillin and streptomycin, 1 mM pyruvate, and 20 mM HEPES pH 7.3). Cells were dipped for 2 min at room temperature in a tube either containing 5 μg/mL of mAb13-Cy3, 9EG7-Cy3, or Tf-Alexa546, diluted in LLS-imaging medium. After one wash in LLS-medium, coverslips were transferred into the imaging chamber of the LLS-microscope, kept at 27 °C. mAb13-Cy3/AP2-GFP and 9EG7- Cy3/AP2-GFP co-tracks were dynamically monitored by LLSM.

### Gal3/mAb13 and Gal3/9EG7 dynamic co-tracking by lattice light sheet microscopy (LLSM)

Galectin-3/integrin co-binding was performed in RPE-1 cells. These experiments were imaged as previously established with minor modifications (Renard et al., 2020). 5 mm #1.5 thickness coverslips seeded with RPE-1 cells were sequentially dipped for 2 min at 4 °C in a solution of Gal3-Cy3 (200 nM, in LLS-imaging medium), once washed in LLS-imaging medium and then plunged at 4 °C in a tube either containing 5 μg/mL of mAb13-ATTO488 or 9EG7-ATTO488, diluted in LLS-imaging medium. After one LLS-medium wash, coverslips were transferred into the imaging chamber. Co-tracks of Gal3-Cy3/mAb13-ATTO488 and Gal3-Cy3/9EG7-ATTO488 were dynamically monitored by LLSM and further processed.

### LLSM acquisition

Acquisitions in LLS-imaging medium were performed at 27 °C for increased stability of the optical system. 4D acquisition started within 2–4 min using a commercial LLSM of 3i (Denver, USA), as previously described (Chen et al., 2014). Cells were scanned incrementally with a 20 µm light sheet in 600 nm steps using a fast piezoelectric flexure stage equivalent to ∼325 nm with respect to the detection objective, and were imaged using two sCMOS cameras (Orca-Flash 4.0; Hamamatsu, Bridgewater, NJ). Excitation was achieved with 488 nm (Sapphire Coherent) or 560 nm (MPB Communications) diode lasers at 10-20% acousto-optic tuneable filter transmittance with 300 mW (initial box power) through an excitation objective (Special Optics 28.6× 0.7 NA 3.74-mm water-dipping lens) and detected via a Nikon CFI Apo LWD 25× 1.1 NA water-dipping objective with a 2.5× tube lens. LLSM imaging was performed using an excitation pattern of outer NA equal to 0.55 and inner NA equal to 0.493. A composite volumetric dataset of 60 slices per cell was acquired within 1.5-1.8 s using 10 ms exposure per slice and channel for mAb13-Cy3/AP2-GFP or 9EG7-Cy3/AP2-GFP co-tracking experiments, and within 2-3 s using 10-20 ms exposure time per slice and channel for Gal3-Cy3/mAb13-ATTO488 and Gal3- Cy3/9EG7-ATTO488 co-tracking experiments. 80-120 time points were acquired per cell. Raw images of obtained datasets were quantitatively analysed using an adapted version of the previously published cmeAnalysis3D software (Renard et al., 2020), and as described in the following section.

### Quantitative analysis and visualization of mAb13/AP2 or 9EG7/AP2 co-tracking experiments

Post-processing of raw data volumes was carried out as described previously (Renard et al., 2020). Automated detection of AP2-coated structures or punctate structures of fluorescently labeled cargoes in 3D (mAb13-Cy3 antibody, 9EG7-Cy3 antibody, and Tf-Alexa546) was performed by numerical fitting with a model of the microscope point spread function (PSF), as described previously (Aguet et al., 2016). Automated tracking of cargo and clathrin was calculated using the u-track software package (Jaqaman et al., 2008), as part of cmeAnalysis3D software (Aguet et al., 2016), which was implemented in Matlab 2021b. AP2 and cargo positions were exploited to map the membrane shape, using the Matlab function of alphaShape (Renard et al., 2020). We then calculated the displacement of each cargo from the plasma membrane. The distinction between point-like structures moving inside the plasma membrane and of internalized molecules, whether AP2-positive or AP2-negative, was made using membrane position detection. An event was counted as endocytic uptake if an object underwent a net displacement of at least 150 nm from the initial position inside the membrane proximal zone (Renard et al., 2020). The membrane proximal zone was limited by the contour of the alphaShape and a line 400 nm inwards of the cell (Renard et al., 2020). To calculate the number of AP2-positive and AP2-negative events, the presence of AP2 was assessed for each qualified internalization event of the cargo channel using the cmeAnalysis3D software. Only tracks with durations of more than 8 sec were used for this analysis. The calculation of the lifetime distributions and intensity cohorts were performed as described previously (Aguet et al., 2016). The raw LLSM images, used in Figures S3I and S3L and Movies S3-6 were deconvolved using LLSpy v0.4.8 (https://doi.org/10.5281/zenodo.3554482). Movies S3 and S4 were rendered and visualized using Imaris software 9.82. Movies S5 and S6 were rendered and visualized using ImageJ (Schneider et al., 2012)/Fiji 1.53c (Schindelin et al., 2012). The analysis code can be found as part of the Github repository of llsmtools in https://github.com/francois-a/llsmtools/. Statistical analyses were performed using Prism v9.4.1 software (Graphpad Inc).

### Quantitative analysis and visualization of mAb13/Gal3 or 9EG7/Gal3 co-tracking experiments

Post-processing of raw data volumes was carried out as described (Renard et al., 2020). Automated detection of punctate structures of Gal3-Cy3 and cargoes (mAb13-ATTO488 and 9EG7-ATTO488 antibodies) in 3D was performed by numerical fitting with a model of the microscope point spread function (PSF) as described previously (Aguet et al., 2016). Automated tracking of cargo and Gal3 was calculated using the u-track software package (Aguet et al., 2016; Jaqaman et al., 2008), which was implemented in Matlab 2021b. To calculate the number of Gal3-positive and Gal3-negative events, the presence of Gal3 was assessed for each qualified internalization event of the cargo channel using the cmeAnalysis3D software. Cargo tracks were considered Gal3-positive if they co-localized more than 10 sec with Gal3. Only tracks with duration of more than 10 timepoints were used for the analysis. Speed and distance of co-tracks were estimated using positions from cmeAnalysis3D software. The raw LLSM images, used in Figure 2D and Movies S1 and S2, were deconvolved using LLSpy (v0.4.8) (https://doi.org/10.5281/zenodo.3554482) before video rendering and visualization using napari (v0.4.12) (doi: 10.5281/zenodo.3555620). Statistical analyses were performed using Prism v9.4.1 software (Graphpad Inc).

### I3 treatment

Cells were pre-treated with 10 μM I3 for 5 min at 37 °C. I3 was washed out with PBS^++^ for acute endocytosis assays, or kept during subsequent incubations for retrograde trafficking experiments.

### Incubations with exogenous Gal3/Gal3ΔNter3

For antibody uptake assays, 10 μg/mL of mAb13 or 9EG7 antibodies were incubated for 30 min at 4 °C with RPE-1 cells. Excess antibodies were removed by washing with PBS^++^, and cells were then shifted to 37 °C in the presence of the indicated concentrations of exogenous Gal3 or Gal3ΔNter in serum-free DMEM-F12 medium. For retrograde transport assays, HeLa cells stably expressing GalT-GFP-SNAP were continuously co-incubated for 4 h at 37 °C with 10 μg/mL of BG-coupled mAb13 antibody and 200 nM of Gal3 in serum-free DMEM-F12 medium.

### Genz treatment

RPE-1 cells were continuously incubated for two days with 5 μM Genz-123346 in DMEM-F12 medium containing 5% FCS, prior to performing mAb13 and 9EG7 antibody uptake assays.

### Gal3/Gal3ΔNter and mAb13/9EG7 or mAb16/SNAKA51 co-binding and co-uptake assays

Exogenous Gal3 or Gal3ΔNter (200 nM) were incubated for 30 min at 4 °C with RPE-1 cells in serum-free DMEM-F12 medium. Excess of Gal3 or Gal3ΔNter was removed by washing with the same ice-cold medium. 10 μg/mL of integrin antibodies (mAb13 or 9EG7 for β_1_ integrin, mAb16 or SNAKA51 for α_5_ integrin) were then incubated for 30 min at 4 °C with the same cells that already had been pre-incubated with Gal3. Excess antibodies were removed by washing, and cells were either fixed in 4% PFA (co-binding assay) or shifted for 10 min to 37 °C (co-uptake assay). For the latter, residual cell surface accessible Gal3 was removed by incubations at 4 °C with 150 mM β-D-lactose (3 times, 5 min), and residual cell surface accessible integrin antibodies by acid washes. Immunofluorescence was then performed as described above.

### Gal3 and mAb13/9EG7 co-binding and co-uptake assays on micropatterned RPE-1 cells

Line patterns were produced, and cells seeded as previously described (Shafaq-Zadah et al., 2016). Coverslips (25 mm) were micropatterned with 9 μm-wide lines and covered with 5 μg/mL fibronectin. RPE-1 cells (30,000 per well) were seeded onto these coverslips and left to adhere in the incubator for at least 5 h before further manipulations. Gal3 and mAb13/9EG7 co-binding and co-uptake were performed as described above.

### Gal3-CFP/mAb13 or 9EG7 cell surface co-immunoprecipitation experiments

10 μg/mL of mAb13 or 9EG7 antibodies were incubated for 30 min at 4 °C with transiently Gal3-CFP expressing HeLa cells. Excess of antibodies were removed with 3-times ice-cold PBS^++^ washes and cells were lysed in lysis buffer. Cleared lysates were incubated with 30 μl slurry of GFP-Trap beads for overnight pull-down at 4 °C. After 3 washes in TNE 0.1% NP40, bound proteins were eluted from beads, and denatured at 95 °C heating in SDS sample buffer. Samples were loaded onto SDS-PAGE gels, immunoblotted with anti-rat HRP-antibodies against the bound antibodies (co-IP mAb13 and 9EG7). Antibody levels were quantified. Gal3-CFP fluorescence signals (pulldown) served as loading controls.

### mAb13-BG and Gal3-BG conjugates

BG-GLA-NHS compound was incubated for 6 h at 4 °C in preservative-free PBS buffer with a 10-fold molar excess over antibodies, or for 2 h at 21 °C with a 3-fold molar excess over Gal3. For antibodies, unreacted BG was eliminated by overnight dialysis against PBS using 10 kDa cut-off dialysis cassettes. For Gal3, unreacted BG was removed using 7 kDa cut-off Zeba spin desalting columns. For fluorescence microscopy analysis of retrograde trafficking, mAb13-BG and Gal3-BG were further incubated for 1 h at 21 °C with a 3- fold molar excess of Cy3-NHS. Unreacted Cy3 was removed using 7 kDa cut-off Zeba spin desalting columns.

### Biochemical retrograde trafficking analysis

HeLa cells stably expressing GalT-GFP-SNAP were continuously incubated for 3 h at 37 °C with 10 μg/mL mAb13-BG or 200 nM Gal3-BG conjugates. Excess of antibody or Gal3 was removed by washing with DMEM, and the unreacted GalT-GFP-SNAP was quenched for 20 min at 37 °C with 10 μM SNAP cell®-Block. Cells were lysed for 30 min at 4 °C with lysis buffer, and cleared lysates were loaded on 30 μl slurry bed G-sepharose beads for overnight pull-down at 4 °C on a rotating wheel. Samples were washed 3- times with TNE 0.1% NP40 buffer. Proteins were eluted by boiling in SDS sample buffer, loaded on denaturing non-reducing SDS-PAGE gels, and immunoblotted with anti-SNAP antibody. When mAb13-BG or Gal3-BG reached the Golgi compartment, a covalent reaction with the SNAPtag occurred, resulting in the formation of mAb13 or Gal3/GalT-GFP-SNAP protein species that were detected by western blotting (anti-SNAP immunoblots) and termed SNAP-mAb13 and SNAP- Gal3, respectively.

### Retrograde trafficking analysis by fluorescence microscopy

10 μg/mL Cy3-mAb13-BG was continuously incubated for 1 h at 37 °C with GalT-GFP-SNAP expressing HeLa cells. Cells were then prepared for fluorescence imaging by confocal microscopy. For investigation of clathrin-dependent endocytosis contribution to retrograde trafficking, 200 nM Cy3-Gal3-BG or 10 μg/mL Cy3-mAb13-BG were incubated as described above, either under siCtl or siCHC depletion conditions.

### mAb13/9EG7 colocalization with Gal3 and retromer complex

Gal3 and mAb13/9EG7 were sequentially bound at 4 °C onto RPE-1 cells, as described above. Cells were then shifted for 15 min to 37 °C, placed at 4 °C, and successively washed with β-D-lactose and acid buffer (see co- binding/co-uptake assays section) to remove cell surface accessible ligands. After PFA fixation, cells were permeabilized and further immunolabeled with anti-VPS26 antibody (see immunofluorescence section).

### mAb13/9EG7 interaction with the retromer complex

10 μg/mL mAb13 or 9EG7 antibodies were continuously incubated for 20 min with RPE-1 cells. Cells were lysed for 30 min at 4 °C in lysis buffer, and the post-nuclear supernatants were applied onto 30 μl slurry of pre-washed G-sepharose beads for an overnight incubation at 4 °C. Proteins were eluted from beads and denatured by boiling in SDS-sample buffer, loaded onto SDS-PAGE gels, and then immunoblotted with anti- VPS35 antibody, or directly with anti-rat IgG-HRP antibody for the detection of mAb13 or 9EG7.

### Purification of α_5_β_1_-integrin from rat livers

Micellar α_5_β_1_ integrin was solubilized and purified as described previously (Dransart et al., 2022). Protein purity was analyzed by stain-free SDS-PAGE. α_5_β_1_ integrin-enriched fractions were then pooled, and final concentration determined by Bradford colorimetric assay. α_5_β_1_ integrin was snap-frozen and stored at -80 °C.

### Negative staining EM of purified α_5_β_1_ integrin

α_5_β_1_ integrin in micelles or reconstituted in lipid nanodiscs or peptidiscs, as described below, was incubated at 50 μg/mL for 30 s on freshly glow-discharged carbon-coated EM grids before staining with 2% of uranyl acetate. Micrographs were recorded at 80 kV (TEM 900 or Tecnai Spirit) or 120 kV (Talos L120C).

### Sialidase treatment of purified α_5_β_1_ integrin

When specified, purified α_5_β_1_ integrin was treated with sialidase (neuraminidase from *Arthrobacter ureafaciens*) at a ratio of 0.08 U of enzyme for 100 μg of α_5_β_1_ integrin to remove terminal sialic acids from integrin glycans.

### Gal3 interaction with α_5_β_1_ integrin in microcavity array suspended lipid bilayers

α_5_β_1_ integrin was reconstituted into microcavity array suspended lipid bilayers at a lipid to protein ratio of 10:1. The microcavity array suspended lipid bilayers were prepared at gold or PDMS polymer substrates for electrochemical impedance spectroscopy (EIS) or FLIM (FCS), respectively, according to procedures described previously (Sarangi# et al., 2022). Membrane capacitance and FLIM measurements were then performed in the presence of increasing concentrations of Gal3. Both FLIM and EIS studies were conducted with microcavity array suspended lipid bilayers filled with and in contact with 10 mM HEPES buffer. Integrin activation was accomplished through sequential addition of 5 mM Mn^2+^, with 30 min incubation, then 1 mM cRGD with 90 min incubation, to the contacting solution. Addition of Gal3 at the indicated concentrations was followed by an equilibration time of 30 min. These times were confirmed to be sufficient for protein binding/equilibration in all cases. All measurements were carried out at room temperature (22±1 °C) and in triplicate. Capacitance values were extracted from EIS data by fit to an equivalent circuit model (ECM) reported previously and analyzed using Z-View software (Scribner Associates, v3.4e). Fits were assessed from both, visual inspection of the fit residuals, and from χ2 (typically ∼ 0.001) As absolute membrane resistance and capacitance can vary with substrate, the average relative changes to membrane resistance (Δ*R*) and capacitance (Δ*Q*) are reported as 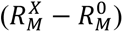 and 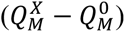; where 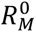 and 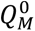 represents respectively the absolute membrane resistance and capacitance in the absence, and 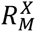 and 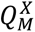 are the respective values of membrane resistance and capacitance values in presence of lectin. For FLIM and FCS measurements, α_5_β_1_ integrin and Gal3 were fluorescently labeled with ATTO488 and Alexa647, respectively. Out of 20 wt% of total integrin, 5 wt% was fluorescently labeled with ATTO488. FLIM/FCS measurements were carried out in triplicate at 20 °C. Data collection, analysis, and extraction of D values were conducted as described previously (Sarangi# et al., 2022).

### Direct Gal3 interaction with α_5_β_1_ integrin conformers in Triton X-100 micelles

The shift from the inactive bent-closed α_5_β_1_ integrin to the active extended ligand-bound conformational state was achieved as follows: 0.5 μg α_5_β_1_ integrin was sialidase-treated as described above, and further incubated for 1 h at 21 °C under gentle agitation at a final concentration of 10 μg/mL in 45 μl HEPES/Triton X-100 buffer, supplemented with 5 mM MnCl_2_. cRGD peptide was then added at a final concentration of 100 μM, and incubated overnight at 4 °C under gentle agitation. This freshly activated α_5_β_1_ integrin, or the inactive bent-closed one, were then incubated for 1 h at 21 °C under gentle agitation with 200 nM of Alexa488-labeled Gal3 (Alexa488-Gal3). Samples were chilled for 10 min on ice, and α_5_β_1_ integrin-Alexa488-Gal3 complexes were cross-linked for 20 min at 4 °C with 2.5 mM glutaraldehyde. Reactions were stopped by incubations for 20 min at 4 °C with 20 mM Tris pH 7.4. Samples were supplemented with 3x non-reducing SDS sample buffer (2% SDS), loaded on SDS-PAGE gels, and run at 100 V without SDS (semi-native conditions). Alexa488-Gal3 signal was directly detected on gels (Alexa488 channel). Proteins were then transferred onto nitrocellulose membranes and immunoblotted for rat β_1_ integrin.

### α_5_β_1_ integrin reconstitution in nanodiscs

Reconstitutions were performed at a R1=Lipid/MSP1D1 ratio of 50 mol/mol, and a R2=α_5_β_1_ integrin/nanodisc ratio of 1 mol/mol, as described previously (Dransart et al., 2022). For one reconstitution reaction (50 μl final volume), His-tagged MSP1D1 scaffold protein was diluted to 4 μM (100 μg/mL) in HEPES buffer supplemented with 0.06% Triton X-100 and incubated for 10 min at 21 °C under gentle stirring. ePC/bPS (90/10, w/w) lipid mix was added at a final concentration of 90 μg/mL, still under stirring, and further incubated for 10 min at 21 °C under gentle stirring. α_5_β_1_ integrin in HEPES/Triton X- 100 buffer was added at a final concentration of 0.25 mg/mL, and incubated for 20 min at 4 °C under gentle stirring. Triton X-100 detergent was finally removed by incubation under gentle stirring for 30 min at 4 °C with 6 mM of Heptakis (2,6-di-O-methyl)-β-cyclodextrin.

### Direct Gal3 interaction with α_5_β_1_ integrin conformers in nanodiscs

For *in vitro* activation, α_5_β_1_ integrin in nanodiscs was first incubated for 30 min at 21 °C with 5 mM MnCl_2_, followed by an additional 2 h incubation at 21 °C with 10 μM FNIII_9-10_ fibronectin fragment. This freshly activated α_5_β_1_ integrin, or the untreated inactive bent-closed one, were adjusted to 500 μl HEPES buffer and incubated with 200 nM Cy3-Gal3-His for 15 min at 18 °C on rotating wheel. The mixture was then incubated for 1 h at 4 °C with 30 μl bed volume of cobalt beads. Beads were washed 3 times with 500 μl HEPES buffer, and α_5_β_1_ integrin-Gal3 complexes were eluted by incubation for 1 h at 21 °C with 40 μl of HEPES buffer supplemented with 10 mM EDTA. Samples were denatured with 3x concentrated non-reducing SDS-sample buffer, boiled for 5 min at 95 °C, and run on SDS-PAGE. α_5_β_1_ integrin and Gal3 were detected using the stain-free and fluorescence (Cy3 channel) modes, respectively.

### Glycan-dependent binding of Gal3 to purified α_5_β_1_ integrin

Gal3 was pre-incubated or not for 10 min at room temperature with either 50 mM β-D-lactose or 100 μM I3 inhibitor. Binding to α_5_β_1_ integrin in Triton X-100 micelles or reconstituted in nanodiscs was then performed as described above.

### Elution of Gal3 or Gal3ΔNter from α_5_β_1_ integrin in nanodiscs

4 reconstitution reactions as described above were incubated overnight at 4 °C on rotating wheel with 30 μl cobalt bead bed volume in 500 μl HEPES buffer. Beads were washed 3 times with 500 μl HEPES buffer. The cobalt bead-immobilized α_5_β_1_ integrin in nanodiscs was incubated in 500 μl HEPES buffer for 2 h at 21 °C on a rotating wheel, either with 4 μM of Gal3 or Gal3ΔNter. Beads were washed 3 times with 500 μl HEPES buffer. Gal3/Gal3ΔNter were specifically eluted with 40 μl of 50 μM I3 inhibitor diluted in HEPES buffer. Elution was performed by gentle manual shaking for 10 min at room temperature, and samples were immediately negatively stained for EM. For the oligomer stability study, eluted samples were kept for the indicated times at 4 °C until loading onto EM grids.

### α_5_β_1_ integrin reconstitution in peptidiscs

For one reconstitution reaction, 10 μg of purified α_5_β_1_ integrin were incubated for 20 min at 21 °C at a final concentration of 0.5 mg/mL with 4.5 μg of peptidisc/peptidisc-His mixture (1:1, w/w). Triton X-100 detergent was removed by the addition of 15 mM of Heptakis (2,6-di-O-methyl)-β-cyclodextrin. 4 reconstitution reactions were pooled in 500 μl HEPES buffer, mixed with 30 μl bed volume cobalt beads and incubated overnight at 4 °C on a rotating wheel. Beads were then washed 3 times with 500 μl of HEPES buffer. Peptidisc-reconstituted α_5_β_1_ integrin was finally eluted by incubation for 20 min at 21 °C under shaking in 40 μl of the same buffer supplemented with 10 mM EDTA. Samples were further analyzed by Blue Native PAGE and EM to qualitatively assess mono-insertion of α_5_β_1_ integrin heterodimers in peptidiscs.

### α_5_β_1_ integrin activation in peptidiscs

α_5_β_1_ integrin in peptidisc (2 reconstitution reactions as above) was incubated for 30 min at 21 °C with 5 mM MnCl_2_, followed by an additional incubation for 2 h at 21 °C with 100 μM cRGD peptide. Samples were then incubated for 2 h at 4 °C on a rotating wheel with 30 μl bed volume of cobalt beads. Beads were washed 3 times with HEPES buffer, supplemented with 5 mM MnCl_2_, and integrin eluted for 20 min at 21 °C with 250 mM imidazole HEPES solution, 5 mM MnCl_2_. Imidazole was removed using 7 kDa cut-off Zeba spin desalting columns equilibrated with HEPES buffer, 5 mM MnCl_2_. Efficient switch from the inactive bent-closed conformation to the active extended ligand-bound one was monitored by negative staining EM and Blue Native PAGE.

### Preparation of α_5_β_1_ integrin-Gal3 complexes in peptidiscs for photobleaching experiments

α_5_β_1_ integrin was reconstituted into peptidiscs as described above, except that a peptidisc/peptidisc- His/peptidisc-Biotin (1/1/1, w/w/w) mixture was used. After overnight incubation with cobalt beads, immobilized peptidiscs were resuspended in 500 μl of HEPES buffer and incubated with sialidase on a rotating wheel according to conditions described above. Beads were washed 3 times with 500 μl of HEPES buffer and further incubated for 2 h at 21 °C on a rotating wheel with 4 μM of Gal3-Cy3. Beads were washed 3 times with 500 μl of HEPES buffer, α_5_β_1_ integrin-Gal3-Cy3 complexes were acutely eluted for 10 min with 40 μl of HEPES buffer supplemented with 10 mM EDTA with manual and gentle resuspension of beads, and immediately analyzed in photobleaching experiments.

### Photobleaching analysis of α_5_β_1_ integrin-Gal3 complexes in peptidiscs

Glass coverslips (Menzel-Gläser, thickness #1) were first washed for 20 min under sonication with chloroform, followed by 5 min washes with water. The coverslips were then sonicated for 20 min with 1 M KOH buffer, followed by 3 times rinsing with water, and final sonication for 20 min with water. The clean coverslips were then dried under a gentle nitrogen stream, and plasma cleaned for 1 min. A double-sided tape mask was used to create a micro chamber for sample incubation to generate surface immobilized peptidisc samples for fluorescence imaging (Damm, 2019). To immobilize the freshly prepared α_5_β_1_ integrin-Gal3 complexes, chambers were first incubated for 1 h with silane-PEG2000 mixed with 1.5% of silane-PEG3400-biotin (LaysanBio) at 5 mM concentration, for 10 min with neutravidin solution at 20 µg/mL, and for 15 min with β-casine at 0.5 mg/mL. The chambers were thoroughly rinsed at each incubation step by injecting 100 µL of buffer. Finally, the α_5_β_1_ integrin-Gal3 complexes in peptidiscs were incubated for 10 min in the chamber and were imaged under glucose oxidase oxygen scavenger solution. For photobleaching experiments, a custom built TIRF Microscope (with 100 x objective, 1.45 NA) was used to excite Gal3-Cy3 with a 532 nm laser. Fluorescence images were recorded with 50 ms exposure time using an ORCA- Flash4.0 V3 Digital CMOS camera (Hamamatsu). All experiments were performed at 21 °C. The recorded fluorescence stream was processed using Matlab-based open-source iSMS and AutoStepfinder software (Loeff et al., 2021; Preus et al., 2015). α_5_β_1_ integrin reconstituted in biotin-free peptidiscs as well as Gal3-Cy3 alone were used as controls.

### Gal3 oligomer elution from RPE-1 cells

RPE-1 cells were cooled for 10 min on ice and then washed 3 times for 5 min with pre-chilled 150 mM β-D-lactose solution to remove endogenous surface-bound Gal3. Cells were then extensively washed with ice-cold PBS^++^ and further incubated for 30 min at 4 °C with 50 μg/mL exogenous Gal3 diluted in serum-free DMEM F12. Unbound Gal3 was removed by 3 washes with PBS^++^. Surface-bound Gal3 was eluted for 30 min at 4 °C with 20 μM of Gal3 inhibitor compound I3. A part of the samples was analyzed by negative staining cryo-EM. Another part was desalted two times with 7 kDa cut-off spin-columns to remove the I3 compound, and processed as described below. Of note, for binding and uptake assays with eluted Gal3 oligomers, Cy3-labeled Gal3 was used.

### Binding and uptake of cell-eluted Gal3 oligomers

RPE-1 cells were incubated (or not) for 24 h at 37 °C with 5000 unit/mL of PNGase F in DMEM F12 supplemented with 1% FCS, to remove N-linked glycans on membrane glycoproteins. For binding assays, cells were cooled for 10 min on ice. 5 μg/mL of Gal3-Cy3 oligomers from cells (see above) or monomeric Gal3-Cy3 were incubated for 10 min at 4 °C with PNGase F-treated or untreated RPE-1 cells. These cells were then washed with cold PBS^++^, immediately fixed with 4% PFA and analyzed by immunofluorescence. For uptake assay, cells were continuously incubated for 2 min at 37 °C. Cells were then placed on ice and membrane-bound fraction of Gal3-Cy3 was removed by β-D-lactose wash. Cells were finally washed with cold PBS^++^, and immediately fixed with 4% PFA. In some conditions, these experiments were performed on RPE-1 cells from which GSLs had also been depleted using Genz-123346 treatment.

### Negative stain EM of Gal3 oligomers

Gal3 oligomers eluted from α_5_β_1_ integrin-Gal3 complexes in nanodiscs or from RPE-1 cells (1:3 dilution in PBS) were incubated for 3 min on freshly glow-discharged carbon-coated EM grids before staining with 2% of uranyl acetate. 20 μg/mL of Gal3 alone (monomer) was used as control. Micrographs were recorded on a Talos L120C TEM equipped with a Ceta16M CCD detector at a pixel size of 1.58 Å per pixel in low-dose mode.

### Data processing of Gal3 oligomers

Gal3 oligomers eluted from RPE-1 cells. 83 negative stain micrographs were imported in cryoSPARC (Punjani et al., 2017). An initial set of 16,640 particles were picked using the blob picker, followed by two rounds of 2D classification to generate templates for autopicking. Template-based autopicking then identified 29,789 particles, which were extracted at a pixel size of 3.16 Å per pixel and subjected to 2D classification. The obtained 2D classes were inspected and sorted in Gal3 dimers, trimers and tetramers. A 3D model was created in cryoSPARC using particles that resembled regular annular tetramers with C4 symmetry applied. Four Gal3 CRDs (PDB 1KJL) (Sörme et al., 2005) were rigid-body fitted into the four corners of the annular density map using UCSF Chimera (Pettersen et al., 2004).

#### Auto-picking method for Gal3 oligomers eluted from nanodics-embedded α_5_β_1_ integrin-Gal3 complexes

Monomers and oligomers on negative stain EM micrographs of Gal3 eluted from inactive and active α_5_β_1_ integrin were automatically picked and quantified using crYOLO (Wagner et al., 2019). First, Gal3 monomers and oligomers, distinguishable by their different size, were manually picked on 11 micrographs of inactive α_5_β_1_ integrin and used to train two autopicking models. 100 and 145 negative stain EM micrographs of Gal3 eluted from inactive and active α_5_β_1_ integrin, respectively, were then automatically picked with the two models.

#### Manual method for Gal3 oligomers eluted from nanodics-embedded α_5_β_1_ integrin-Gal3 complexes

Auto-quantification may include irrelevant objects that resemble to Gal3 oligomers, but which don’t have the defined ring-shaped structure. For example, free nanodiscs can erroneously be selected as Gal3 oligomers. We therefore also provided additional quantification by manually counting of structures that rigorously meet our defined criteria. From EM micrographs of 1 x 0.665 μm size, 4 smaller fields of 0.4 μm x 0.25 μm were selected to facilitate the counting of individual objects. Percentages of defined oligomers and monomers per rectangle were quantified.

### Cryo-EM of neuraminidase-treated α_5_β_1_ integrin

Peptidisc-embedded and neuraminidase-treated α_5_β_1_ integrin was vitrified on Quantifoil 2/1 Cu 300 mesh grids using a Vitrobot Mark IV set to a blot force of -1, blotting time of 3.0 s, 100% humidity, and temperature of 12 °C. 2905 micrographs were acquired using a FEI Titan Krios G3i microscope (Thermo Fisher Scientific) operated at 300 kV equipped with a FEI Falcon 3EC detector (Thermo Fisher Scientific) running in counting mode at a nominal magnification of 96,000×, giving a calibrated pixel size of 0.832 Å/px. Movies were recorded for 40.78 s accumulating a total electron dose of 42 e^−^/Å^2^ fractionated into 33 frames. EPU 2.8 was utilized for automated data acquisition with AFIS enabled using a nominal defocus between -0.8 and -2 µm.

### Data processing of peptidisc-embedded α_5_β_1_ integrin

Data processing is outlined in Figure S5B. 2905 movies were aligned in MotionCor2 (Zheng et al., 2017) and imported in cryoSPARC for subsequent patch CTF estimation. After sorting out bad images, 2884 micrographs were chosen for further processing. 45,681 particles were selected and extracted using the blob picker for initial 2D classification and generation of autopicking templates. Subsequent template-based autopicking identified 2,005,236 particles. After particle curation, 1,444,502 particles were extracted (2x binning) for 2D classification. 470,092 particles in integrin-shaped 2D classes were re-extracted without binning, an initial model was generated from 66,630 particles, and all particles were processed in a further round of 2D classification, after which 432,656 particles were selected. 3D refinements of these particles resulted in anisotropic density maps because of preferred orientation, therefore “Rebalance 2D classes” was executed with two different rebalance factors, resulting in 320,816 and 277,162 remaining particles. These were subjected to heterogeneous 3D refinement with three classes each. In both cases, the 3D class with the least pronounced preferred particle orientation showed also the most well-defined head and upper leg parts. Particles from these two classes were combined, duplicates removed (101,741 remained), and subjected to homogeneous refinement, followed by non-uniform refinement. The headpiece of the density (3.8 Å overall resolution) was well-resolved, whereas the leg piece was fragmented. Therefore, we subtracted the density of the leg piece and carried out local refinement of the remaining headpiece, followed by local CTF refinement. The resolution of the headpiece density map was 3.7 Å according to the gold-standard Fourier shell correlation (FSC) criterion (Figure S5B,E,G). DeepEMhancer (Sanchez-Garcia et al., 2021) was applied for map sharpening. For the leg piece, homogeneous refinement of 277,162 particles after “Rebalance 2D classes” yielded a density map (4.3 Å resolution) in which headpiece and leg piece were equally pronounced. Here, we subtracted the signal of the headpiece and carried out local refinement of the remaining leg piece, yielding a resolution of 4.7 Å according to the gold-standard FSC criterion (Figure S5B,F,H). This map was filtered according to local resolution. Finally, the individually processed maps of headpiece and leg piece were fitted into the 4.3 Å global map of α_5_β_1_ integrin.

### Atomic modeling of α_5_β_1_ integrin

A previously generated homology model of the extracellular domain of rat α_5_β_1_ integrin (residues 94 - 1041 and 23 - 719, respectively) (Mirgorodskaya et al., 2022) based on human α_5_β_1_ integrin (PDB 7NXD) (Schumacher et al., 2021) was used as starting model. The homology model was split into headpiece and leg piece (α_5_, residues 94 - 691 and 692 - 1041; β_1_, residues 23 - 504 and 505 - 720, respectively) and rigid-body fitted into the sharpened density maps of headpiece and leg piece using UCSF Chimera (Pettersen et al., 2004), followed by flexible fitting with imodfit (Lopez-Blanco and Chacon, 2013). The model of the headpiece was then manually adjusted in Coot (Emsley et al., 2010) and ISOLDE (Croll, 2018), and refined by real-space refinement in Phenix (Afonine et al., 2018) in an iterative manner (Figure S5K,L). Cryo-EM data processing and model refinement statistics are summarized in Table S1.

### Glycan attachment to atomic model with GlycoShield

Glycans were added to the glycosylation sites in α_5_β_1_ integrin using the reductionist molecular dynamics simulation method described in GlycoSHIELD (Gecht et al., BioRxiv). Glycans were added as identified in (Mirgorodskaya et al., 2022) (α_5_ integrin, biantennary at N136, N231, N356, N642, N761, N773, N918, triantennary at N822, hybrid at N346, high mannose at N365, N657, N724; β_1_ integrin, complex biantennary at N50, N97, N212, N269, N363, N406, N417, N482, N521), and models were obtained in coarse grain mode. 100 conformations for every glycan were then combined on the α_5_β_1_ integrin model.

### Preparation of neuraminidase-treated α_5_β_1_ integrin-Gal3 complexes for cryo-EM

α_5_β_1_ integrin was reconstituted in peptidiscs as described above and immobilized onto cobalt beads by overnight incubation. Bead-immobilized peptidiscs were resuspended in 500 μl of HEPES buffer and incubated with sialidase on a rotating wheel according to conditions described earlier. Beads were washed 3 times with 500 μl of HEPES buffer and further incubated for 2 h at 21 °C on a rotating wheel with 100 μg/mL of Gal3. Beads were washed 3 times with 500 μl of HEPES buffer and α_5_β_1_ integrin-Gal3 complexes were rapidly eluted for 10 min with 40 μl of HEPES buffer supplemented with 10 mM EDTA, with manual and gentle resuspension of beads. Eluted complexes were immediately used for subsequent negative staining and cryo-EM analysis.

### Negative stain EM of α_5_β_1_ integrin alone or in complex with Gal3

Peptidisc-embedded α_5_β_1_ integrin alone or in complex with Gal3 was diluted 1:10 after elution using 20 mM HEPES buffer and applied on freshly glow-discharged carbon-coated EM grids before staining with 2% of uranyl acetate. Micrographs were recorded at 80 kV on a TEM 900 (Zeiss) equipped with a Morada G2 camera (Olympus) or a Tecnai Spirit equipped with QUEMESA camera.

### Cryo-EM of neuraminidase-treated α_5_β_1_ integrin-Gal3 complexes

Immediately after elution from cobalt beads, samples were vitrified on Quantifoil 2/1 Cu 300 mesh grids using a Vitrobot Mark IV set to a blot force of -1, blotting time of 3.5 s, 100% humidity, and temperature of 12 °C. Micrographs were acquired using a FEI Titan Krios G3i microscope (Thermo Fisher Scientific) operated at 300 kV equipped with a Bioquantum K3 direct electron detector and energy filter (Gatan) running in CDS super-resolution mode at a slit width of 20 eV and at a nominal magnification of 81,000×, giving a calibrated physical pixel size of 1.06 Å/px. Movies were recorded for 3.0 s accumulating a total electron dose of 61 e^−^/Å^2^ fractionated into 60 frames. EPU 2.12 was utilized for automated data acquisition with AFIS enabled using a nominal defocus between -1.0 and -2.8 µm. Two data sets (6,028 and 10,325 movie images, respectively) were recorded under identical conditions and merged after the initial processing steps.

### Data processing of α_5_β_1_ integrin-Gal3 complexes

Drift correction and estimation of CTF parameters of movies of both data sets were carried out in cryoSPARC (version 3.3.2) using the built-in patch motion correction and patch CTF estimation algorithms. After sorting out bad images, 5,614 and 10,225 micrographs remained from the data sets for subsequent processing steps, respectively. With the blob picker on 1,000 images of the first data set and subsequent 2D classification in cryoSPARC, autopicking templates were generated and used to pick 1,327,764 and 3,894,571 particles from the data sets, respectively. These were subjected to two subsequent rounds of 2D classification with 8x and 4x binning (pixel sizes of 4.24 and 2.12 Å/pixel, respectively), leaving 289,613 and 556,786 integrin-shaped particles, respectively. These were combined for the subsequent processing steps. Non-uniform refinement (Punjani et al., 2020) in cryoSPARC with 2x binned data (1.06 Å/pixel) using an initial model generated from the second data set resulted in a density map with 4.1 Å overall resolution, according to the gold-standard FSC criterion. Different 3D sorting approaches in cryoSPARC (version 3.3 and 4.0.3) and Relion (version 4.0) (Scheres, 2012) were applied, with the best final results achieved by 3D classification in cryoSPARC (version 4.03) in combination with starting models generated by previous 3D classifications in Relion and a focused mask on the space between headpiece and leg piece. Transfer of particles between cryoSPARC and Relion was performed by pyem (DOI: 10.5281/zenodo.3576630). Two rounds of 3D classification were carried out with four classes each, the first round in input mode, the second round in simple mode. For the first round, the 3D map originating from all particles (see above) and three different 3D maps originating from Relion 3D classifications served as input models, lowpass filtered to 20 Å resolution. 387,525 particles from the two classes with most pronounced densities outside α_5_β_1_ integrin were transferred to a second round of 3D classification in simple mode. This resulted in two classes with 111,868 and 112,569 particles, respectively, that showed more density outside α_5_β_1_ integrin than the others. The class with 111,869 particles (termed subset 1) that has the most pronounced extra density was then refined (non-uniform refinement), resulting in a final resolution of 6.9 Å, according to the gold-standard FSC criterion.

### Data availability

Cryo-EM density maps of α_5_β_1_ integrin and the α_5_β_1_ integrin-Gal3 complex have been deposited to the EMDB under accession codes 17269 (head piece), 17270 (leg piece), 17271 (complex), respectively, and in the pdb under accession code 8OXZ (headpiece).

### Micellar α_5_β_1_ integrin-Gal3 complex cross-linking and mass spectrometry analysis

α_5_β_1_ integrin at 40 μg/mL and His-tagged Gal3 (Gal3-His) at 400 nM in 500 μl HEPES/Triton X-100 buffer were co-incubated for 20 min on a rotating wheel. Samples were chilled for 10 min on ice, and α_5_β_1_ integrin-Gal3-His complexes cross-linked for 2 h at 4 °C using a 40 μM BS3 d0/d4 mixture (20 μM each), or 100 μM DSBU. Excess cross-linkers were quenched for 20 min on ice with 20 mM Tris pH 7.4. Gal3 and excess cross-linkers were removed by loading samples on 40 kDa cut- off 2 ml desalting spin-columns equilibrated with HEPES/Triton X-100 buffer. Two reactions as described above were pooled and incubated with 25 μl bed volume of cobalt beads prepared according to manufacturer instructions. Beads were washed 3 times with 500 μl HEPES/Triton X-100 buffer. After the last wash, beads were resuspended in HEPES/Triton X-100 buffer, supplemented with deglycosylation denaturing buffer and boiled for 5 min at 95 °C. Samples were chilled for 10 min on ice and supplemented with 1% (v/v) NP40. Deglycosylation was achieved upon addition of 2 μl of PNGase F and incubation for 2 h at 37 °C. Reactions were stopped by the addition of 3x non-reducing SDS sample buffer and boiling for 5 min at 95 °C. For biochemical characterization of cross-linked complexes, samples were loaded on SDS-PAGE and proteins were detected in the stain-free mode. For mass spectrometry, samples were run on SDS-PAGE at 100 V until the migration front reached half of the 4% acrylamide stacking gel. Proteins were fixed by incubating gels for 30 min at room temperature in ethanol/acetic acid 50/3 (v/v) solution. After 3 washes with ultra-pure water, gels were stained for 1 h at room temperature with Coomassie blue solution. Stained protein bands that remained after washing in water were cut, washed with 25 mM NH_4_HCO_3_ before reduction and alkylation with 10 mM DTT and 55 mM iodoacetamide. In-gel digestion was performed overnight with trypsin. Peptides were recovered and injected on a nanoLC-MS/MS system. XL-BSA was used as a quality control. NanoLC-MS/MS analyses were performed with a nanoAcquity UPLC device coupled to a Q-Exactive HF-X mass spectrometer. Peptide separation was performed on an Acquity UPLC BEH130 C18 column (250 mm × 75 μm with 1.7-μm-diameter particles) and a Symmetry C18 precolumn (20 mm × 180 μm with 5-μm- diameter particles, Waters). The solvent system consisted of 0.1% formic acid (FA) in water and 0.1% FA in acetonitrile (ACN). The system was operated in data dependent acquisition mode with automatic switching between MS and MS/MS modes. The ten most abundant ions were selected on each MS spectrum for further isolation. HCD fragmentation method was used with different collision energy (NCE 27, 30 and 33). The dynamic exclusion time was set to 60 s. Raw data were processed and converted into *.mgf format. The MS/MS data were analyzed using MeroX software v2.0.1.4 (Götze et al., 2015). Mass tolerances of 5 ppm for precursor ions and of 10 ppm for product ions were applied. A 5% FDR cut-off and a signal-to-noise > 2 were applied. Lys and Arg residues were considered as protease sites with a maximum of three missed cleavages. Carbamidomethylation of cysteine was set as static modification and oxidation of methionine as variable modification (max. mod. 2). Primary amino groups (Lys side chains and N-termini) as well as primary hydroxyl groups (Ser, Thr and Tyr side chains) were considered as cross-linking sites. An in-house database comprising rat α_5_ and β_1_ integrin, human galectin-3 and bovine serum albumin was used. Cross-links composed of consecutive amino acid sequences were ignored. Each cross-linked product automatically annotated with MeroX was manually validated. Two biological replicates with technical triplicate were made for DSBU experiment (total of 6 samples). For BS3, three different experiments were performed. Cross-linked peptides were validated when seen in 2 out of 6 experiments and 2 out of 3 experiments for DSBU and BS3, respectively. The XL-MS data set has been deposited to the ProteomeXchange Consortium via the PRIDE partner repository with the dataset identifier PXD041522 (Perez-Riverol et al., 2022).

### Gal3 placement on α_5_β_1_ integrin

For fitting Gal3 into the map of the α_5_β_1_ integrin-Gal3 complex, the headpiece and leg piece maps of α_5_β_1_ integrin including the fitted atomic models were fitted into the respective domains of the complex map first. Subsequently, Gal3 CRDs (PDB 1KJL) (Sörme et al., 2005) were placed into the density between headpiece and leg piece. This density is compatible to fit two Gal3 CRDs that interacted with glycans at N356 and N918, or at N773 and N918 of α_5_ integrin.. Gal3 CRDs were manually placed in density outside the α_5_β_1_ integrin model such that the following requirements were fulfilled: (i) match of galactose in the Gal3 PDB with terminal galactose of glycan chains (N356 and N918, or N773 and N918 of α_5_ integrin), (ii) close spatial proximity of Gal3 K227 with α_5_ K965/K1028 (Gal3 contacting glycan at α_5_ N918), (iii) N-terminal ends of Gal3 CRDs protruding to the same directions to allow for oligomerization via the N-terminal domain, (iv) no overlap/clashes of Gal3 CRDs with each other and the α_5_β_1_ integrin model. For representation of the glycan distribution on active extended ligand-bound α_5_β_1_ integrin, α_5_ and β_1_ integrin headpiece and leg piece of the glycosylated homology model were rigid-body fitted into the cryo-EM density of the active extended ligand-bound conformation of human α_5_β_1_ integrin (EMDB 12634, working map including the low-resolution parts) (Schumacher et al., 2021). Gal3 CRDs were placed on glycans at N356, N773, and N918 of α_5_ integrin, as identified by cryo-EM, cross-linking proteomics, and site-directed mutagenesis, respectively.

### Gal3-induced locking of α_5_β_1_ integrin in the inactive bent-closed conformational state

Cellular approach: RPE-1 cells were cooled for 10 min on ice and incubated for 30 min at 4 °C in the presence or absence of 200 nM Gal3 or Gal31′Nter. Cells were then washed, and integrins were activated for 1 h at 4 °C with 1 mM MnCl_2_ and 5 μg/mL of soluble fibronectin mixture. The efficiency of activation was monitored by surface immunostaining using mAb13 antibody, whose labeling intensity expectedly decreased in the absence of exogenous Gal3 addition. *in vitro* approach: 100 μg/mL of sialidase-treated α_5_β_1_ integrin in micelles were pre-incubated or not for 3 h at 18 °C with 2 μM of Alexa488-Gal3, and then activated by incubation with 5 mM MnCl_2_ and 100 μM cRGD. Samples were supplemented with 3x non-reducing SDS (2% SDS) sample buffer, loaded on PAGE, and run at 100 V without SDS (semi-native conditions). Alexa488-Gal3 signal was directly detected on gels (Alexa488 channel). Proteins were then transferred onto nitrocellulose membranes and immunoblotted against rat β_1_ integrin.

### Expression of human α_5_β_1_ integrin N-glycosylation mutants in MKF-dKO cells

N/Q substitutions of key N-glycosylation sites of human α_5_ integrin were designed, and both, α_5_ integrin (wildtype or N/Q mutants) and β_1_ integrin (wildtype) cDNAs (GeneScript, 3.5 μg of each) were co-electroporated into MKF-dKO cells. Expression was monitored 24 h post-transfection, using mAb13 and 9EG7 antibodies. mAb13 or 9EG7 antibody uptake and Gal3/mAb13 or 9EG7 co-binding experiments and confocal imaging were performed as described above.

## Acknowledgments

We acknowledge the Cell and Tissue Imaging core facility (PICT IBiSA), Institut Curie, member of the French National Research Infrastructure France-BioImaging (ANR10-INBS-04) for help with microscopy, the Recombinant Protein core facility CNRS-Institut Curie for protein purification, Reinhard Fässler and Jakob Reber (Max Planck Institute of Biochemistry, Munich, Germany) for the MKF-dKO cell line and helpful comments on the manuscript, Jean Salamero for help with LLSM experiments and data analysis, Fredrik Zetterberg (Galecto Biotech Inc.) for I3 compound, and Ewan MacDonald for helpful comments on the manuscript. We also acknowledge access to electron microscopic equipment at the core facility BioSupraMol of Freie Universität Berlin, supported through grants from the Deutsche Forschungsgemeinschaft (DFG) and the state of Berlin for large equipment according to Art. 91b GG (INST 335/588-1 FUGG, INST 335/589- 1 FUGG, INST 335/590-1 FUGG), Tarek Hilal for cryo-EM data collection of α_5_β_1_ integrin, and Thiemo Sprink of the Core Facility for cryo-Electron Microscopy (CFcryoEM) of the Charité - Universitätsmedizin Berlin supported by DFG (INST 335/588-1 FUGG) for cryo-EM data collection of α_5_β_1_ integrin-Gal3. Finally, we acknowledge the use of resources of the French Proteomic Infrastructure ProFI ANR-10-INBS-08–03. This work was supported by grants from Mizutani Foundation reference n° 200014 (LJ), Agence National de la Recherche ANR-19-CE13- 0001-01, ANR-20-CE15-0009-01, ANR-22-CE11-0030-03 (LJ), DALLISH-ANR-16-CE23-0005 (CAV-C, LL, LJ), Fondation pour la Recherche Médicale EQU202103012926 (LJ), LabEx Cell(n)Scale (ANR-11-LABX-0038) as part of the Idex PSL (ANR-10-IDEX-0001-02) (CAV-C, LL, LJ), ITMO Cancer (18CQ091) (CAV-C, LL), Science Foundation Ireland [19/FFP/6428] and [14/IA/2488] (TEK). RR acknowledges funding from the European Union’s Horizon 2020 research and innovation programme under the Marie Skłodowska-Curie grant agreement no: 101025342.

## Author Contributions

LJ, MS-Z, ED, DR are listed for conceptualization, LJ, MS-Z, ED, DR, SR, CW, VC, CAV-C, RR, LL, DL, AH, NKS, JR, UJN for methodology, MS-Z, ED, CAV-C, LL, RR, DL, ADC, AH, NKS, JR, RB for investigation, MS-Z, ED, DR, VC, DL, ADC, RR, AH, NKS, JR for visualization, LJ for funding acquisition, LJ for project administration, LJ, MS-Z, ED, TEK, SCS, SR, DR for supervision, LJ, MS-Z, ED, DR for writing of the original draft, and DL, CW, HL, VC, RR, CAV-C, LL, TEK for review and editing.

## References

Adair, B.D., Xiong, J.P., Yeager, M., and Arnaout, M.A. (2023). Cryo-EM structures of full-length integrin αIIbβ3 in native lipids. Nat Commun 14, 4168.

Afonine, P.V., Poon, B.K., Read, R.J., Sobolev, O.V., Terwilliger, T.C., Urzhumtsev, A., and Adams, P.D. (2018). Real-space refinement in PHENIX for cryo-EM and crystallography. Acta Crystallogr D Struct Biol 74, 531–544.

Aguet, F., Upadhyayula, S., Gaudin, R., Chou, Y.Y., Cocucci, E., He, K., Chen, B.C., Mosaliganti, K., Pasham, M., Skillern, W., et al. (2016). Membrane dynamics of dividing cells imaged by lattice light-sheet microscopy. Mol Biol Cell 27, 3418–3435.

Ahmad, N., Gabius, H.J., Andre, S., Kaltner, H., Sabesan, S., Roy, R., Liu, B., Macaluso, F., and Brewer, C.F. (2004). Galectin-3 precipitates as a pentamer with synthetic multivalent carbohydrates and forms heterogeneous cross-linked complexes. J Biol Chem 279, 10841–10847.

Almeida-Souza, L., Frank, R.A.W., Garcia-Nafria, J., Colussi, A., Gunawardana, N., Johnson, C.M., Yu, M., Howard, G., Andrews, B., Vallis, Y., et al. (2018). A flat BAR protein promotes actin polymerization at the base of clathrin-coated pits. Cell 174, 325–337.e314.

Bazzoni, G., Shih, D.T., Buck, C.A., and Hemler, M.E. (1995). Monoclonal antibody 9EG7 defines a novel beta 1 integrin epitope induced by soluble ligand and manganese, but inhibited by calcium. J Biol Chem 270, 25570–25577.

Benmerah, A., Lamaze, C., Begue, B., Schmid, S.L., Dautry-Varsat, A., and Cerf-Bensussan, N. (1998). AP-2/Eps15 interaction is required for receptor-mediated endocytosis. J Cell Biol 140, 1055–1062.

Birdsall, B., Feeney, J., Burdett, I.D., Bawumia, S., Barboni, E.A., and Hughes, R.C. (2001). NMR solution studies of hamster galectin-3 and electron microscopic visualization of surface-adsorbed complexes: evidence for interactions between the N- and C-terminal domains. Biochemistry 40, 4859–4866.

Bojić-Trbojević, Ž., Jovanović Krivokuća, M., Vilotić, A., Kolundžić, N., Stefanoska, I., Zetterberg, F., Nilsson, U.J., Leffler, H., and Vićovac, L. (2019). Human trophoblast requires galectin-3 for cell migration and invasion. Sci Rep 9, 2136.

Boscher, C., and Nabi, I.R. (2013). Galectin-3- and phospho-caveolin-1-dependent outside-in integrin signaling mediates the EGF motogenic response in mammary cancer cells. Mol Biol Cell 24, 2134–2145.

Boucrot, E., Ferreira, A.P., Almeida-Souza, L., Debard, S., Vallis, Y., Howard, G., Bertot, L., Sauvonnet, N., and McMahon, H.T. (2015). Endophilin marks and controls a clathrin-independent endocytic pathway. Nature 517, 460–465.

Bridgewater, R.E., Norman, J.C., and Caswell, P.T. (2012). Integrin trafficking at a glance. J Cell Sci 125, 3695–3701.

Caldieri, G., Barbieri, E., Nappo, G., Raimondi, A., Bonora, M., Conte, A., Verhoef, L.G., Confalonieri, S., Malabara, M.G., Bianchi, F., et al. (2017). Reticulon3-dependent ER-PM contact sites control EGFR nonclathrin endocytosis. Science 256, 617–624.

Caswell, P.T., Chan, M., Lindsay, A.J., McCaffrey, M.W., Boettiger, D., and Norman, J.C. (2008). Rab-coupling protein coordinates recycling of alpha5beta1 integrin and EGFR1 to promote cell migration in 3D microenvironments. J Cell Biol 183, 143–155.

Chao, W.T., and Kunz, J. (2009). Focal adhesion disassembly requires clathrin-dependent endocytosis of integrins. FEBS Lett 583, 1337–1343.

Chen, B.C., Legant, W.R., Wang, K., Shao, L., Milkie, D.E., Davidson, M.W., Janetopoulos, C., Wu, X.S., Hammer, J.A., 3rd, Liu, Z., et al. (2014). Lattice light-sheet microscopy: imaging molecules to embryos at high spatiotemporal resolution. Science 346, 1257998.

Chiu, Y.P., Sun, Y.C., Qiu, D.C., Lin, Y.H., Chen, Y.Q., Kuo, J.C., and Huang, J.R. (2020). Liquid-liquid phase separation and extracellular multivalent interactions in the tale of galectin-3. Nat Commun 11, 1229.

Croll, T.I. (2018). ISOLDE: a physically realistic environment for model building into low-resolution electron-density maps. Acta Cryst D74, 519–530.

Damm, A. (2019). Interplay between the conformational dynamics of a transmembrane protein and the mechanical properties of its surrounding membrane. Doctoral dissertation, Sorbonne Université, Paris, France

Dennis, J.W. (2015). Many Light Touches Convey the Message. Trends Biochem Sci 40, 673–686.

Dransart, E., Di Cicco, A., El Marjou, A., Lévy, D., Johansson, S., Johannes, L., and Shafaq-Zadah, M. (2022). Solubilization and purification of alpha5beta1 integrin from rat liver for reconstitution into nanodiscs. Meth Mol Biol 2507, 1–18.

Emsley, P., Lohkamp, B., Scott, W.G., and Cowtan, K. (2010). Features and development of Coot. Acta Crystallogr D Biol Crystallogr 66, 486–501.

Ewers, H., Römer, W., Smith, A.E., Bacia, K., Dmitrieff, S., Chai, W., Mancini, R., Kartenbeck, J., Chambon, V., Berland, L., et al. (2010). GM1 structure determines SV40-induced membrane invagination and infection. Nat Cell Biol 12, 11–18.

Ezratty, E.J., Bertaux, C., Marcantonio, E.E., and Gundersen, G.G. (2009). Clathrin mediates integrin endocytosis for focal adhesion disassembly in migrating cells. J Cell Biol 187, 733–747.

Ezratty, E.J., Partridge, M.A., and Gundersen, G.G. (2005). Microtubule-induced focal adhesion disassembly is mediated by dynamin and focal adhesion kinase. Nat Cell Biol 7, 581–590.

Flores-Ibarra, A., Vertesy, S., Medrano, F.J., Gabius, H.J., and Romero, A. (2018). Crystallization of a human galectin-3 variant with two ordered segments in the shortened N-terminal tail. Sci Rep 8, 9835.

Furtak, V., Hatcher, F., and Ochieng, J. (2001). Galectin-3 mediates the endocytosis of beta-1 integrins by breast carcinoma cells. Biochem Biophys Res Commun 289, 845–850.

Gecht, M., von Bülow, S., Penet, C., Hummer, G., Hanus, C., and Sikora, M. (BioRxiv). GlycoSHIELD: a versatile pipeline to assess glycan impact on protein structures. https://doiorg/101101/20210804455134.

Goetz, J.G., Joshi, B., Lajoie, P., Strugnell, S.S., Scudamore, T., Kojic, L.D., and Nabi, I.R. (2008). Concerted regulation of focal adhesion dynamics by galectin-3 and tyrosine-phosphorylated caveolin-1. J Cell Biol 180, 1261–1275.

Götze, M., Pettelkau, J., Fritzsche, R., Ihling, C.H., Schäfer, M., and Sinz, A. (2015). Automated assignment of MS/MS cleavable cross-links in protein 3D-structure analysis. J Am Soc Mass Spectrom 26, 83–97.

Gu, J., and Taniguchi, N. (2004). Regulation of integrin functions by N-glycans. Glycoconj J 21, 9–15.

Hang, Q., Isaji, T., Hou, S., Wang, Y., Fukuda, T., and Gu, J. (2017). A key regulator of cell adhesion: Identification and characterization of important N-glycosylation sites on integrin alpha5 for cell migration. Mol Cell Biol 37, e00558–00516.

Hopkins, C.R., Miller, K., and Beardmore, J.M. (1985). Receptor-mediated endocytosis of transferrin and epidermal growth factor receptors: a comparison of constitutive and ligand-induced uptake. J Cell Sci Suppl 3, 173–186.

Hou, S., Hang, Q., Isaji, T., Lu, J., Fukuda, T., and Gu, J. (2016). Importance of membrane-proximal N-glycosylation on integrin beta1 in its activation and complex formation. Faseb J 30, 4120–4131.

Isaji, T., Sato, Y., Fukuda, T., and Gu, J. (2009). N-glycosylation of the I-like domain of beta1 integrin is essential for beta1 integrin expression and biological function: identification of the minimal N-glycosylation requirement for alpha5beta1. J Biol Chem 284, 12207–12216.

Isaji, T., Sato, Y., Zhao, Y., Miyoshi, E., Wada, Y., Taniguchi, N., and Gu, J. (2006). N-glycosylation of the beta-propeller domain of the integrin alpha5 subunit is essential for alpha5beta1 heterodimerization, expression on the cell surface, and its biological function. J Biol Chem 281, 33258–33267.

Ivashenka, A., Wunder, C., Chambon, V., Johannes, L., and Shafaq-Zadah, M. (2022). Transcytosis of galectin-3 in mouse intestine. Meth Mol Biol 2442, 367–390.

Jamison, F.W., 2nd, Foster, T.J., Barker, J.A., Hills, R.D., Jr., and Guvench, O. (2011). Mechanism of binding site conformational switching in the CD44-hyaluronan protein-carbohydrate binding interaction. J Mol Biol 406, 631–647.

Jaqaman, K., Loerke, D., Mettlen, M., Kuwata, H., Grinstein, S., Schmid, S.L., and Danuser, G. (2008). Robust single-particle tracking in live-cell time-lapse sequences. Nat Methods 5, 695–702.

Johannes, L., Parton, R.G., Bassereau, P., and Mayor, S. (2015). Building endocytic pits without clathrin. Nat Rev Mol Cell Biol 16, 311–321.

Johannes, L., and Shafaq-Zadah, M. (2013). SNAP-tagging the retrograde route. Methods in Cell Biology 118, 139–155.

Johannes, L., Wunder, C., and Shafaq-Zadah, M. (2016). Glycolipids and lectins in endocytic uptake processes J Mol Biol 428, 4792–4818.

Kabbani, A.M., Raghunathan, K., Lencer, W.I., Kenworthy, A.K., and Kelly, C.V. (2020). Structured clustering of the glycosphingolipid GM1 is required for membrane curvature induced by cholera toxin. Proc Natl Acad Sci USA 117, 14978–14986.

Kaksonen, M., and Roux, A. (2018). Mechanisms of clathrin-mediated endocytosis. Nat Rev Mol Cell Biol 19, 313–326.

Kaplan, M., Narasimhan, S., de Heus, C., Mance, D., van Doorn, S., Houben, K., Popov-Celeketic, D., Damman, R., Katrukha, E.A., Jain, P., et al. (2016). EGFR dynamics change during activation in native membranes as revealed by NMR. Cell 167, 1241–1251.

Kenworthy, A.K., Schmieder, S.S., Raghunathan, K., Tiwari, A., Wang, T., Kelly, C.V., and Lencer, W.I. (2021). Cholera toxin as a probe for membrane biology. Toxins (Basel) 13, 543.

Kirchhausen, T., Owen, D., and Harrison, S.C. (2014). Molecular structure, function, and dynamics of clathrin-mediated membrane traffic. Cold Spring Harb Perspect Biol 6, a016725.

Krzeminski, M., Singh, T., André, S., Lensch, M., Wu, A.M., Bonvin, A.M., and Gabius, H.J. (2011). Human galectin-3 (Mac-2 antigen): defining molecular switches of affinity to natural glycoproteins, structural and dynamic aspects of glycan binding by flexible ligand docking and putative regulatory sequences in the proximal promoter region. Biochim Biophys Acta 1810, 150–161.

Kural, C., Akatay, A.A., Gaudin, R., Chen, B.C., Legant, W.R., Betzig, E., and Kirchhausen, T. (2015). Asymmetric formation of coated pits on dorsal and ventral surfaces at the leading edges of motile cells and on protrusions of immobile cells. Mol Biol Cell 26, 2044–2053.

Kyumurkov, A., Bouin, A.P., Boissan, M., Manet, S., Baschieri, F., Proponnet-Guerault, M., Balland, M., Destaing, O., Régent-Kloeckner, M., Calmel, C., et al. (2023). Force tuning through regulation of clathrin-dependent integrin endocytosis. J Cell Biol 222, e202004025.

Labun, K., Krause, M., Torres Cleuren, Y., and Valen, E. (2021). CRISPR Genome Editing Made Easy Through the CHOPCHOP Website. Curr Protoc 1, e46.

Lagana, A., Goetz, J.G., Cheung, P., Raz, A., Dennis, J.W., and Nabi, I.R. (2006). Galectin binding to Mgat5-modified N-glycans regulates fibronectin matrix remodeling in tumor cells. Mol Cell Biol 26, 3181–3193.

Lakshminarayan, R., Wunder, C., Becken, U., Howes, M.T., Benzing, C., Arumugam, S., Sales, S., Ariotti, N., Chambon, V., Lamaze, C., et al. (2014). Galectin-3 drives glycosphingolipid-dependent biogenesis of clathrin-independent carriers. Nat Cell Biol 16, 595–606.

Lenter, M., Uhlig, H., Hamann, A., Jenö, P., Imhof, B., and Vestweber, D. (1993). A monoclonal antibody against an activation epitope on mouse integrin chain beta 1 blocks adhesion of lymphocytes to the endothelial integrin alpha 6 beta 1. Proc Natl Acad Sci USA 90, 9051–9055.

Lepur, A., Salomonsson, E., Nilsson, U.J., and Leffler, H. (2012). Ligand induced galectin-3 self-association. J Biol Chem 287, 21751–21756.

Lete, M.G., Franconetti, A., Delgado, S., Jiménez-Barbero, J., and Ardá, A. (2022). Oligosaccharide presentation modulates the molecular recognition of glycolipids by galectins on membrane surfaces. Pharmaceuticals (Basel) 15, 145.

Li, J., Su, Y., Xia, W., Qin, Y., Humphries, M.J., Vestweber, D., Cabanas, C., Lu, C., and Springer, T.A. (2017). Conformational equilibria and intrinsic affinities define integrin activation. Embo J 36, 629–645.

Lin, Y.H., Qiu, D.C., Chang, W.H., Yeh, Y.Q., Jeng, U.S., Liu, F.T., and Huang, J.R. (2017). The intrinsically disordered N-terminal domain of galectin-3 dynamically mediates multisite self-association of the protein through fuzzy interactions. J Biol Chem 292, 17845–17856.

Liu, W., Hsu, D.K., Chen, H.Y., Yang, R.Y., Carraway, K.L., 3rd, Isseroff, R.R., and Liu, F.T. (2012). Galectin-3 regulates intracellular trafficking of EGFR through Alix and promotes keratinocyte migration. J Invest Dermatol 132, 2828–2837.

Loeff, L., Kerssemakers, J.W.J., Joo, C., and Dekker, C. (2021). AutoStepfinder: A fast and automated step detection method for single-molecule analysis. Patterns (N Y) 2, 100256.

Lopez-Blanco, J.R., and Chacon, P. (2013). iMODFIT: efficient and robust flexible fitting based on vibrational analysis in internal coordinates. J Struct Biol 184, 261–270.

MacDonald, E., Forrester, A., Valades Cruz, C.A., Madsen, T.D., Hetmanski, J.H.R., Dransart, D., Yeap, N., Godbole, R., Akhil Shp, A., Leconte, L., et al. (submitted). EGF-induced desialylation for the fast control of endocytosis.

Marsico, G., Russo, L., Quondamatteo, F., and Pandit, A. (2018). Glycosylation and Integrin Regulation in Cancer. Trends Cancer 4, 537–552.

Mathew, M.P., and Donaldson, J.G. (2019). Glycosylation and glycan interactions can serve as extracellular machinery facilitating clathrin independent endocytosis. Traffic 20, 295–300.

Mettlen, M., Chen, P.H., Srinivasan, S., Danuser, G., and Schmid, S.L. (2018). Regulation of Clathrin-Mediated Endocytosis. Annu Rev Biochem 87, 871–896.

Mirgorodskaya, E., Dransart, E., Shafaq-Zadah, M., Roderer, D., Sihlbom, C., Leffler, H., and Johannes, L. (2022). Site-specific N-glycan profiles of a5b1 integrin from rat liver. Biol Cell 114, 160–176.

Moreno-Layseca, P., Icha, J., Hamidi, H., and Ivaska, J. (2019). Integrin trafficking in cells and tissues. Nat Cell Biol 21, 122–132.

Moreno-Layseca, P., Jäntti, N.Z., Godbole, R., Sommer, C., Jacquemet, G., Al-Akhrass, H., Conway, J.R.W., Kronqvist, P., Kallionpää, R.E., Oliveira-Ferrer, L., et al. (2021). Cargo-specific recruitment in clathrin- and dynamin-independent endocytosis. Nat Cell Biol.

Mould, A.P., Garratt, A.N., Askari, J.A., Akiyama, S.K., and Humphries, M.J. (1995). Identification of a novel anti-integrin monoclonal antibody that recognises a ligand-induced binding site epitope on the beta 1 subunit. FEBS Lett 363, 118–122.

Nabi, I.R., Shankar, J., and Dennis, J.W. (2015). The galectin lattice at a glance. J Cell Sci 128, 2213–2219.

Nieminen, J., Kuno, A., Hirabayashi, J., and Sato, S. (2007). Visualization of galectin-3 oligomerization on the surface of neutrophils and endothelial cells using fluorescence resonance energy transfer. J Biol Chem 282, 1374–1383.

Pan, D., and Song, Y. (2010). Role of altered sialylation of the I-like domain of beta1 integrin in the binding of fibronectin to beta1 integrin: thermodynamics and conformational analyses. Biophys J 99, 208–217.

Perez-Riverol, Y., Bai, J., Bandla, C., García-Seisdedos, D., Hewapathirana, S., Kamatchinathan, S., Kundu, D.J., Prakash, A., Frericks-Zipper, A., Eisenacher, M., et al. (2022). The PRIDE database resources in 2022: a hub for mass spectrometry-based proteomics evidences. Nucleic Acids Res 50, D543–d552.

Pettersen, E.F., Goddard, T.D., Huang, C.C., Couch, G.S., Greenblatt, D.M., Meng, E.C., and Ferrin, T.E. (2004). UCSF Chimera--a visualization system for exploratory research and analysis. J Comput Chem 25, 1605–1612.

Pezeshkian, W., Shillcock, J.C., and Ipsen, J.H. (2021). Computational approaches to explore bacterial toxin entry into the host cell. Toxins (Basel) 13, 449.

Pike, J.A., Styles, I.B., Rappoport, J.Z., and Heath, J.K. (2017). Quantifying receptor trafficking and colocalization with confocal microscopy. Methods 115, 42–54.

Pretzlaff, R.K., Xue, V.W., and Rowin, M.E. (2000). Sialidase treatment exposes the beta1- integrin active ligand binding site on HL60 cells and increases binding to fibronectin. Cell Adhes Commun 7, 491–500.

Preus, S., Noer, S.L., Hildebrandt, L.L., Gudnason, D., and Birkedal, V. (2015). iSMS: single-molecule FRET microscopy software. Nat Methods 12, 593–594.

Punjani, A., Rubinstein, J.L., Fleet, D.J., and Brubaker, M.A. (2017). cryoSPARC: algorithms for rapid unsupervised cryo-EM structure determination. Nat Methods 14, 290–296.

Punjani, A., Zhang, H., and Fleet, D.J. (2020). Non-uniform refinement: adaptive regularization improves single-particle cryo-EM reconstruction. Nat Methods 17, 1214–1221.

Renard, H.-F., Simunovic, M., Lemière, J., Boucrot, E., Garcia-Castillo, M.D., Arumugam, S., Chambon, V., Lamaze, C., Wunder, C., Kenworthy, A.K., et al. (2015). Endophilin-A2 functions in membrane scission in clathrin-independent endocytosis. Nature 517, 493–496.

Renard, H.F., and Boucrot, E. (2021). Unconventional endocytic mechanisms. Curr Opin Cell Biol 71, 120–129.

Renard, H.F., Tyckaert, F., Lo Guidice, C., Hirsch, T., Valades Cruz, C.A., Lemaigre, C., Shafaq-Zadah, M., Wunder, C., Wattiez, R., Johannes, L., et al. (2020). Endophilin-A3 and galectin-8 control the clathrin-independent endocytosis of CD166. Nat Commun 11, 1457.

Robinson, M.S. (2015). Forty Years of Clathrin-coated Vesicles. Traffic 16, 1210–1238.

Römer, W., Berland, L., Chambon, V., Gaus, K., Windschiegl, B., Tenza, D., Aly, M.R., Fraisier, V., Florent, J.-C., Perrais, D., et al. (2007). Shiga toxin induces tubular membrane invaginations for its uptake into cells. Nature 450, 670–675.

Salameh, B.A., Cumpstey, I., Sundin, A., Leffler, H., and Nilsson, U.J. (2010). 1H-1,2,3-triazol- 1-yl thiodigalactoside derivatives as high affinity galectin-3 inhibitors. Bioorg Med Chem 18, 5367–5378.

Sanchez-Garcia, R., Gomez-Blanco, J., Cuervo, A., Carazo, J.M., Sorzano, C.O.S., and Vargas, J. (2021). DeepEMhancer: a deep learning solution for cryo-EM volume post-processing. Commun Biol 4, 874.

Sarangi, N.K., Shafaq-Zadah, M., Berselli, G.B., Robinson, J., Dransart, D., Di Cicco, A., Levy, D., Johannes, L., and Keyes, T.E. (2022). Galectin-3 binding to apha5beta1 integrin in pore suspended biomembranes. J Phys Chem B 126, 10000–10017.

Sathe, M., Muthukrishnan, G., Rae, J., Disanza, A., Thattai, M., Scita, G., Parton, R.G., and Mayor, S. (2018). Small GTPases and BAR domain proteins regulate branched actin polymerisation for clathrin and dynamin-independent endocytosis. Nat Commun 9, 1835.

Scheres, S.H. (2012). RELION: implementation of a Bayesian approach to cryo-EM structure determination. J Struct Biol 180, 519–530.

Schindelin, J., Arganda-Carreras, I., Frise, E., Kaynig, V., Longair, M., Pietzsch, T., Preibisch, S., Rueden, C., Saalfeld, S., Schmid, B., et al. (2012). Fiji: an open-source platform for biological-image analysis. Nat Methods 9, 676–682.

Schneider, C.A., Rasband, W.S., and Eliceiri, K.W. (2012). NIH Image to ImageJ: 25 years of image analysis. Nat Methods 9, 671–675.

Schumacher, S., Dedden, D., Nunez, R.V., Matoba, K., Takagi, J., Biertümpfel, C., and Mizuno, N. (2021). Structural insights into integrin α(5)β(1) opening by fibronectin ligand. Sci Adv 7, eabe9716.

Seales, E.C., Jurado, G.A., Brunson, B.A., Wakefield, J.K., Frost, A.R., and Bellis, S.L. (2005). Hypersialylation of beta1 integrins, observed in colon adenocarcinoma, may contribute to cancer progression by up-regulating cell motility. Cancer Res 65, 4645–4652.

Seaman, M.N. (2004). Cargo-selective endosomal sorting for retrieval to the Golgi requires retromer. J Cell Biol 165, 111–122.

Shafaq-Zadah, M., Gomes-Santos, C.S., Bardin, S., Maiuri, P., Maurin, M., Iranzo, J., Gautreau, A., Lamaze, C., Caswell, P., Goud, B., et al. (2016). Persistent cell migration and adhesion rely on retrograde transport of beta1 integrin. Nat Cell Biol 18, 54–64.

Shi, F., and Sottile, J. (2011). MT1-MMP regulates the turnover and endocytosis of extracellular matrix fibronectin. J Cell Sci 124, 4039–4050.

Shi, G., Azoulay, M., Dingli, F., Lamaze, C., Loew, D., Florent, J.C., and Johannes, L. (2012). SNAP-tag based proteomics approach for studying retrograde transport. Traffic 13, 914–925.

Simunovic, M., Manneville, J.B., Renard, H.F., Evergren, E., Raghunathan, K., Bhatia, D., Kenworthy, A.K., Voth, G.A., Prost, J., McMahon, H., et al. (2017). Friction mediates scission of tubular membrane scaffolded by BAR proteins. Cell 170, 172–184.

Sörme, P., Arnoux, P., Kahl-Knutsson, B., Leffler, H., Rini, J.M., and Nilsson, U.J. (2005). Structural and thermodynamic studies on cation-Pi interactions in lectin-ligand complexes: high-affinity galectin-3 inhibitors through fine-tuning of an arginine-arene interaction. J Am Chem Soc 127, 1737–1743.

Sottile, J., and Chandler, J. (2005). Fibronectin matrix turnover occurs through a caveolin-1- dependent process. Mol Biol Cell 16, 757–768.

Stegmayr, J., Zetterberg, F., Carlsson, M.C., Huang, X., Sharma, G., Kahl-Knutson, B., Schambye, H., Nilsson, U.J., Oredsson, S., and Leffler, H. (2019). Extracellular and intracellular small-molecule galectin-3 inhibitors. Sci Rep 9, 2186.

Suzuki, T., Suzuki, M., Ogino, S., Umemoto, R., Nishida, N., and Shimada, I. (2015). Mechanical force effect on the two-state equilibrium of the hyaluronan-binding domain of CD44 in cell rolling. Proc Natl Acad Sci USA 112, 6991–6996.

von Mach, T., Carlsson, M.C., Straube, T., Nilsson, U., Leffler, H., and Jacob, R. (2014). Ligand binding and complex formation of galectin-3 is modulated by pH variations. Biochem J 457, 107–115.

Wagner, T., Merino, F., Stabrin, M., Moriya, T., Antoni, C., Apelbaum, A., Hagel, P., Sitsel, O., Raisch, T., Prumbaum, D., et al. (2019). SPHIRE-crYOLO is a fast and accurate fully automated particle picker for cryo-EM. Commun Biol 2, 218.

Yu, S., Fan, J., Liu, L., Zhang, L., Wang, S., and Zhang, J. (2013). Caveolin-1 up-regulates integrin α2,6-sialylation to promote integrin α5β1-dependent hepatocarcinoma cell adhesion. FEBS Lett 587, 782–787.

Zhao, H., Przybylska, M., Wu, I.H., Zhang, J., Siegel, C., Komarnitsky, S., Yew, N.S., and Cheng, S.H. (2007). Inhibiting glycosphingolipid synthesis improves glycemic control and insulin sensitivity in animal models of type 2 diabetes. Diabetes 56, 1210–1218.

Zhao, Z., Xu, X., Cheng, H., Miller, M.C., He, Z., Gu, H., Zhang, Z., Raz, A., Mayo, K.H., Tai, G., et al. (2021). Galectin-3 N-terminal tail prolines modulate cell activity and glycan-mediated oligomerization/phase separation. Proc Natl Acad Sci USA 118, e2021074118.

Zheng, S.Q., Palovcak, E., Armache, J.P., Verba, K.A., Cheng, Y., and Agard, D.A. (2017). MotionCor2: anisotropic correction of beam-induced motion for improved cryo-electron microscopy. Nat Methods 14, 331–332.

Zhuo, Y., and Bellis, S.L. (2011). Emerging role of alpha2,6-sialic acid as a negative regulator of galectin binding and function. J Biol Chem 286, 5935–5941.

Zhuo, Y., Chammas, R., and Bellis, S.L. (2008). Sialylation of beta1 integrins blocks cell adhesion to galectin-3 and protects cells against galectin-3-induced apoptosis. J Biol Chem 283, 22177–22185.

