## Supplemental Figures, Table and Movies captions for "Spatial N-glycan rearrangement on α_5_β_1_ integrin nucleates galectin-3 oligomers to determine endocytic fate"

**This supplemental file includes:**

Figures S1 to S8
Tables S1
Movies captions S1 to S6

### SUPPLEMENTAL FIGURES

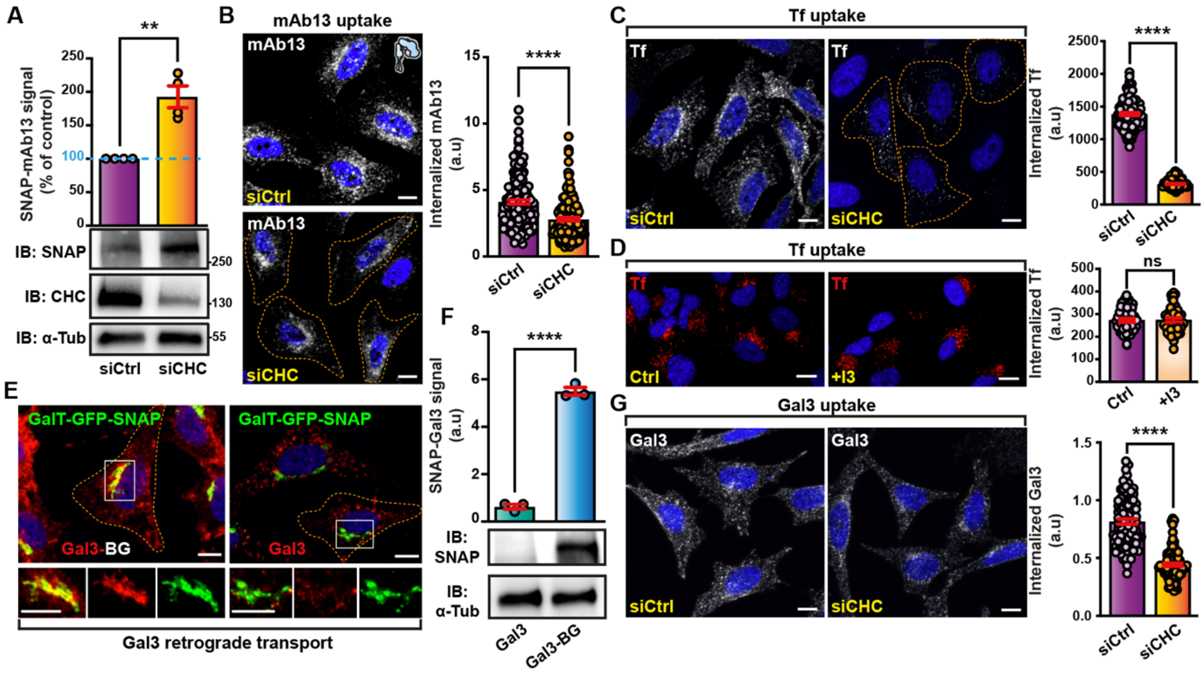

**Figure S1. Clathrin-independent retrograde trafficking of the inactive bent-closed $\alpha_5\beta_1$ integrin depends on Gal3. (A)** Use of GFP-trap to pull down stably expressed Golgi-localized GalT-GFP-SNAP from HeLa cells that had continuously been incubated for 3 h at 37 °C with mAb13-BG, either in control condition (siCtrl) or after inhibition of clathrin expression (siCHC). mAb13-BG that reached the Golgi was quantified by immunoblotting (IB: SNAP) for the corresponding SNAP-mAb13 conjugates. Clathrin inhibition was assessed by immunoblotting (IB: CHC). α-tubulin was used for normalization (IB: α-Tub). 3 independent experiments, means ± SEM, unpaired t-test; $^{**}P < 0.002$. (**B**) Quantification by confocal microscopy of mAb13 uptake after continuous incubation for 10 min at 37 °C in HeLa cells (GalT-GFP-SNAP), either in control condition (siCtrl), or after inhibition of clathrin expression (siCHC). 3 independent experiments, means ± SEM, unpaired t-test; $^{****}P < 0.0001$. Scale bars = 10 µm. Nuclei in blue. (**C**) Quantification by confocal microscopy of Transferrin (Tf) uptake after continuous incubation for 10 min at 37 °C in HeLa cells (GalT-GFP-SNAP), either in control condition (siCtrl), or after inhibition of clathrin expression (siCHC). 3 independent experiments, means ± SEM, unpaired t-test; $^{****}P < 0.0001$. Scale bars = 10 µm. Nuclei in blue. (**D**) I3 effect on Tf uptake. Tf was continuously incubated for 10 min at 37 °C with RPE-1 cells that were either untreated or pre-treated with the cell-impermeable Gal3 inhibitor I3. Signals coming from internalized Tf were quantified. 3 independent experiments, means ± SEM, unpaired t-test; ns = $P > 0.05$. Scale bars = 10 µm. Nuclei in blue. (**E**) Gal3 trafficking to the Golgi. Continuous incubation of GalT-GFP-SNAP-expressing HeLa cells for 1 h at 37 °C with Cy3-labelled Gal3 (200 nM) that was also BG-coupled, or not. Note that Gal3-BG colocalized in perinuclear Golgi with GalT-GFP-SNAP. Scale bars = 10 µm. Nuclei in blue. (**F**) Use of GFP-trap to pull down stably expressed Golgi-localized GalT-GFP-SNAP from HeLa cells that had continuously been incubated for 3 h at 37 °C with Gal3 or Gal3-BG. SNAP-Gal3 was quantified by immunoblotting (IB: SNAP). α-tubulin was used for normalization (IB: α-Tub). 3 independent experiments, means ± SEM, unpaired t-test; $^{****}P < 0.0001$. (**G**) Quantification by confocal microscopy of Gal3 uptake (200 nM) after continuous incubation for 10 min at 37 °C in HeLa cells (GalT-GFP-SNAP), either in control condition (siCtrl), or after inhibition of clathrin expression (siCHC). 3 independent experiments, means ± SEM, unpaired t-test; ****$P < 0.0001$. Scale bars = 10 µm. Nuclei in blue.

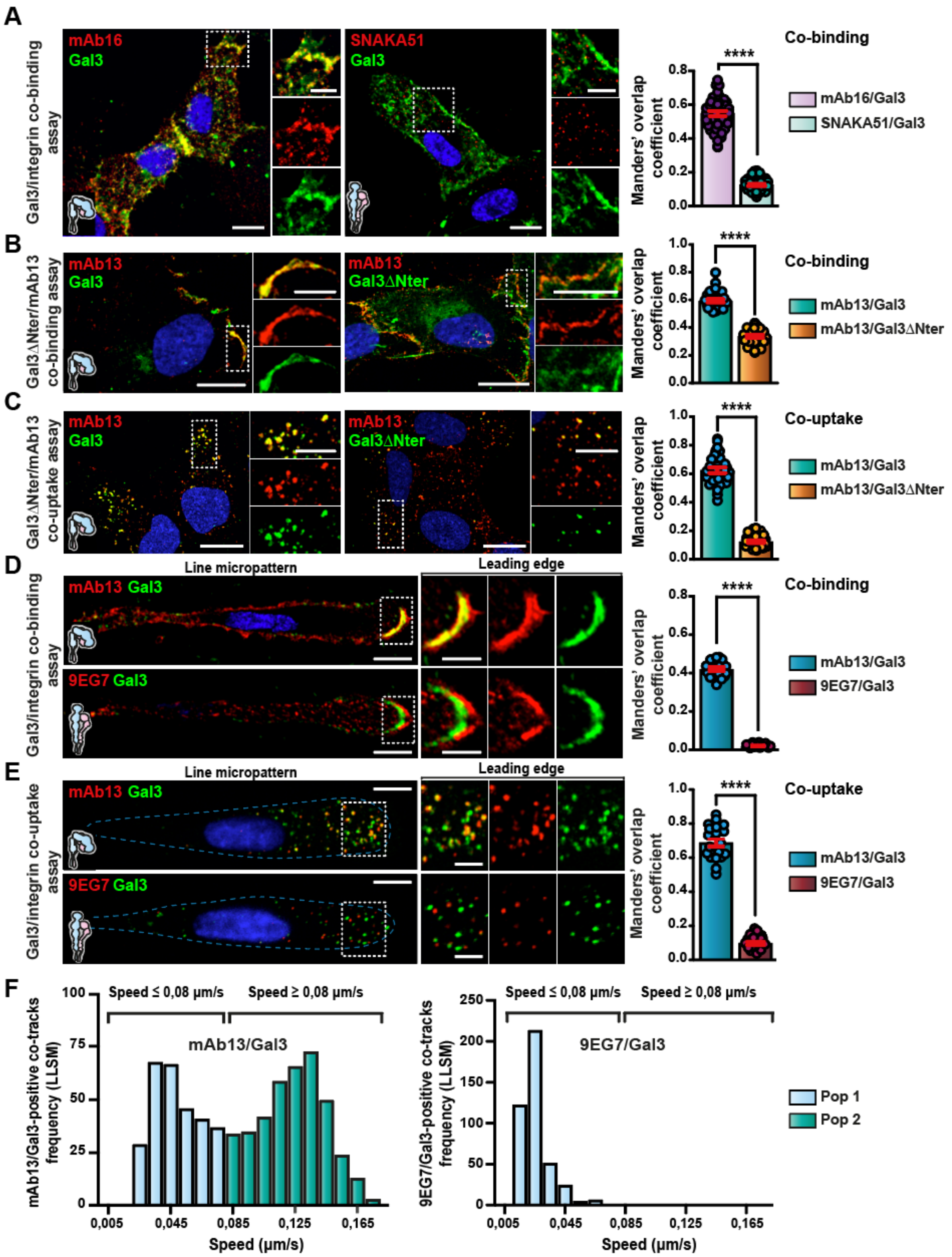

**Figure S2. Gal3 oligomerization capacity is required for its preferential binding to the inactive bent-closed $\alpha_5\beta_1$ integrin.** (**A**) Co-binding with $\alpha_5$ integrin. RPE-1 cells were sequentially incubated at 4 °C with 200 nM of Gal3 and the conformation-specific $\alpha_5$ integrin antibodies mAb16 (inactive) or SNAKA51 (active), and directly fixed. The overlap of fluorescence signals was quantified. 3 independent experiments, means ± SEM, unpaired t-test; ****$P < 0.0001$. Scale bars = 10 µm, and 5 µm in zoomed insets. Nuclei in blue. (**B**) Co-binding with $\beta_1$ integrin. RPE-1 cells were sequentially incubated at 4 °C with 200 nM of Gal3 or

Gal3ΔNter and mAb13 antibodies and directly fixed. The overlap of fluorescence signals from 3 independent experiments was quantified. Means ± SEM, unpaired t-test; ****P < 0.0001. Scale bars = 10 µm, and 5 µm in zoomed insets. Nuclei in blue. (**C**) Co-uptake with $\beta_1$ integrin. RPE-1 cells were sequentially incubated at 4 °C with 200 nM of Gal3 or Gal3ΔNter and mAb13 antibodies and then shifted for 10 min to 37 °C. The overlap of fluorescence signals from 3 independent experiments was quantified. Means ± SEM, unpaired t-test; ****P < 0.0001. Scale bars = 10 µm, and 5 µm in zoomed insets. Nuclei in blue. (**D**) Same as in (B), using Gal3 and either mAb13 or 9EG7, in RPE-1 cells that were seeded onto line micropatterns. Of note, Gal3 was massively distributed towards the leading edge and specifically overlapped with mAb13. The overlap of fluorescence signals from 3 independent experiments was quantified. Means ± SEM, unpaired t-test; ****P < 0.0001. Scale bars = 10 µm, and 5 µm in zoomed insets. Nuclei in blue. (**E**) Same as in (C), using Gal3 and either mAb13 or 9EG7, in RPE-1 cells that were seeded onto line micropatterns. The overlap of fluorescence signals from 3 independent experiments was quantified. Means ± SEM, unpaired t-test; ****P < 0.0001. Scale bars = 10 µm, and 5 µm in zoomed insets. Nuclei in blue. (**F**) Lattice light sheet microscopy. Co-tracking of Cy3-labeled Gal3 with ATTO488-labeled mAb13 (left panel) or 9EG7 (right panel) antibodies. Frequency distributions of Gal3-positive mAb13 or 9EG7 co-tracks in function of their velocity. Of note, the dynamic population (Pop 2) was totally absent for 9EG7/Gal3 co-tracks, indicating that 9EG7/Gal3-containing structures were largely immobile.

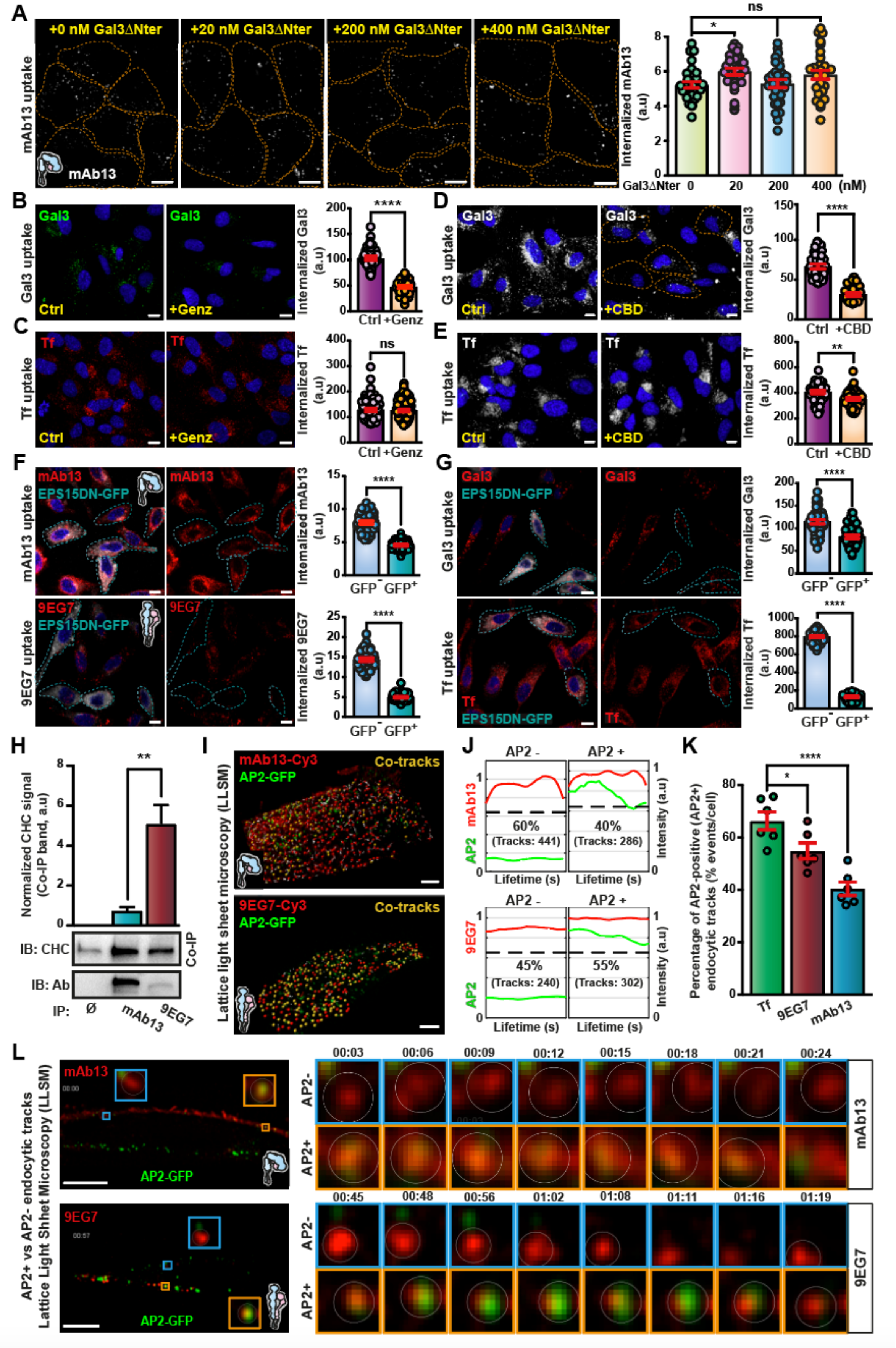

**Figure S3. Internalization of inactive bent-closed $\alpha_5\beta_1$ integrin relies on the GL-Lect driven endocytosis.** (A) Gal3ΔNter effect on mAb13 uptake. mAb13 antibody was bound at

4 °C to RPE-1 cells, which were then incubated for 10 min at 37 °C in the absence or the presence of the indicated concentrations of exogenous Gal3ΔNter. Signals coming from internalized mAb13 were quantified. Dashed lines represent the contours of cells. 3 independent experiments, means ± SEM, one-way ANOVA; ns = P > 0.05; *P < 0.02. Scale bar = 10 µm. (**B,C**) Glycosphingolipid (GSL) depletion effect on endocytosis. Gal3 (200 nM) (B), or Transferrin (Tf) (C) were continuously incubated for 5 min or 10 min, respectively, at 37 °C with RPE-1 cells that were pre-treated or not with the GSL synthesis inhibitor Genz. Signals coming from internalized Gal3 or Tf were quantified. 3 independent experiments, means ± SEM, unpaired t-test; ns = P > 0.05, ****P < 0.0001. Scale bars = 10 µm. Nuclei in blue. (**D**) Effect of the dynein inhibitor ciliobrevin D (CBD) on Gal3 endocytosis. 200 nM of Gal3 was continuously incubated for 10 min at 37 °C with RPE-1 cells that were treated or not with CBD. 3 independent experiments, means ± SEM, unpaired t-test; ****P < 0.0001. Scale bars = 10 µm. Nuclei in blue. (**E**) Same as in (D), for Transferrin (Tf) uptake. 3 independent experiments, means ± SEM, unpaired t-test; **P < 0.002. Scale bars = 10 µm. Nuclei in blue. (**F**) Effect of Epsin 15 dominant-negative (EPS15DN-GFP) on $\alpha_5\beta_1$ integrin endocytosis. Quantification by confocal microscopy of mAb13 and 9EG7 uptake after continuous incubation for 10 min at 37 °C with RPE-1 cells. Dashed lines indicate EPS15DN-GFP expressing cells. 3 independent experiments, means ± SEM, unpaired t-test; ****P<0.0001. Scale bars = 10 µm. Nuclei in blue. (**G**) Effect of EPS15DN-GFP on Gal3 and Transferrin (Tf) endocytosis. Gal3 (200 nM), or Tf were continuously incubated for 10 min at 37 °C with RPE-1 cells transiently transfected with EPS15DN-GFP. Dashed lines indicate cells that were GFP-positive ($GFP^+$) and therefore expressed the dominant negative mutant. The GFP-negative ($GFP^-$) cells were used as internal controls. Fluorescence signals were analyzed and quantified by confocal microscopy. 3 independent experiments, means ± SEM, unpaired t-test; ****P < 0.0001. Scale bars = 10 µm. Nuclei in blue. (**H**) $\alpha_5\beta_1$ integrin interaction with clathrin heavy chain (CHC) in RPE-1 cells. Cell surface immunoprecipitation (IP) with mAb13, 9EG7, and beads (Ø) as controls. Immunoblot (IB) for indicated antigens. 3 independent experiments, means ± SEM, unpaired t-test; **P < 0.002. (**I**) Monitoring mAb13 and 9EG7 endocytosis using lattice light sheet microscopy on AP2-GFP genome-edited RPE-1 cells. 3D projections of acquired 3D stacks of cargo (red) and AP2 (green). Events (yellow) for which the indicated markers were co-tracked during endocytic uptake. Scale bars = 8 µm. (**J**) Median normalized endocytic intensity tracks of mAb13 and 9EG7 uptake from experiments as in (I). Intensities below dashed black lines were considered background. (**K**) Percentages of AP2 positive uptake events from (H) for mAb13 (727 total tracks), 9EG7 (542 total tracks), and Tf (1333 total tracks). 6 cells per condition, means ± SEM, one-way ANOVA; *P < 0.05, ****P < 0.0001. (**L**). Experiment as in (I). Left, 2D side views of cell slices perpendicular to the detection objective. Examples of mAb13 or 9EG7-positive structures that overlapped (orange squares, AP2+) or not (blue squares, AP2-) with AP2-GFP in RPE-1 cells. Scale bars = 10 µm. Right, time-resolved evolution (min:sec) of mAb13 and 9EG7 signals in relation to AP2-GFP. Examples of AP2 positive (AP2+) and AP2 negative (AP2-) tracks are shown. Images were extracted from Movies S3 and S4.

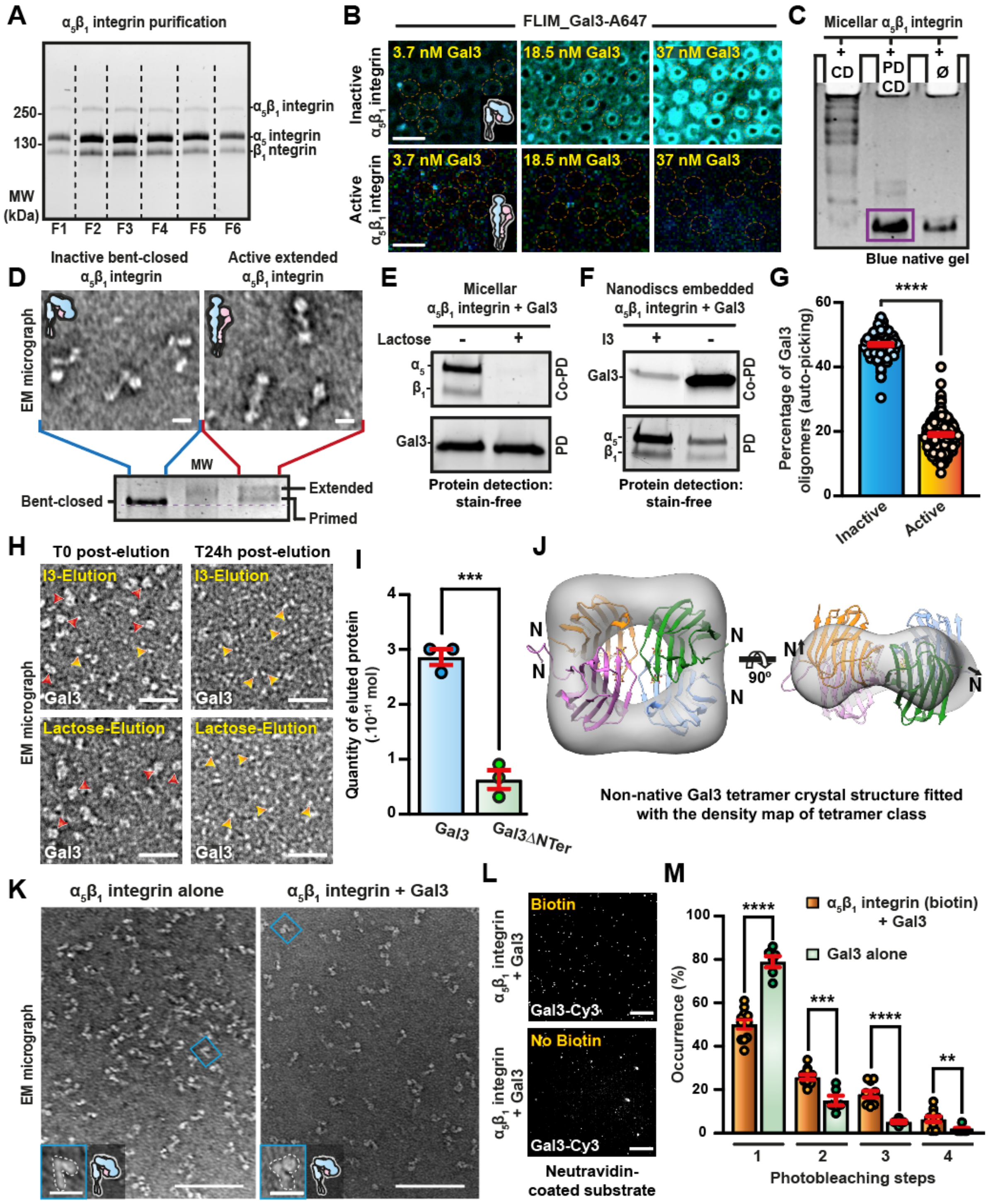

**Figure S4. Specifically, the inactive bent-closed $\alpha_5\beta_1$ integrin nucleates the formation of Gal3 oligomers.** (**A**) $\alpha_5\beta_1$ integrin purification. $\alpha_5\beta_1$ integrin was solubilized from rat liver and purified in a two-step process using wheat germ agglutinin and fibronectin affinity columns. Fractions F1 to F6 from the fibronectin column were analyzed by SDS-PAGE and total protein staining to assess purity. (**B**) FLIM measurements on MSLB confirming the preferential interaction of Gal3 with inactive $\alpha_5\beta_1$ integrin in this minimal membrane environment. Dashed circles indicate microcavity pores. Scale bars = 5 µm. (**C**) Reconstitution of $\alpha_5\beta_1$ integrin into peptidiscs. The reconstitution from Triton X-100 micelles into peptidiscs was analyzed by blue native gel electrophoresis. Upon detergent removal using cyclodextrin (+CD) and in the absence of Apo-1 derived peptides, several bands were visible, indicative of protein aggregation. However, in the presence of Apo1-derived peptides and cyclodextrin (PD+CD), a major band was found (purple rectangle) that migrated at the same level as micellar integrin (Ø). These data document an efficient mono-incorporation of $\alpha_5\beta_1$ integrin into peptidiscs. (**D**) Electron micrographs of $\alpha_5\beta_1$ integrin in peptidiscs. Left: inactive Bent-closed inactive conformation. Right: The protein was incubated with 5 mM $MnCl_2$ and 100 µM cRGD. Note the switch from the inactive bent-closed to the active extended ligand-bound conformational state. Scale bars = 10 nm. These samples were also analyzed by blue native PAGE, which allowed to document corresponding changes in electrophoretic mobility. (**E**) Glycan-dependent binding of Gal3 to micellar $\alpha_5\beta_1$ integrin. 200 nM of Gal3-His was pre-incubated or not with lactose, and then co-incubated with micellar $\alpha_5\beta_1$ integrin (Co-PD). After pulldown (PD) on cobalt beads, proteins were separated on SDS-PAGE gels and visualized. Note that lactose efficiently competed with Gal3 binding to micellar $\alpha_5\beta_1$ integrin, as expected for a glycan-dependent interaction between both proteins. (**F**) Glycan-dependent binding of Gal3 to nanodisc-embedded $\alpha_5\beta_1$ integrin (PD). Untagged 200 nM of Gal3 (Co-PD) was pre-incubated or not with the I3 compound, and then co-incubated with $\alpha_5\beta_1$ integrin in nanodiscs immobilized on cobalt beads. Samples were analyzed as in (E). The loss of Gal3 from $\alpha_5\beta_1$ integrin in the presence of I3 again was consistent with a glycan-dependent interaction between both proteins. (**G**) Quantification of Gal3 oligomers by particle autopicking in crYOLO (Wagner et al., 2019). $\alpha_5\beta_1$ integrin was embedded in His-tagged nanodiscs, immobilized on cobalt beads, activated with $MnCl_2$/cRGD or not, and incubated with Gal3. I3 was used to elute Gal3, which was analyzed in all conditions by negative staining EM. Oligomers were quantified by automatic picking of particles. Note that erroneous scoring of irrelevant contaminating objects such as nanodiscs could not be excluded. Inactive $\alpha_5\beta_1$ integrin/Gal3: 100 EM-fields; active $\alpha_5\beta_1$integrin/Gal3: 145 EM-fields. Means ± SEM, unpaired t-test; ****$P < 0.0001$. (**H**) EM micrographs of Gal3 elution as in (G) in the inactive $\alpha_5\beta_1$ integrin/Gal3 condition that were analyzed right away (T0), or after 24 h storage in solution at 4 °C (T24 h). Note that Gal3 oligomers (red arrowheads) had disassembled into monomers (orange arrowheads) after 24 h. Scale bars = 20 nm. (**I**) Quantification of Gal3 and Gal3ΔNter binding to inactive $\alpha_5\beta_1$ integrin. After binding (4 µM) and elution as in (G), Gal3 and Gal3ΔNter were loaded on a denaturing SDS-PAGE, and protein signals (stain-free detection) were quantified. 3 independent experiments, means ± SEM, unpaired t-test; ***$P < 0.0002$. (**J**) Fit of a Gal3 tetramer crystallized under non-native conditions (PDB 6FOF) into the density map of Gal3 tetramers eluted from RPE1 cells as in Figure 3D. (**K**) EM micrographs of peptidisc-embedded $\alpha_5\beta_1$ integrin, either alone (left panel) or after co-incubation with 4 µM of Gal3 (right panel). Scale bars = 100 nm. Insets show individual integrin particles; scale bars = 20 nm. (**L**) Peptidisc immobilization for photobleaching experiments. $\alpha_5\beta_1$ integrin that was embedded into biotin-tagged or untagged peptidiscs, complexed with Cy3-Gal3 as described in K, and incubated with neutravidin-coated substrate. Note that in the absence of biotin, expectedly fewer Gal3 spots were visible on the substrate. Scale bars = 8 µm. (**M**) Photobleaching experiments performed with biotin-tagged Gal3-Cy3/$\alpha_5\beta_1$ integrin peptidiscs, or Gal3-Cy3 alone. Bar plots show the number of steps detected via fluorescence imaging. For Gal3 alone, predominantly single-step photobleaching behavior was observed, while higher number of photo-bleaching steps were measured for $\alpha_5\beta_1$integrin-Gal3 complexes, which we interpret as Gal3 oligomers. 11 fields for the $\alpha_5\beta_1$integrin-Gal3 condition; 6 fields for the Gal3 alone condition.

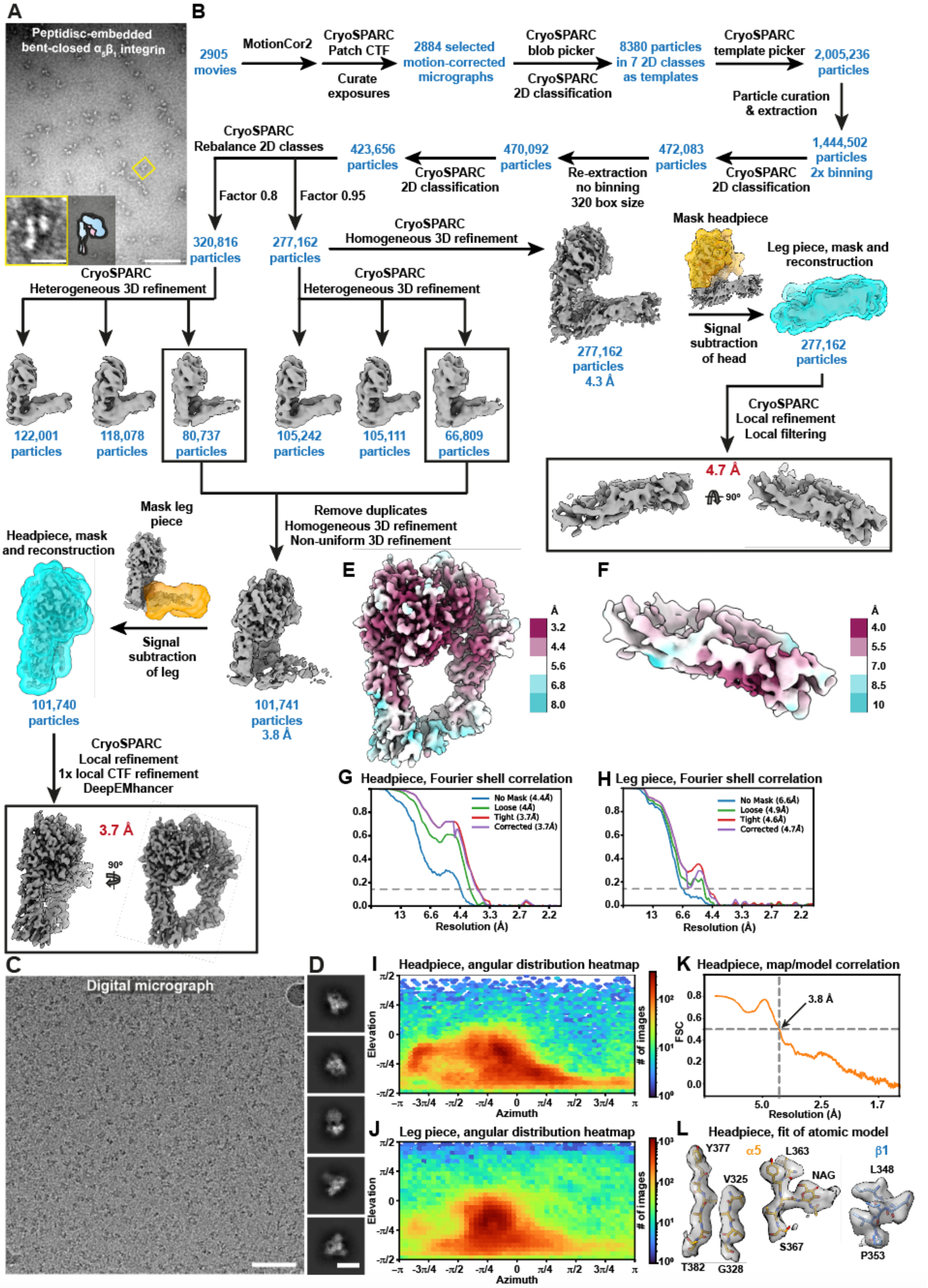

**Figure S5. Cryo-EM processing of peptidisc-embedded $\alpha_5\beta_1$ integrin.** (**A**) Negative stain EM micrograph showing efficient mono-insertion of $\alpha_5\beta_1$ integrin into peptidiscs. Scale bar = 100 nm and 20 nm for zoom-in. (**B**) Schematics of the cryo-EM data processing pipeline, including signal subtraction and local refinements of the headpiece and leg piece. (**C**) Typical digital micrograph at 2.0 µm defocus, recorded with a Falcon-3 camera at 300 kV. Scale bar = 50 nm. (**D**) Representative 2D class averages of $\alpha_5\beta_1$ integrin, showing different projection views. Scale bar = 10 nm. (**E,F**) Density maps of the headpiece (E) and leg piece (F) of $\alpha_5\beta_1$ integrin, colored by local resolution as calculated in CryoSPARC. (**G,H**) Fourier shell correlation (FSC) curves of the final local refinements of cryo-EM densities of headpiece (G) and leg piece (H). (**I,J**) Angular distribution heatmaps of the particles used in the final refinements of headpiece (H) and leg piece (I). (**K**) Map-to-model correlation of the headpiece of $\alpha_5\beta_1$ integrin ($\alpha_5$, residues 94 - 691, $\beta_1$, 25-504). (**L**) Fit of atomic model in selected parts of the headpiece (left: $\alpha_5$, β-propeller, middle: $\alpha_5$, glycan at N365, right: $\beta_1$, α-helix).

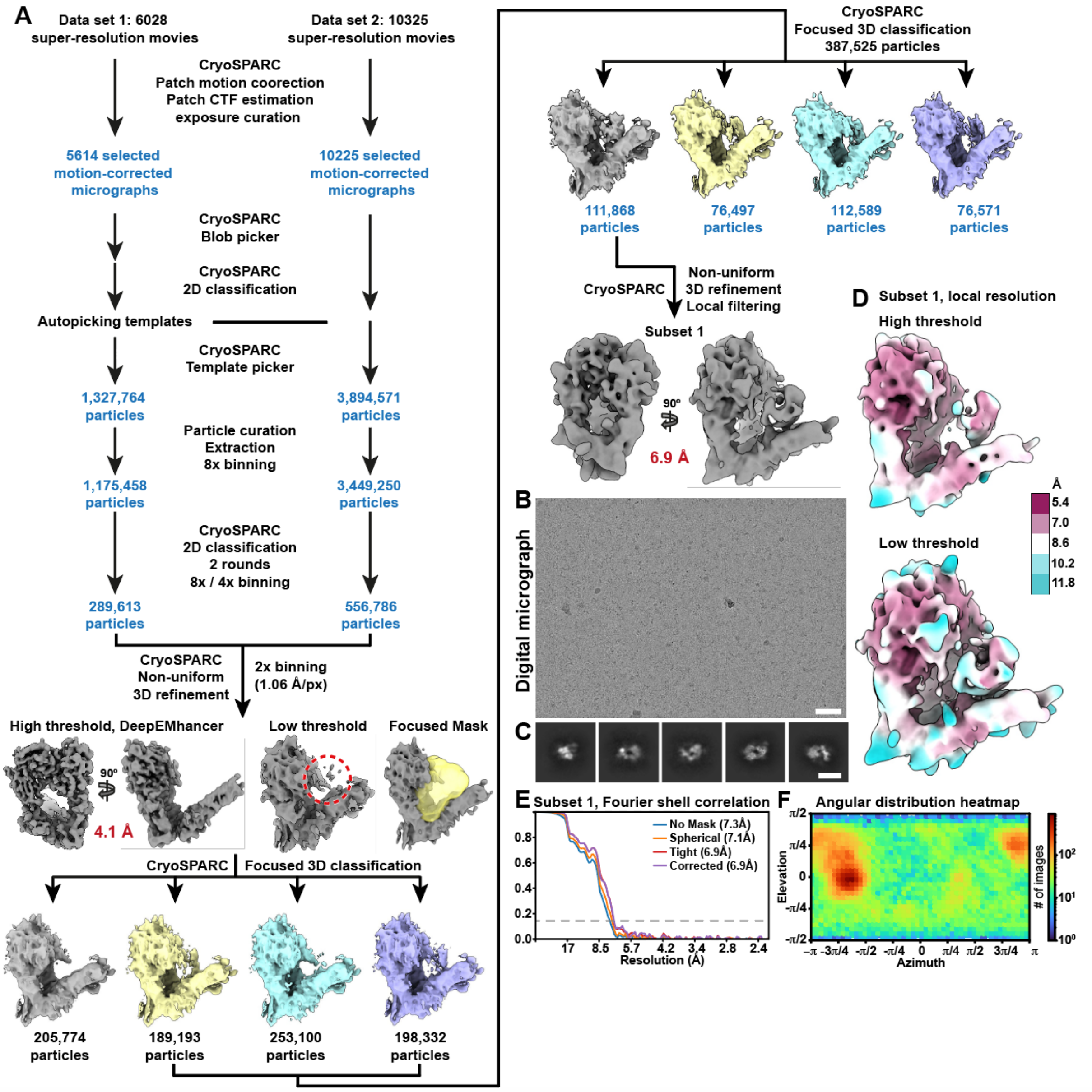

**Figure S6. Cryo-EM processing of peptidisc-embedded $\alpha_5\beta_1$ integrin in complex with Gal3**. (**A**) Schematic of the cryo-EM data processing pipeline of the $\alpha_5\beta_1$ integrin/Gal3 complex. With 111,868 particles, subset 1 with the most clearly defined density between headpiece and leg piece accounted for 13% of the total data set, indicating that only a fraction of $\alpha_5\beta_1$ integrin in this preparation was competent for Gal3 binding and oligomerization. (**B**) Typical digital micrograph at 2.0 µm defocus, recorded with a Gatan K3 camera at 300 kV. Scale bar = 50 nm. (**C**) Representative 2D class averages of the $\alpha_5\beta_1$ integrin/Gal3 complex, showing different projection views. Scale bar = 10 nm. (**D**) Cryo-EM density map of subset 1 of the $\alpha_5\beta_1$ integrin/Gal3 complex, colored by local resolution as obtained in CryoSPARC. (**E**) Fourier shell correlation (FSC) curves of the final non-uniform refinement of subset 1 of the $\alpha_5\beta_1$ integrin/Gal3 complex. (**F**) Angular distribution heatmap of the particles used in the final non-uniform refinement of subset 1 of the $\alpha_5\beta_1$ integrin/Gal3 complex.

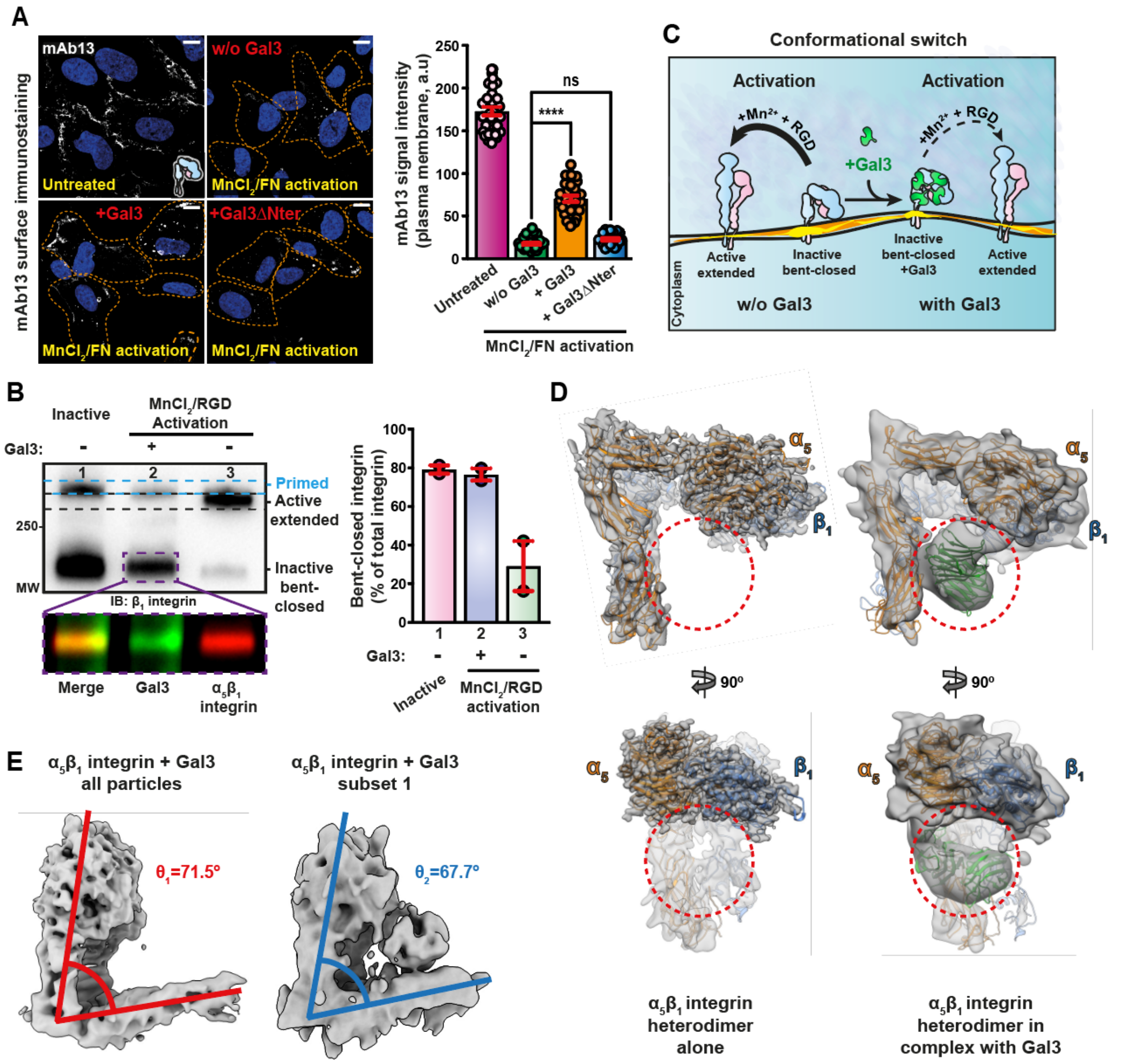

**Figure S7. Gal3 clamps the inactive bent-closed conformational state of $\alpha_5\beta_1$ integrin.** (**A**) Effect of Gal3 on $\alpha_5\beta_1$ integrin activation in cells. RPE-1 cells were incubated or not at 4 °C with either Gal3 or Gal3ΔNter (200 nM). Cells were then incubated or not for 1 h at 4 °C with 1 mM $MnCl_2$ and 5 µg/mL soluble fibronectin (FN) to switch integrins to the active extended ligand-bound conformation. After fixation, the cells were immunolabeled with mAb13 antibody against the inactive bent-closed conformation. Scale bars = 10 µm. Nuclei in blue. Quantification of 3 independent experiments showed that Gal3 binding to $\alpha_5\beta_1$ integrin reduced its propensity to be activated, an effect that was dependent on Gal3's oligomerization capacity as it was not observed with the Gal3ΔNter mutant. Means ± SEM, one-way ANOVA; ns = P > 0.05; ****P < 0.0001. (**B**) Effect of Gal3 on $\alpha_5\beta_1$ integrin activation. Micellar $\alpha_5\beta_1$ integrin was pre-incubated or not with 2 µM of Gal3-Alexa488 and then incubated with 5 mM $MnCl_2$ and 100 µM cRGD. Samples were loaded on semi-native gels and bands were detected by fluorescence and total protein staining. Inactive $\alpha_5\beta_1$ integrin showed a dominant lower band, likely the inactive bent-closed conformer (lane 1), and a weaker upper band, likely representing the primed conformational state (delimited by blue dashed line). Upon incubation with $MnCl_2$ and cRGD, a clear band switch was observed in favor of a new upper band (delimited by black dashed line), likely corresponding to the active extended ligand-bound conformation (lane 3). Upon pre-incubation with Gal3, this band switch was strongly reduced (lane 2). Note that as expected, Gal3 and $\alpha_5\beta_1$ integrin overlapped in the lower band of lane 2. In all conditions, the lower bands (i.e., the inactive bent-closed conformer) were then quantified from 2 independent experiments as the percentage of total $\alpha_5\beta_1$ integrin. Means ± SEM are shown. (**C**) Schematic summary of results in (A) and (B). (**D**) Cryo-EM density map of peptidisc-embedded $\alpha_5\beta_1$ integrin alone (left) or in complex with Gal3, with modeled $\alpha_5\beta_1$ integrin (orange for $\alpha_5$ and blue for $\beta_1$ subunits). The red dashed circle clearly indicates the presence of additional cryo-EM densities when incubated with Gal3, with two modeled Gal3 CRDs as in Fig. 6E. (**E**) Comparison of the angles between the headpiece and leg piece of $\alpha_5\beta_1$ integrin-Gal3 complex. Unsharpened maps are shown. Left: density map of the full data set, including many $\alpha_5\beta_1$ integrin without bound Gal3 and non-cohesive $\alpha_5\beta_1$ integrin-Gal3 complexes. Right: Subset 1 with visible additional densities that fitted with Gal3 binding to $\alpha_5\beta_1$ integrin. A larger angle was observed in the full data set ($\theta_1 = 71.5°$ all particle) compared to subset 1 ($\theta_2 = 67.7°$, $\Delta\theta = 3.8°$), indicating that Gal3 binding led to a compaction of $\alpha_5\beta_1$ integrin, likely by clamping the protein in the inactive bent-closed conformational state that prevents activation, as seen in the experiments of (A) and (B).

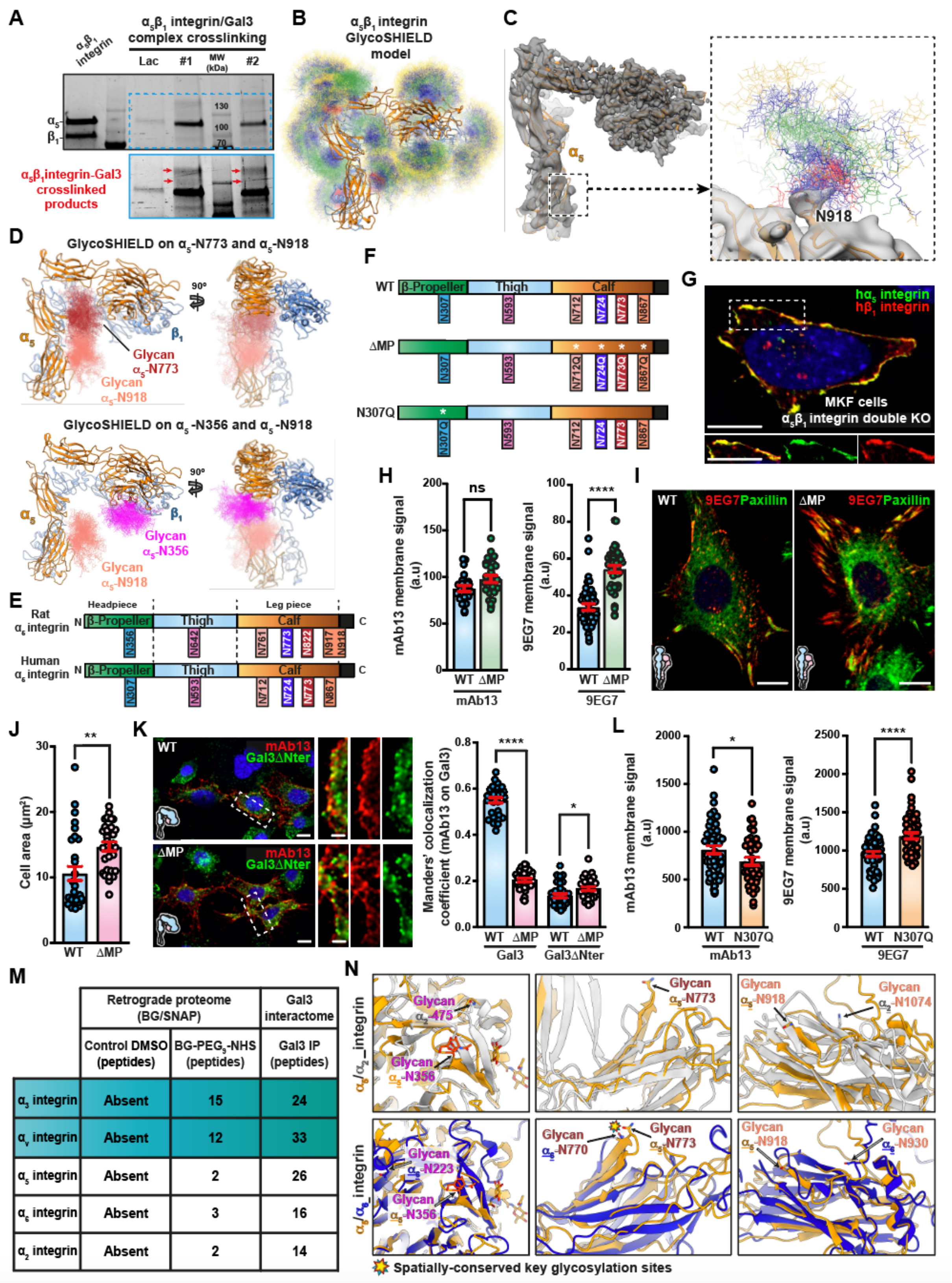

| | Retrograde proteome (BG/SNAP) | | Gal3 interactome |
|---|---|---|---|
| | Control DMSO (peptides) | BG-PEG$_9$-NHS (peptides) | Gal3 IP (peptides) |
| $\alpha_3$ integrin | Absent | 15 | 24 |
| $\alpha_v$ integrin | Absent | 12 | 33 |
| $\alpha_5$ integrin | Absent | 2 | 26 |
| $\alpha_6$ integrin | Absent | 3 | 16 |
| $\alpha_2$ integrin | Absent | 2 | 14 |

**Figure S8. Specifically inactive bent-closed $\alpha_5\beta_1$ integrin displays defined spatial glycan positioning for Gal3-driven endocytic uptake.** (A) Cross-linking data analysis on gels. 2 cross-linking reactions (lanes #1 and #2) between $\alpha_5\beta_1$ integrin and Gal3 were analyzed by SDS-PAGE. The zoomed box with stronger exposure revealed bands with lower electrophoretic mobility (red arrows) than $\alpha_5$ and $\beta_1$ integrin chains, likely representing cross-linking products. These bands were lost in the presence of lactose (Lac), which indicated that their appearance relied on glycan-Gal3 interaction. (B) 3D-structure of $\alpha_5\beta_1$ integrin with GlycoSHIELD models of all identified N-glycans in rat $\alpha_5$ integrin. Blue, N-acetylglucosamine; green, mannose; red, fucose; yellow, galactose. 100 possible conformations are shown for each glycan position. (C) $\alpha_5\beta_1$ integrin model in cryo-EM density map. The box highlights the proximal leg piece of $\alpha_5$ integrin with a density that can be ascribed to a glycan at position N918. Glycan conformations on N918 (42 projections by GlycoSHIELD) are shown. (D) 3D-structure model of $\alpha_5\beta_1$ integrin. Glycan conformations as obtained by GlycoSHIELD are shown for glycans at $\alpha_5$-N773 and $\alpha_5$-N918 (top panels) and $\alpha_5$-N356 and $\alpha_5$-N918 (lower panels), respectively. (E) Linear alignment of key N-glycosylation sites in rat and human $\alpha_5$ integrins, as identified by cross-linking proteomics and cryo-EM. (F) Linear representation of human $\alpha_5$ integrin, and N-glycosylation sites whose role in Gal3 binding were identified by cross-linking proteomics and cryo-EM. N to Q mutations in ΔMP and N307Q are indicated by white stars. (G) Transient expression of human $\alpha_5$ integrin-GFP (green) and human $\beta_1$ integrin-Halo tag (red) in MKF-dKO cells. The co-localization (yellow) between both chains indicates that these heterodimerized and successfully reached the plasma membrane. Scale bars = 10 µm. Nuclei in blue. (H) Expression of $\alpha_5\beta_1$ integrin at the cell surface. mAb13 or 9EG7 antibody binding experiments were performed at 4 °C in wildtype and ΔMP $\alpha_5$ integrin/wildtype $\beta_1$ integrin-expressing MKF-dKO cells. Plasma membrane signals of mAb13 and 9EG7 were quantified from 3 independent experiments. Means ± SEM, unpaired t-test; ns = P > 0.5, ****P < 0.0001. (I) 9EG7 antibody binding experiment at 4 °C as in (H), with immunolabeling against the focal adhesion protein paxillin. Scale bars = 10 µm. Nuclei in blue. (J) Cell areas from 2 independent experiments as in (I) were measured to assess cell spreading. Means ± SEM, unpaired t-test; **P < 0.002. (K) Gal3 or Gal3ΔNter (200 nM) and mAb13 or 9EG7 co-binding experiments within the wildtype or ΔMP $\alpha_5$ integrin/wildtype $\beta_1$ integrin genetic background of MKF-dKO cells. Colocalization of mAb13 and 9EG7 with Gal3 or Gal3ΔNter was quantified. 3 independent experiments, means ± SEM, unpaired t-test; *P < 0.02, ****P < 0.0001. Scale bars = 5 µm. Nuclei in blue. (L) Expression of $\alpha_5\beta_1$ integrin at the cell surface. mAb13 or 9EG7 binding experiments were performed at 4°C within the wildtype or N307Q $\alpha_5$ integrin/wildtype $\beta_1$ integrin genetic background of MKF-dKO cells. Plasma membrane signals of mAb13 and 9EG7 were quantified. 3 independent experiments, means ± SEM, unpaired t-test; *P < 0.02, ****P < 0.0001. (M) Table summarizing α integrins that were commonly found in both the retrograde proteome (BG-PEG$_9$-NHS column) and the Gal3 interactome (Gal3 IP column) (Lakshminarayan et al., 2014). DMSO treated cells served as control. Note that α3 and αv integrins were highly abundant in both mass spectrometry analysis. (N) Overlay of integrin 3D structures at key N-glycosylation sites identified for rat $\alpha_5$ integrin (orange). 3D-model of rat $\alpha_5$ integrin was overlayed with AlphaFold2 models of human $\alpha_2$ and $\alpha_6$ integrins. For both human $\alpha_2$ and $\alpha_6$ integrins, predictive N-glycosylations sites were found to be less/not conserved and spatially distant/not present to key $\alpha_5$-N-356 ($\alpha_2$-N-475, $\alpha_6$-N-223), $\alpha_5$-N-773 ($\alpha_2$-absent, $\alpha_6$-N-770 (conserved)) and $\alpha_5$-N-918 ($\alpha_2$-N-1074, $\alpha_6$-N-930). These less conserved sites within $\alpha_2$ and $\alpha_6$ integrins in comparison to $\alpha_3$ and $\alpha_V$ (Fig. 7F) may consistently correlate with their low abundancy in the retrograde and Gal3 interactomes, while these correlations fitted well with $\alpha_3$ and $\alpha_v$ integrins.

### TABLE

Table S1: Cryo-EM data collection, refinement and validation statistics.

| | Neuraminidase-treated rat α5β1 integrin | Neuraminidase-treated rat α5β1 integrin - Gal3 complex |
|---|---|---|
| **Data collection and processing** | | |
| Magnification | 96,000 | 81,000 |
| Voltage (kV) | 300 | 300 |
| Camera | TFS Falcon 3 | Gatan K3 |
| Electron exposure (e–/Å$^2$) | 42 | 61 |
| Defocus range (μm) | -0.8 – 2.0 | -1.0 – 2.8 |
| Pixel size (Å) | 0.832 | 1.06 (0.53 super resolution) |
| Micrographs used | 2884 | 15,839 |
| Total extracted particle images | 1,444,502 | 4,624,708 |
| Refined particle images | 470,092 | 846,399 |
| Final particle images | 101,740 (head)<br>277,162 (leg) | 111,868 |
| Map resolution, head (Å) | 3.7 (head); 4.7 (leg) | 6.9 |
| FSC threshold | 0.143 | 0.143 |
| Map resolution range (Å) | 3.2 – 8.0 (head)<br>4.0 - 10.0 (leg) | 5.4 – 11.8 |
| **Refinement** | **head:** α5, 94-691, β1, 25-504 | |
| Refinement package | Phenix dev 4778 | |
| Model resolution (Å) | 3.8 | |
| FSC threshold | 0.5 | |
| Map sharpening *B* factor (Å$^2$) | -144.0 | |
| Model composition | | |
| Non-hydrogen atoms | 8324 | |
| Protein residues | 1078 | |
| Ligands | 8 | |
| *B* factors (Å$^2$) | | |
| Protein | 85.55 | |
| Ligand | 78.53 | |
| R.m.s. deviations | | |
| Bond lengths (Å) | 0.004 | |
| Bond angles (°) | 0.799 | |
| Validation | | |
| MolProbity score | 2.18 | |
| Clashscore | 15.56 | |
| EMRinger score | 1.92 | |
| Poor rotamers (%) | 1.12 | |
| Ramachandran plot | | |
| Favored (%) | 93.02 | |
| Allowed (%) | 6.98 | |
| Disallowed (%) | 0.00 | |

### MOVIES

**Movie S1. 3D movie of the dynamic co-tracking of mAb13/Gal3 monitored by lattice light sheet microscopy, related to Figure 2D**. RPE-1 cells were sequentially incubated with 200 nM Gal3-Cy3 for 2 min at 4 °C, and then with 5 µg/ml of mAb13-ATTO488 at 4 °C. Coverslips were then transferred into the imaging chamber. Full 3D volume of 60 planes per cell acquired within 2.77 seconds. Sphere color indicates Gal3 co-localization: Green (Gal3-) and yellow (Gal3+). mAb13/Gal3 co-tracking paths are shown in yellow.

**Movie S2. 3D movie of the dynamic co-tracking of 9EG7/Gal3 monitored by lattice light sheet microscopy, related to Figure 2D**. RPE-1 cells were sequentially incubated with 200 nM Gal3-Cy3 for 2 min at 4 °C, and then with 5 µg/ml of 9EG7-ATTO488 at 4 °C. Coverslips were then transferred into the imaging chamber. Full 3D volume of 60 planes per cell acquired within 2.77 seconds. Sphere color indicates Gal3 co-localization: Green (Gal3-) and yellow (Gal3+). 9EG7/Gal3 co-tracking paths are shown in yellow.

**Movie S3. 3D movie of mAb13 antibody endocytosis as monitored by lattice light sheet microscopy, related to Figure S3I, top**. RPE-1 cells stably expressing AP2-GFP (RPE-1-AP2-TagGFP2, green) were incubated for 2 min at room temperature with 5 µg/mL mAb13-Cy3 antibody and transferred into the imaging chamber for LLSM imaging at 27 °C. Full 3D volume of 60 planes per RPE-1-AP2-TagGFP2 cell acquired within 1.56 sec. Sphere color indicates AP2 co-localization: red (AP2-) and yellow (AP2+). Track paths are represented in white color.

**Movie S4. 3D movie of 9EG7 antibody endocytosis as monitored by lattice light sheet microscopy, related to Figure S3I, bottom**. RPE-1 cells stably expressing AP2-GFP (RPE-1-AP2-TagGFP2, green) were incubated for 2 min at room temperature with 5 µg/mL 9EG7-Cy3 antibody and transferred into the imaging chamber for LLSM imaging at 27 °C. Full 3D volume of 60 planes per RPE-1-AP2-TagGFP2 cell acquired within 1.56 sec. Sphere color indicates AP2 co-localization: Red (AP2-) and yellow (AP2+). Track paths are represented in white color.

**Movie S5. 2D movie of time-resolved mAb13/AP2-positive and mAb13/AP2-negative endocytosis imaged by lattice light sheet microscopy, related to Figure S3L, top**. Endocytosis experiments performed as in Movie S3, monitoring characteristic dynamic events from mAb13 antibody signals that either show an overlap with AP2 (AP2+, right inset), or not (AP2-, left inset). Full 3D volume of 60 planes per RPE-1-AP2-TagGFP2 cell acquired within 1.56 sec. This 2D movie shows a side view of a cell slice perpendicular to the detection objective.

**Movie S6. 2D movie of time-resolved 9EG7/AP2-positive and 9EG7/AP2-negative endocytosis imaged by lattice light sheet microscopy, related to Figure S3L, bottom**. Endocytosis experiments performed as in Movie S4, monitoring characteristic dynamic events from 9EG7 signals that either show an overlap with AP2 (AP2+, bottom inset), or not (AP2-, top inset). Full 3D volume of 60 planes per RPE-1-AP2-TagGFP2 cell acquired within 1.56 sec. This 2D movie shows a side view of a cell slice perpendicular to the detection objective.
